# Activating silent glycolysis bypasses in *Escherichia coli*

**DOI:** 10.1101/2021.11.18.468982

**Authors:** Camillo Iacometti, Katharina Marx, Maria Hönick, Viktoria Biletskaia, Helena Schulz-Mirbach, Ari Satanowski, Beau Dronsella, Valérie A. Delmas, Anne Berger, Ivan Dubois, Madeleine Bouzon, Volker Döring, Elad Noor, Arren Bar-Even, Steffen N. Lindner

**Affiliations:** Max Planck Institute of Molecular Plant Physiology, Am Mühlenberg 1, 14476 Potsdam-Golm, Germany; Génomique Métabolique, Genoscope, Institut François Jacob, CEA, CNRS, Univ Evry, Université Paris-Saclay, 91057 Evry-Courcouronne, France; Institute of Molecular Systems Biology, ETH Zürich, Otto-Stern-Weg 3, 8093 Zürich, Switzerland; Department of Plant and Environmental Sciences, Weizmann Institute of Science, Rehovot, Israel; Department of Biochemistry, Charité Universitätsmedizin, Virchowweg 6, 10117 Berlin, Germany

**Author notes:** equal contributors (Camillo Iacometti, Katharina Marx).

**Keywords:** glycolysis, metabolism, serine, methylglyoxal, synthetic biology

## Abstract

All living organisms share similar reactions within their central metabolism to provide precursors for all essential building blocks and reducing power. To identify whether alternative metabolic routes of glycolysis can operate in *E. coli*, we complementarily employed *in silico* design, rational engineering, and adaptive laboratory evolution. First, we used a genome-scale model and identified two potential pathways within the metabolic network of this organism replacing canonical Embden-Meyerhof-Parnas (EMP) glycolysis to convert phosphosugars into organic acids. One of these glycolytic routes proceeds via methylglyoxal, the other via serine biosynthesis and degradation. Then, we implemented both pathways in *E. coli* strains harboring defective EMP glycolysis. Surprisingly, the pathway via methylglyoxal immediately operated in a triosephosphate isomerase deletion strain cultivated on glycerol. By contrast, in a phosphoglycerate kinase deletion strain, the overexpression of methylglyoxal synthase was necessary for implementing a functional methylglyoxal pathway. Furthermore, we engineered the ‘serine shunt’ which converts 3-phosphoglycerate via serine biosynthesis and degradation to pyruvate, bypassing an enolase deletion. Finally, to explore which of these alternatives would emerge by natural selection we performed an adaptive laboratory evolution study using an enolase deletion strain. The evolved mutants were shown to use the serine shunt. Our study reveals the flexible repurposing of metabolic pathways to create new metabolite links and rewire central metabolism.

## Introduction

Excluding some marked exceptions (e.g., (1-3)), the central carbon metabolism of all living organisms can be divided into few interacting pathways, i.e. glycolysis, the pentose phosphate pathway, and the tricarboxylic acid (TCA) cycle, each representing highly conserved reaction patterns. The emergence of these conserved patterns can be explained in two complementary ways. First, the structure of central metabolism mostly reflects a metabolic network that could have emerged under primordial conditions where primitive systems prevailed solely by relying on abiotic catalysis (4, 5). This primordial network could have served as the backbone supporting the emergence of the first cells and, as such, was ‘frozen’ at the origin of life, offering only little flexibility for adaptation throughout the specific evolutionary trajectory of current organisms. Alternatively, and complementarily, central metabolism might constitute an optimal solution for interconverting essential cellular metabolites under a given set of biochemical constraints, including pathway length, favorable thermodynamics, proper kinetics, chemical properties of metabolic intermediates, avoidance of radical enzymes and toxic intermediates (6-10). In this scenario it might be that many pathway solutions have existed which were outcompeted by the ones providing optimal ATP yield (Embden-Meyerhof-Parnas (EMP)) or highest rates (Entner-Doudoroff (ED)).

The universal structure of central metabolism facilitates the study of cellular physiology by enabling to apply knowledge gained from one organism to the understanding of another. On the other hand, this conserved metabolism also severely restricts the metabolic and chemical space we can easily explore and impacts bioproduction by limiting the set of key starting metabolites. Therefore, there is a growing interest in restructuring central metabolism for exploring alternative metabolic networks and pave the way to new bioproduction capabilities, e.g. by the replacing EMP glycolysis with the stoichiometric favorable none oxidative glycolysis (11-14). Such novel routes can recruit enzymes from heterologous sources or even integrate new-to-nature metabolic conversions (15). A possible strategy of this sort is to explore how the native enzymes and pathways of an organism for implementing new pathways that can replace segments of central metabolism. There are two main advantages for this approach. From a practical point of view, it relies only on enzymes that were optimized during evolution to operate within the desired cellular environment. From a scientific perspective, the recruitment of enzymes belonging to different cellular processes to establish novel metabolic networks is akin to the emergence of pathways during evolution and thus provides a platform to explore such key evolutionary events (16).

In this study, we aimed to fully replace the endogenous EMP glycolysis in the model bacterium *Escherichia coli* by relying only on native enzymes. An *in-silico* analysis identified multiple possible pathways allowing the required conversion of phosphosugars into pyruvate. We engineered two of these routes in *E. coli* strains lacking key EMP glycolysis enzymes. First, we show that a methylglyoxal-dependent route (17-19) can carry all glycolytic flux to support a high growth rate. Furthermore, we implemented the ‘serine shunt’, which bypasses glycolysis via serine biosynthesis and deamination to pyruvate. We demonstrate that the operation of this synthetic route relies on a delicate balance between the rate of serine production and consumption, avoiding the inhibitory accumulation of this amino acid. In parallel, we submitted an *E. coli* strain lacking EMP glycolysis to adaptive evolution, which selected for the emergence of the serine shunt after ∼140 days of cultivation.

## Results

### *In silico* analysis of potential pathways bypassing glycolysis in *E. coli*

We aimed to uncover latent metabolic routes that can potentially replace the canonical EMP glycolysis and offer new connections between phosphosugar and organic acid metabolism. We used the latest metabolic model of *E. coli* from the BiGG database (20) and systematically searched for all thermodynamically feasible combinations of native reactions that can convert the feedstock glycerol into pyruvate, an organic acid from which all metabolites of the TCA cycle can be derived (Methods). We chose glycerol as the feedstock as it can be directly converted into the simplest phosphosugars – the triose phosphates dihydroxyacetone phosphate (DHAP) and glyceraldehyde 3-phosphate (GAP) – such that its conversion to pyruvate is expected to follow a 1:1 stoichiometry. Using this approach, we were able to identify more than 100 thermodynamically feasible routes that can potentially convert glycerol to pyruvate (Supplementary Figure S1). These routes and their variants, e.g. routes that share the same intermediates and general metabolic conversions but use different cofactors (e.g., quinones instead of NAD^+^), represent combinations of four broad strategies to convert glycerol to pyruvate (Fig. 1): (i) oxidation of triose phosphates to phosphoglycerate, followed by conversion to phosphoenolpyruvate and pyruvate, as within canonical EMP glycolysis; (ii) generation of pyruvate from phosphosugars via the Entner Doudoroff (ED) pathway (blue arrows in Fig. 1); (iii) conversion of triose phosphates to methylglyoxal, which is then oxidized to pyruvate (magenta arrows in Fig. 1); (iv) diversion of 3-phosphoglycerate (3PG) towards serine biosynthesis, followed by serine deamination to pyruvate (green arrows in Fig. 1). Since the ED pathway has been analyzed and compared to EMP glycolysis in multiple previous studies (7, 21-23), we decided to focus on the methylglyoxal-dependent and serine-dependent routes and test their potential of bypassing the EMP in *E. coli*. Notably, our approach differs from a previous study which used a similar computational method but included all KEGG database reported reactions (not only from *E. coli*), limited their search to pathways yielding at least 1 ATP (10), thus would not find the routes, identified by our approach, via serine or methylglyoxal which are ATP-neutral or even waste ATP, respectively.

**Figure 1:**
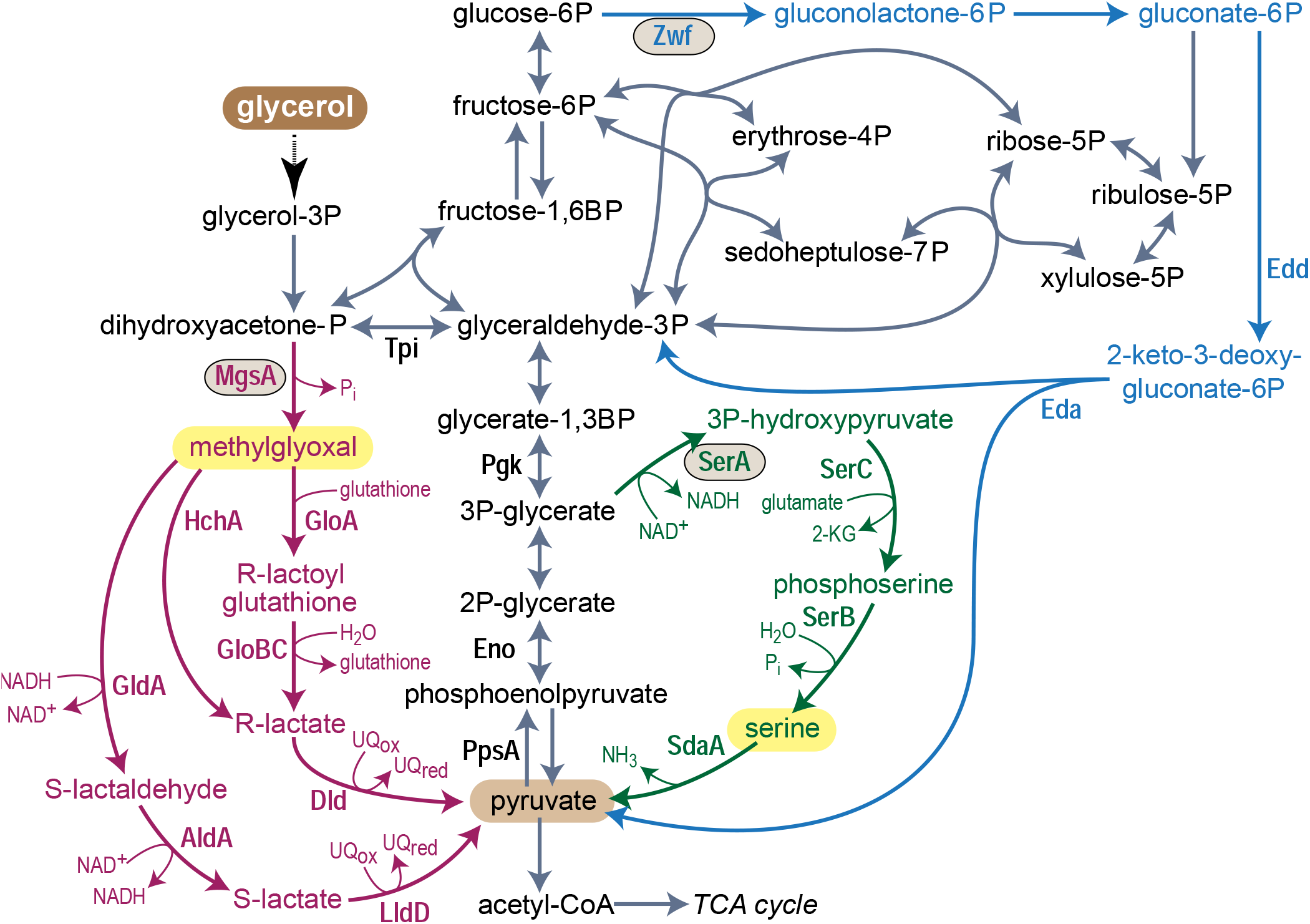
Overview of central metabolism of *E. coli*. Embden-Meyerhof-Parnas (EMP) glycolysis is shown by gray arrows, Entner-Doudoroff pathway is shown by blue arrows, the methylglyoxal bypass is shown in pink, and the serine shunt is shown in green. Key initial committing enzymes of the pathways are circled and namesake intermediates are highlighted in yellow, abbreviated as follows: Entner-Doudoroff pathway: Zwf, glucose 6-phosphate dehydrogenase; Edd, 6-phosphogluconate dehydratase; Eda, 2-keto-3-deoxygluconate 6-phosphate aldolase. Methylglyoxal pathway: MgsA, methylglyoxal synthase; GldA, glycerol dehydrogenase; AldA, aldehyde dehydrogenase; LldD, L-lactate dehydrogenase; HchA, D-lactate dehydratase; GloA, glyoxalase; GloCB, hydroxyacylglutathione hydrolase; Dld, D-lactate dehydrogenase. Serine shunt: SerA, 3-phosphoglycerate dehydrogenase; SerC, phosphoserine aminotransferase; SerB, phosphoserine phosphatase; SdaA, serine deaminase. Glycolysis: Pgk, 3-phosphoglycerate kinase; Eno, enolase; PpsA, PEP synthase.

### A glycolytic pathway via methylglyoxal

*E. coli* is known to channel flux towards methylglyoxal upon accumulation of triose phosphates which results from phosphate depletion limiting the activity of GAP dehydrogenase or from excessive carbon intake (17, 24). The operation of the methylglyoxal bypass thus enables the cells to adapt to imbalanced central metabolism (25). Moreover, previous studies demonstrated that strains deleted in the gene coding for triose phosphate isomerase (Δ*tpi*) grown on glucose led to activation of the methylglyoxal bypass, splitting carbohydrate metabolism between EMP glycolysis from GAP and methylglyoxal metabolism from DHAP (18, 19, 26). However, growth of the *tpi* deletion strain on glucose still partially relies on a functional EMP glycolysis, converting GAP into pyruvate. To our knowledge, rerouting the entire glycolytic flux towards methylglyoxal metabolism, without any pyruvate generated via EMP glycolysis, has not been demonstrated before. Considering the high toxicity of methylglyoxal (27, 28), it was not clear whether the methylglyoxal bypass could indeed support all glycolytic flux without resulting in severe growth defects.

To assess the capability of methylglyoxal metabolism to replace glycolysis we first constructed a Δ*tpi* strain. This strain, when fed with glycerol as sole carbon source requires the activity of the methylglyoxal shunt to support almost the entire carbon assimilation flux (different to when using glucose as feedstock). To our surprise, when cultivated on glycerol as sole carbon source, the Δ*tpi* strain immediately grew without the need for any dedicated overexpression of methylglyoxal synthase MgsA or evolution (yellow line in Fig. 2A). As previously described, a Δ*tpi* strain utilizes glucose derived GAP via EMP and DHAP via the methylglyoxal route (26). On glycerol the *tpi* deletion can theoretically be bypassed by the ED pathway. Here, a GAP is “borrowed” to condense with a DHAP to make fructose 1,6-bisphosphate via fructose 1,6-bisphosphate aldolase (Fba) and then use the ED pathway to generate GAP and pyruvate, which both can recover the GAP condensed with DHAP by Fba. In order to exclude this theoretical option, we deleted the genes *zwf* and *eda* individually in the Δ*tpi* strain to block the ED pathway. The resulting strains Δ*tpi* Δ*zwf* and Δ*tpi* Δ*eda* grew similar to the Δ*tpi* strain, ensuring no contribution of the ED pathway to the metabolic bypass in the *tpi* deletion strain. An even stronger confirmation of the activity of the methylglyoxal pathway is provided by the fact that the deletion of *mgsA* in the Δ*tpi* strain abolished growth (Fig. 2B).

**Figure 2:**
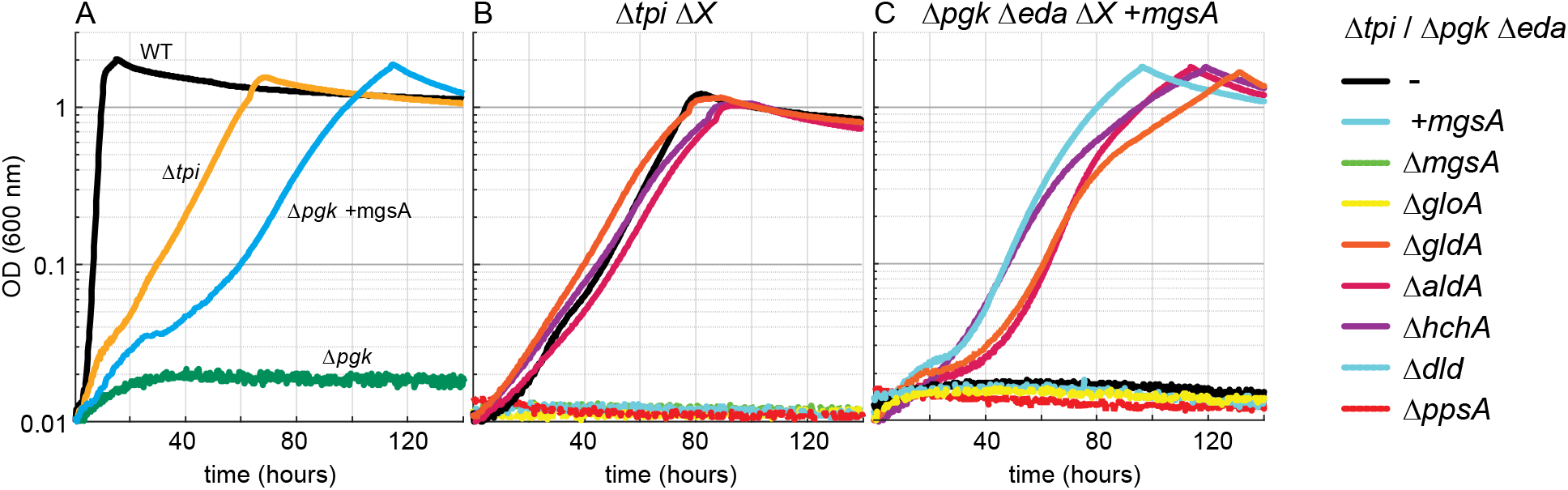
The methylglyoxal pathway bypasses EMP glycolysis in the Δ*tpi* strain and Δ*pgk* Δ*eda* strain overexpressing *mgsA*. Growth on 20 mM glycerol of Δ*tpi* and Δ*pgk* Δ*eda +mgsA* strains (A). Growth on 20 mM glycerol of Δ*tpi* strain (B) and Δ*pgk* Δ*eda* +*mgsA* strain (C) harboring an additional deletion of one of the enzymes potentially involved in methylglyoxal degradation. Graphs represent triplicate repeats, showing similar growth (± < 5%).

We then constructed a strain lacking EMP glycolysis by deleting phosphoglycerate kinase (Δ*pgk*). As expected the Δ*pgk* strain was only able to grow, if two carbon sources were provided, feeding both parts of the metabolism (divided by the *pgk* deletion), e.g. glycerol and succinate. As the ED pathway could theoretically bypass the *pgk* deletion we additionally blocked it by deleting the 2-keto-3-deoxygluconate 6-phosphate aldolase gene (*eda*), resulting in strain Δ*pgk* Δ*eda*. In contrast to the Δ*tpi* strain, the Δ*pgk* Δ*eda* strain required overexpression of the gene coding for methylglyoxal synthase (*mgsA*) from a plasmid to grow on glycerol as sole carbon source (green *vs*. blue line in Fig. 2A).

Our results indicate that the methylglyoxal bypass can indeed completely replace glycolysis, despite the high reactivity of its namesake intermediate. Notably, when glycerol is the carbon source, DHAP is expected to accumulate to a high concentration in the Δ*tpi* strain, where its metabolism is completely blocked, while in the Δ*pgk* Δ*eda* strain such accumulation can be prevented by DHAP conversion to other phosphosugars (Fig. 1). As it has been reported that accumulation of DHAP leads to the expression of *mgsA* and activation of the methylglyoxal bypass in *E. coli* (24) we compared the transcription levels of *mgsA* between the WT, Δ*tpi* and Δ*pgk* Δ*eda* strains in qPCR experiments on cells grown on M9 minimal medium containing glycerol (WT and Δ*tpi*) or glycerol and succinate (WT, Δ*tpi*, Δ*pgk* Δ*eda*). When grown on glycerol, *mgsA* transcript levels in Δ*tpi* was lower compared to the transcript levels in the WT. Similarly, when glycerol and succinate were the carbon sources, *mgsA* transcript level was lower in the Δ*tpi* strain but similar in Δ*pgk* Δ*eda* in comparison to the levels determined for the WT (supplementary Figure S2). These results suggest that growth of the Δ*tpi* strain fed on glycerol is enabled by an elevated cellular DHAP concentration which enforces a high carbon flux through the methylglyoxal pathway without requiring any change of expression of *mgsA*.

In the case the Δ*tpi* strain is fed on glycerol, pyruvate as the final product of the methylglyoxal pathway should aliment all parts of central carbon metabolism through gluconeogenesis, TCA cycle and anaplerotic reactions, whereas in the case of the Δ*pgk* Δ*eda* strain overexpressing *mgsA*, pyruvate should be essential for the operation of TCA, anaplerotic reactions and for providing 3-phosphoglycerate as precursor of serine and glycine biosynthesis. Thus, in both strains a deletion of phosphoenolpyruvate synthase (*ppsA)*, which is essential for to provide PEP from pyruvate for e.g. anaplerotic reactions, should impede methylglyoxal-dependent growth (Fig. 1). Indeed, the Δ*tpi* Δ*ppsA* strain, and the Δ*pgk* Δ*eda* Δ*ppsA* (overexpressing *mgsA*) strain lost the ability to grow on glycerol (Fig. 2B,C), thus confirming the pivotal activity of the methylglyoxal bypass.

*E. coli* can potentially convert methylglyoxal to pyruvate using three different routes (Fig. 1, magenta arrows): (i) methylglyoxal attachment to glutathione, followed by hydrolysis to D-lactate and oxidation to pyruvate; (ii) direct electron rearrangement of methylglyoxal to give D-lactate, which is oxidized to pyruvate; (iii) reduction of methylglyoxal to lactaldehyde, followed by oxidation to D-lactate and pyruvate. Previous studies suggest that the dominant route is the glutathione-dependent methylglyoxal detoxification pathway (18, 25, 29, 30). However, an unequivocal proof that this pathway channels the entire methylglyoxal metabolism flux and cannot be replaced by the two other routes is still lacking. To finally settle this question, we used the Δ*tpi* and Δ*pgk* Δ*eda* strains to generate multiple deletion strains, each carrying the deletion of a different enzyme of the methylglyoxal catabolism. We found that the absence of HchA, GldA, or AldA did not impair growth on glycerol via the methylglyoxal bypass (Fig. 2B), indicating that the two glutathione-independent routes described above contribute very little, if at all, to methylglyoxal metabolism. On the other hand, the absence of GloA or Dld completely abolished growth on glycerol (Fig. 2B). These findings confirm the glutathione-dependent route to be indispensable and sufficient for methylglyoxal metabolism. While in many cases ^13^C-labeling experiments are valuable to verify a pathway’ s activity, in the case of the methylglyoxal route no difference in the labeling pattern compared to EMP glycolysis was expected, thus ^13^C-labeling experiments were omitted.

### The serine shunt can replace EMP glycolysis

The second glycolytic bypass identified by our *in-silico* analysis is the serine shunt which combines biosynthesis and degradation of serine (green arrows in Fig. 1; supplementary Fig S1). To assess the feasibility of this pathway, we generated an enolase deletion strain (Δ*eno*), which cannot operate EMP glycolysis, while still being able to generate 3-phosphoglycerate from which the serine biosynthesis pathway begins (Fig. 1). For growth on a minimal medium the Δ*eno* strain requires two carbon sources for feeding both upper and lower parts of central carbon metabolism (e.g. glycerol and succinate or xylose and succinate).

We assumed that overexpression of the four enzymes involved in serine biosynthesis from 3-phosphoglycerate and degradation to pyruvate would enable the activity of the serine shunt and support growth of the Δ*eno* strain on glycerol only (Fig. 1, green arrows). Therefore we constructed a plasmid constitutively expressing a synthetic operon containing the four following genes: (i) a variant of the native gene coding for 3-phosphoglycerate dehydrogenase which was engineered to remove its native allosteric inhibition by serine (*serA\**, coding for SerA H344A N346A N364A, catalyzing the first step in serine biosynthesis) (31); (ii) *serC*, coding for phosphoserine aminotransferase; (iii) *serB*, coding for phosphoserine phosphatase; and (iv) *sdaA*, coding for the major serine deaminase isozyme in *E. coli*, converting serine into pyruvate (32). However, the resulting plasmid (p-*serA\**-*serB*-*serC*-*sdaA*) failed to support growth of the Δ*eno* strain on glycerol. Moreover, PCR analysis of the transformants revealed that the strains did not carry the whole operon but only fractions of it. We assume that this occurred due to toxic effects of the overexpression of the serine biosynthesis genes and hence resulted in gene inactivation/removal from the plasmid.

Serine’ s toxicity is well known and can be attributed, at least partially, to its deamination to the highly reactive and toxic compound hydroxypyruvate (33). Indeed, we found that addition of even small amounts of serine (1-8 mM) substantially delayed the growth of the WT (Fig. 3A) and the Δ*eno* strain (Fig. 3B) with glycerol and succinate as co-carbon sources. The Δ*eno* strain seems to be more sensitive to serine than the WT strain, as its growth was completely inhibited at serine concentrations of ∼8 mM. Further deletion of the genes coding for serine deaminases (Δ*sdaA* Δ*sdaB* Δ*tdcB* Δ*tdcG* (34)), thus removing a serine sink, increased the sensitivity of both the WT and the Δ*eno* strain further: even small amounts of serine (0.5 mM) completely inhibited growth on glycerol and succinate (Fig. 3C,D). Interestingly, in one of the Δ*eno* strain cultures incubated in the presence of 48.8 mM serine, the cells began to grow after >100 hours (Fig. 3B). This suggests the emergence of mutations enabling the cells to tolerate this high level of serine. We isolated three of these individual serine tolerant strains and sequenced their genomes. In the resulting reads we found that in all three strains a somewhat different genomic region was multiplied several fold (as indicated by a 4-8 fold increased sequencing coverage, Supplementary Figure S3): 1,761,810 bp to 1,816,965 bp, 1,762,676 bp to 1,781,465 bp, or 1,762,962 bp to 1,798,478 bp. Notably, these genomic regions harbor the genes *sufA* and *sufB*, coding for subunits of the iron-sulfur cluster scaffold complex (35). Among its other functions, this cluster serves as an essential component of the primary serine deaminase enzyme SdaA (36). Hence, it seems that increasing the availability of iron-sulfur clusters in the cell contributed to a faster and more efficient degradation of serine, making the cells more tolerant to this amino acid. To see if the increased tolerance to serine would enable the mutated strain to grow on glycerol via the serine shunt we overexpressed the four genes described above. Indeed, we found that transformation of the G3 mutant strains with p-*serA\**-*serB*-*serC*-*sdaA* enabled them to grow on glycerol as sole carbon source (Fig. 4A). It therefore seems that the above-mentioned changes supported a strong metabolic sink for toxic serine to enable high flux via serine biosynthesis and degradation with minimal adverse effects.

**Figure 3:**
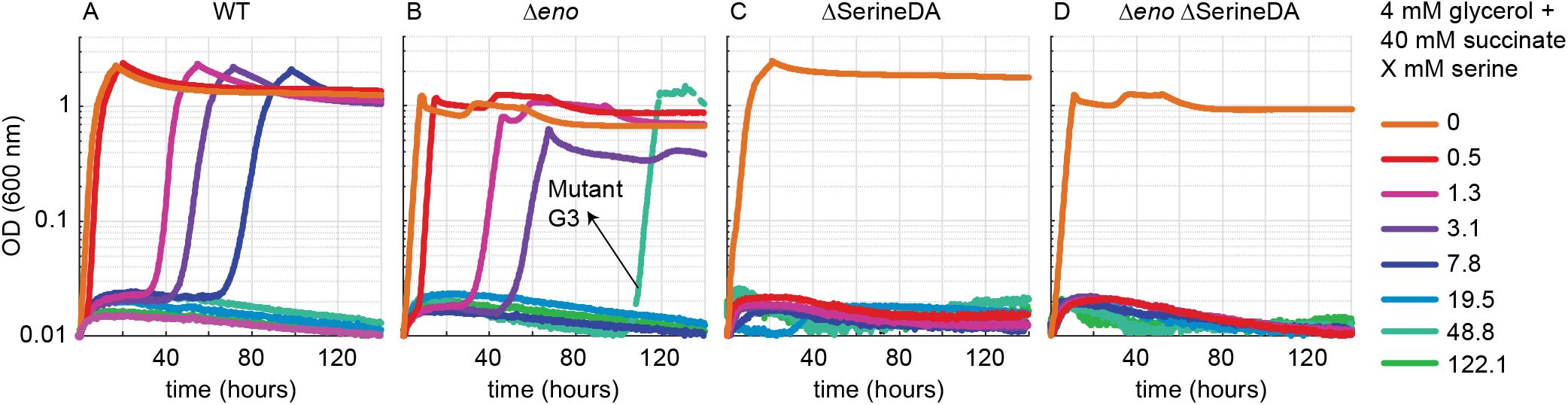
Serine inhibits growth at low millimolar concentrations in WT *E. coli* and abolishes growth at submillimolar concentrations in strains lacking serine deaminase (‘SerineDA’) enzymes. Strain WT (A), Δ*eno* (B), ΔSerineDA (C, Δ*sdaA* Δ*sdaB* Δ*tdcB* Δ*tdcG*), and Δ*eno* ΔSerineDA (D, Δ*eno* Δ*sdaA* Δ*sdaB* Δ*tdcB* Δ*tdcG*) were incubated in media containing 4 mM glycerol and 40 mM succinate. Serine concentrations were added as indicated. An isolate (G3) from the Δ*eno* culture growing in the presence of 48.8 mM serine after an extended lag-phase was obtained and analyzed by genome sequencing. Graphs represent triplicate repeats, showing similar growth (± 5%).

**Figure 4:**
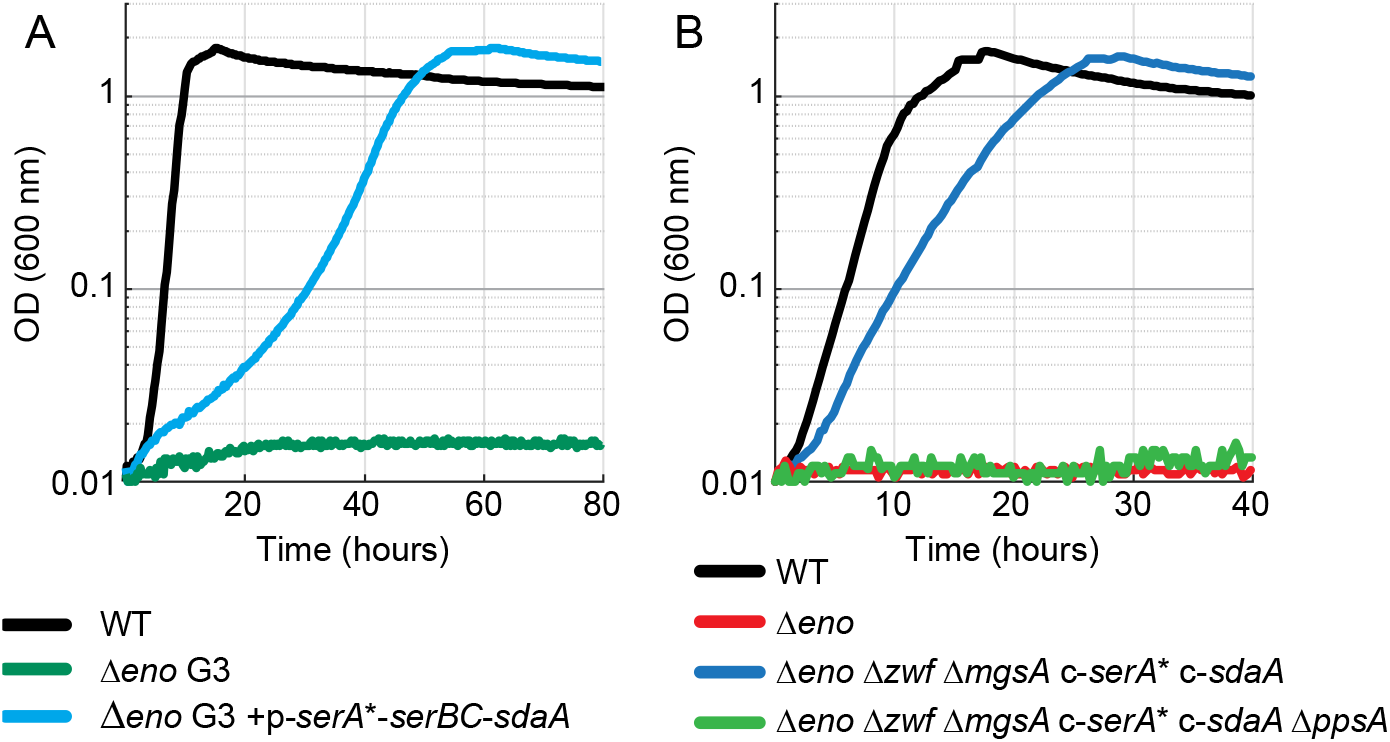
Growth on glycerol of the serine-resistant Δ*eno* isolat G3 transformed with p-*serA\**-*serB*-*serC*-*sdaA* (A) and Δ*eno* strain with chromosome-integrated *serA\** and *sdaA* genes (B, Δ*eno* c-*sdaA* c-*serA*\*). Growth experiments were performed in at least three repeats, showing similar growth behavior (± 5%).

In order to decrease and stabilize the rate of serine biosynthesis to avoid its accumulation, we pursued a strategy of gene expression from the chromosome rather than from a plasmid in the naïve Δ*eno* strain. The genes *serA*\* and *sdaA* were overexpressed from the chromosome (see method section), while the native expression was preserved for the genes *serB* and *serC*. We found that overexpression of *serA\** from the chromosome in the Δ*eno* strain is possible only after the chromosomal overexpression of *sdaA* is established, again pointing to the necessity of a balanced serine synthesis and degradation activity for growth via the serine shunt (data not shown). Finally, to avoid any carbon flux through the methylglyoxal bypass and the ED pathway, we deleted *mgsA* and the gene encoding glucose 6-phosphate dehydrogenase (Δ*zwf*). The Δ*eno* Δ*mgsA* Δ*zwf* strain overexpressing *serA\** and *sdaA* from the chromosome was indeed able to grow on glycerol with no need for prior adaptation (Fig. 4B). As expected, the deletion of *ppsA*, which blocks the conversion of pyruvate to phosphoenolpyruvate, abolished growth (Fig. 4B), thus confirming that this essential metabolite cannot be produced by some variant of EMP glycolysis that bypasses or replaces the enolase reaction. Overall, these results confirm that the serine shunt can be implemented in the cells to replace the canonical EMP glycolysis using a rational engineering approach, where the rate of serine biosynthesis is balanced with that of serine degradation, consequently avoiding deleterious accumulation of this amino acid. Similar to the methylglyoxal rout ^13^C-labeling experiments were not expected to show any difference in the labeling pattern of serine shunt and EMP glycolysis and hence were omitted.

### Continuous culture evolution results in the emergence of the serine shunt

As an alternative option to the rational engineering of a synthetic glycolysis bypass, we resorted to continuous culture protocols to investigate which of the three bypass routes - ED pathway, methylglyoxal bypass, or serine shunt - would emerge naturally during long-term cultivation of the Δ*eno* strain under selective conditions (we chose to start with the Δ*eno* strain as it has the potential to activate all three mentioned bypass routes). We applied a medium-swap regime using GM3 continuous cultivation devices (37, 38) to a growing population of the Δ*eno* strain, fed alternatively with a permissive medium containing both glycerol and succinate, and a stressing medium containing only glycerol (Methods). When the turbidity of the culture, measured in real time, was below a predefined value, the culture was diluted with the permissive medium; otherwise, the stressing medium was used to dilute the culture. Such medium-swap protocol enables gradual adaptation of the bacterial population to the stressing growth conditions (37, 38), in our case proliferation without succinate as substrate of the lower part of the carbon metabolism.

We conducted the adaptive evolution experiment in two parallel cultures, both of which adapted to grow on the stressing medium (containing glycerol but no succinate) after 130-140 days amounting to more than 900 generations (Fig. 5A). We then applied a turbidostat mode – cultivating the cells solely on the stressing medium and diluting the culture every time a pre-defined turbidity is reached – to evolve the culture towards a higher growth rate (Fig. 5A). We isolated two strains from each of the evolved cultures and sequenced their genomes. Strains isolated from each of the evolved cultures showed highly similar mutation profiles (Supplementary Table S2). All sequenced strains harbored a non-synonymous mutation in *serA*, either replacing T372 with asparagine or replacing L370 with methionine. No mutation was observed in genes coding for enzymes participating in the methylglyoxal shunt or the ED pathway. This suggested that the adaptive evolution of the Δ*eno* strain led to the emergence of the serine shunt, rather than to any other of the possible glycolytic bypass routes. We found that the isolates from only one of the evolved cultures could stably grow on glycerol alone when cultivated within a 96-well plate (Fig. 5B). The strain iso1 was further characterized.

**Figure 5:**
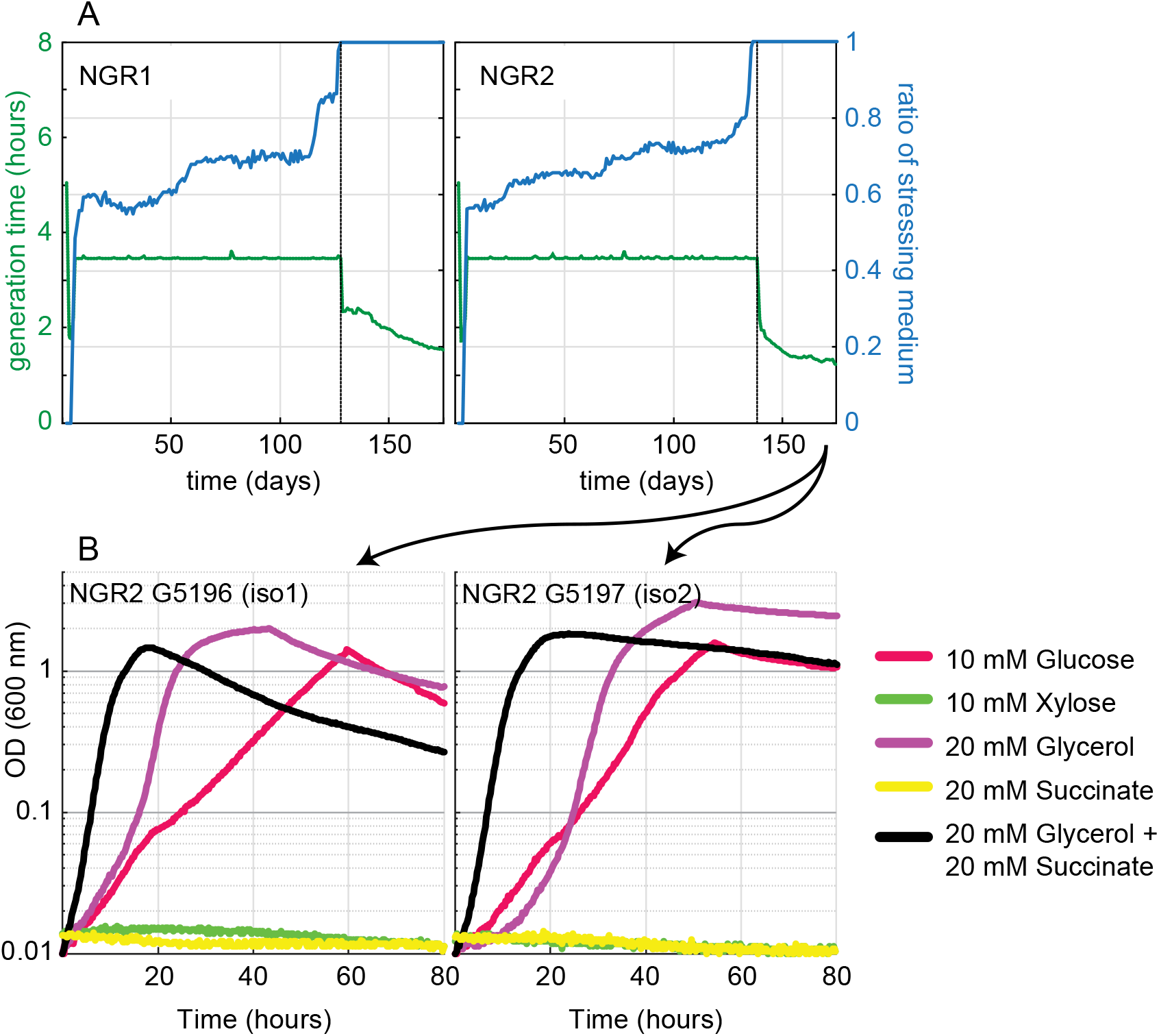
Adaptive evolution in continuous culture of the Δ*eno* strain to growth on glycerol. (A) Two independent cultures were subjected to a medium swap regime in GM3 devices (see Methods section). Blue lines show the ratio of stressing medium over relaxing medium (right axis). The stressing medium contained 20 mM glycerol, relaxing medium 20 mM glycerol plus 10 mM succinate. The generation time of the growing population was set to 3.5 hours. Once steady growth was obtained in the stressing medium, the bacterial population was cultivated under turbidostat regime. Generation times are indicated by the green dashed lines (left axes). (B) Growth analysis of two isolates from NGR2 on the indicated carbon sources. In all cases growth experiments were performed in triplicates, showing similar growth (± 5%).

The four reactions of the serine shunt usually carry much less flux than the amount needed for the serine shunt to operate as sole glycolytic option. To analyze if the expression of the corresponding genes is altered in the iso1 strain, we performed qPCR experiments after growth on glycerol. The transcript level of all serine biosynthesis genes was increased in the iso1 strain compared to the WT reference, showing a 3-, 11- and 2.5-fold increased RNA level of *serA, serC* and *serB* respectively. Surprisingly, *sdaA* expression did not significantly differ from the WT control (supplementary Fig. S4). To ensure that no glycerol is converted via the alternative bypasses we deleted their key reactions in the iso1 strain. Neither deletion of *mgsA* nor *zwf* abolished growth (Fig. 6A,B,C), indicating that the glycolysis bypass in the iso1 strain does not rely on methylglyoxal bypass or the ED pathway. Despite our continuous efforts, we were unable to delete *sdaA*, which suggests that serine deamination is essential for growth or survival also in rich medium (e.g., Medium X, see methods), pointing again to the serine shunt as the bypassing route.

**Figure 6:**
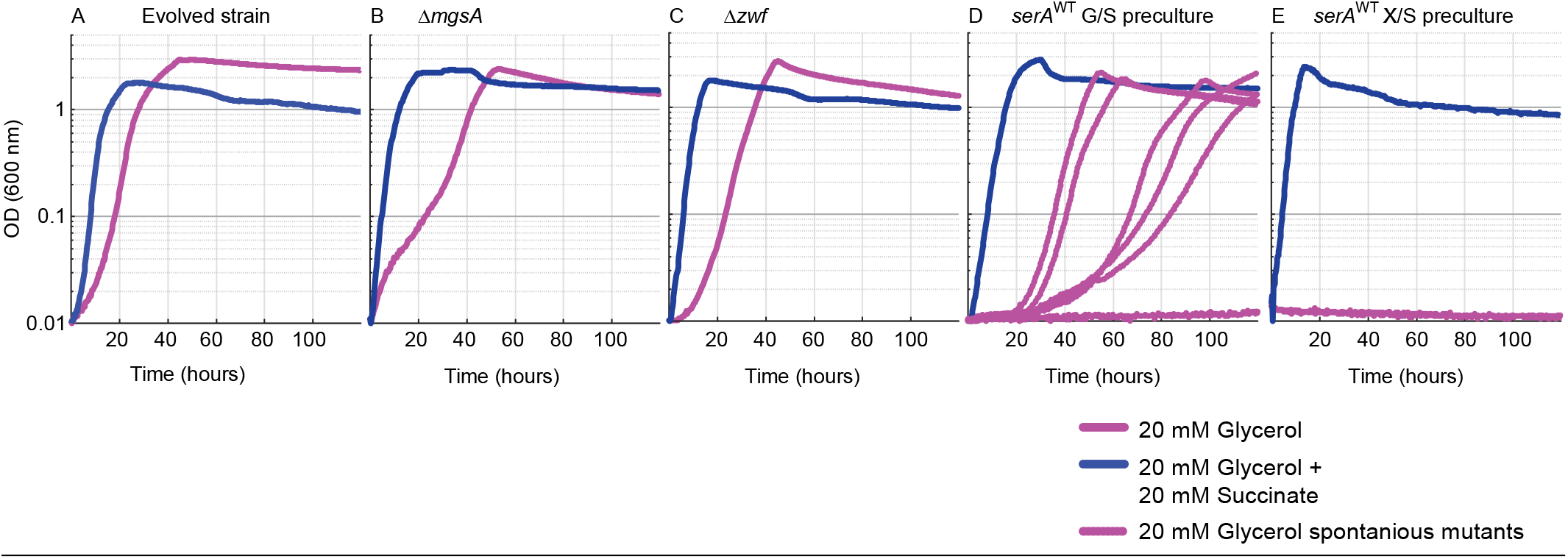
Growth of an evolved Δ*eno* strain (iso1) and derivatives deleted in key enzymes of glycolysis bypassing routes demonstrate the activity of the serine shunt. Growth of the iso1 strain (A), with an additional deletion in *mgsA* (B), *zwf* (C), or reversion of the mutated *serA* (L370M) into the wildtype version (D,E). Cultures of *serA*^WT^ were inoculated from precultures growing in glycerol + succinate (D) or xylose + succinate (E). Growth was recorded with glycerol (pink line) and glycerol + succinate (blue line). Growth of at least 2 independent biological replicates (8 in the case of *serA*^WT^ reversion), were analyzed in three technical repeats, showing similar growth behavior (± 5%).

To further analyze the functioning of the serine shunt in the evolved strain, we restored the WT version of the gene *serA* in strain iso1 (Methods) and performed growth tests on glycerol. The SerA WT derivatives of iso1 had lost the ability to grow on glycerol, but the growth phenotype appeared to be very leaky.

Interestingly, we observed contrasted growth behaviors between various experimental replicates (Fig. 6D): some were unable to use glycerol as sole carbon source, others started to grow slowly after a short lag-phase, while a few grew faster after a similar delay. These observations strongly indicated the emergence of mutations. Indeed, Sanger sequencing of PCR-products of the *serA* locus of every isolate from a growing culture revealed that SerA was mutated. Each of the tested strains contained one of the following mutations: A367T, H342Y, L332P, or R338C, while none of them harbored the SerA variants L370M or T372N identified in isolates from the long-term evolution experiments. Since these mutants arose rather quickly, within hours or days, we concluded that they already appeared in the precultures, which contained glycerol and succinate. Replacing glycerol with xylose for alimenting the upper part of central metabolism in the preculture prevented the emergence of mutants growing on glycerol as sole carbon source within the duration of the experiment (> 5 days) (Fig. 6E).

### Mutated SerA variants lost feedback inhibition by serine

The observations described above point to the pivotal role of SerA reaction for the activity of the serine biosynthesis and degradation pathway as glycolytic bypass: e.g. inability to delete the serine deaminase gene *sdaA* in the evolved strains, mutations fixed in *serA* during the evolution experiment, and spontaneous mutations arising in *serA* in the evolved genetic context of iso1 strain where the WT allele of *serA* had been restored. Interestingly, all the mutations identified in SerA in the course of our study are located in the C-terminal allosteric regulatory domain of the enzyme and thus could possibly modulate the negative allosteric feedback of L-serine on SerA activity. To test whether this hypothesis was correct, we characterized both the kinetic parameters and the regulatory effect of L-serine on enzyme activity of the purified SerA WT, the SerA variants which emerged during the evolution experiments (SerA T372N, culture NGR1) and (SerA L370M, culture NGR2), as well as the triple SerA variant (H344A N346A N364A), which was used for the rational engineering of the serine shunt and was previously reported to be feedback-resistant (31). The promiscuous 2-ketoglutarate reductase activity of SerA was monitored, which in contrast to the oxidation of 3-phosphoglycerate, is a thermodynamically favorable reaction and is known to be also regulated by L-serine (39). All the variants tested exhibited 2-ketoglutarate reductase activities comparable with that of the WT enzyme (Table 1), which is in accordance with the fact that none of the mutations were close to the catalytic site of the enzyme (31). As expected, the activity of WT SerA was strongly decreased in the presence of micromolar concentrations of L-serine (0.5 -10 µM) (Table 1). This concentration range corresponds to the previously published IC_50_ for L-serine, which was determined to be between 2-10 μM (40). By contrast, the SerA variants showed a significantly lower sensitivity to the presence of L-serine, as demonstrated by the minimal decrease of the activity of L370M, T372N and H344A N346A N364A SerA variants with increasing L-serine concentration (Table 1). Our findings in both, the isolated mutants as well as in the rationally engineered strains, support, that the operation of the serine shunt is highly dependent on the presence of a feedback-resistant SerA variant. Kinetic properties of the downstream serine deaminase SdaA (*K*_*M*_ of ∼2.7 mM for L-serine) support this conclusion. The *K*_*M*_ is three orders of magnitude higher than the IC_50_ of SerA for L-serine (36), thus a feedback-inhibited SerA variant might not allow sufficient flux via the pathway to achieve L-serine concentrations high enough for SdaA activity.

**Table 1:** Apparent steady state kinetic parameters of SerA WT and variants and inhibition by serine.

| Enzyme | $K_M^a$<br>(mM) | $k_{cat}^a$<br>($s^{-1}$) | $k_{cat}/K_M$<br>($M^{-1} s^{-1}$) | Inhibition by serine (μM)<sup>b</sup> | | | | | |
| --- | --- | --- | --- | --- | --- | --- | --- | --- | --- |
|  |  |  |  | 0 | 0.5 | 1 | 2 | 5 | 10 |
| SerA WT | 0.033 | 0.346 | $1.04 \times 10^4$ | 100 | 104 | 63 | 40 | 17 | 7 |
| SerA H344A N346A N364A | 0.007 | 0.483 | $6.80 \times 10^4$ | 100 | 92 | 89 | 89 | 96 | 94 |
| SerA T372N (NGR1) | 0.012 | 0.364 | $2.93 \times 10^4$ | 100 | 89 | 83 | 81 | 79 | 82 |
| SerA L370M (NGR2) | 0.019 | 0.359 | $1.88 \times 10^4$ | 100 | 97 | 99 | 107 | 94 | 54 |

<sup>a</sup> Apparent steady state kinetics of SerA variants measured with 2-ketoglutarate
<sup>b</sup> 2-ketoglutarate reductase activity of SerA variants was measured under saturation conditions in the presence of the indicated μM L-serine concentrations. Data was normalized to 100 % enzyme activity without L-serine addition.

## Discussion

Our computational analysis revealed the presence of multiple feasible glycolytic bypasses in *E. coli*’ s native metabolic network. All of these were combinations of the canonical catabolic EMP and ED pathways and two cryptic routes, which so far had not been described as a significant bypass to EMP glycolysis in any organism (in the case of the serine shunt) or were only described as a sink for excess carbon (i.e., the methylglyoxal route) (18, 19, 26). Here, we were able to engineer these two glycolytic alternatives in *E. coli*, relying exclusively on native enzymes. Previous studies observed the emergence of the methylglyoxal pathway in a Δ*tpi* strain during growth on glucose (41) and the serine shunt was proposed to improve acetyl-CoA precursor supply for the bioproduction of poly(3-hydroxybutyrate) (42). But as of yet, to the best of our knowledge, both pathways have not been described as a complete glycolytic bypass.

In addition to the rational engineering of the pathways, a directed evolution experiment evolving an enolase deletion strain, leaving all bypass options open for natural selection. This experiment resulted in the emergence of the serine shunt. Intuitively, a bypass via the ED pathway, which operates in *E. coli* when growing on gluconate (43), might seem more likely to occur. But, evolving the ED pathway to bypass the glycolytic blockade presumably requires the accumulation of several adaptive mutations susceptible to increase the gluconeogenic flux, to inactivate/downregulate the 6-phosphogluconate dehydrogenase and to enhance phosphogluconate dehydratase (*edd*) and 2-keto-3-deoxygluconate 6-phosphate aldolase (*eda*) activity. In contrast, and in line with the outcome of the evolution experiment, engineering the functional serine shunt required only the balanced overexpression of a feedback-resistant SerA variant and of SdaA. Curiously, we showed that the methylglyoxal pathway needed only the overexpression of MgsA in the Δ*pgk* strain, therefore it was expected that the methylglyoxal pathway could be established most easily as a glycolytic bypass by evolution. One reason for the selection of the serine shunt instead of the methylglyoxal pathway during evolution of the enolase deletion strain might be the higher intracellular concentration of 3-phosphoglycerate (∼4 mM) compared to DHAP (∼0.5 mM) during growth on glycerol (44). The 3-phosphoglycerate concentration, determined for WT *E. coli*, is likely even higher in the Δ*eno* strain, this together with the lower ATP cost of the serine shunt might have triggered the fixation of mutations establishing the serine shunt as Δ*eno* bypass.

As expected, reverting the *serA* mutation which emerged in the evolved Δ*eno* strain, abolished growth on glycerol. However, after a short period of incubation, growth associated with the appearance of various point mutations in the *serA* gene was restored. This indicates that the strain underwent an adaptation processes during the evolution, possibly priming its metabolism for the use of the serine shunt. However, these adaptations do not directly reflect in pathway-associated mutations.

Both of the glycolytic bypasses contain toxic intermediates. In our experiments, only the toxic effects of L-serine caused some stress response in the strain transformed with the plasmid overexpressing of serine biosynthesis and degradation genes. On the other hand, no additional overexpression of methylglyoxal degradation enzymes was necessary to implement the methylglyoxal bypass, indicating the sufficiently fast native glutathione-dependent conversion of the highly reactive methylglyoxal to D-lactate. Besides involving a toxic intermediate, the serine shunt presents a thermodynamic barrier at the level of 3-phosphoglycerate dehydrogenase, which additionally is highly inhibited by low concentrations of L-serine.

Why some glycolytic pathways and their variants emerged and manifested throughout all organisms to convert sugars into biomass building blocks, while other biochemical possibilities either never appeared in nature or were discarded during evolution, is not well understood. Some insights came from previous studies of the connection between sugar catabolism and the organism’ s environment (i.e. anaerobic or aerobic). There is evidence that ATP yield outweighs protein cost of a pathway when operating under anaerobic conditions, hence the EMP pathway is dominantly used under these conditions. Contrarily, obligate aerobic organisms prefer the ED pathway, which ensures a lower protein cost and a higher thermodynamic driving force. Accordingly, facultative anaerobic organisms like *E. coli* possess both pathway options (7). The methylglyoxal pathway and serine shunt, besides their structural differences, significantly differ in their ATP balance, i. e. consumption of ATP (methylglyoxal pathway) versus no consumption/formation of ATP (serine shunt) per pyruvate generated. This differences in ATP yield render both pathways infeasible in the absence of terminal electron acceptors, e.g. under fermentative conditions. This might provide an explanation for the absence of these pathways in nature. However, the non-phosphorylative ED pathway operates as the route for glucose to pyruvate conversion in some archaea, and hence similar to the serine shunt produces no net ATP yield (45).

In the past decade, important breakthroughs were achieved in the field of synthetic metabolism. The implementation of C1 assimilatory routes for the fixation of CO_2_ (46, 47), formate (48) or formaldehyde (49), enabling biomass production from renewable resources, are examples of audacious attempts to rewire central carbon metabolism. While primarily motivated by the need to provide sustainable options for the industrial production of fuels and chemicals, these studies accelerate the understanding of the underlying principles of metabolism. Other examples like the implementation of phosphoketolase-dependent non-oxidative glycolysis (11, 14) illustrate the pace synthetic biology has adopted to define the limits and barriers which might have shaped the structures of central metabolism at the onset of cellular life. Our approach, combining rational design of non-natural pathways and experimental evolution, is in line with this effort trying to “make sense of the biochemical logic of metabolism” (9), and at the same time opens opportunities for new bioproduction routes of potential industrial relevance.

## Methods

### Computational analysis to identify glycolytic bypasses in *E. coli*

In order to identify possible pathways that can bypass EMP glycolysis resorting to native *E. coli* enzymes only we used a system analysis approach based on the genome-scale metabolic model of this bacterium (50). The approach is described in detail in the supplementary method.

### Reagents and chemicals

Primers were synthesized by Eurofins (Ebersberg, Germany) (Supplementary Table S3). Screening PCRs were performed using DreamTaq polymerase (Thermo Fisher Scientific, Dreieich, Germany). PrimeSTAR GXL DNA Polymerase (Takara) was used for gene cloning and amplification of deletion cassettes

### Media

LB medium (1% NaCl, 0.5% yeast extract, 1% tryptone) was used for molecular biology work and strain maintenance (except Δ*eno* and Δ*pgk* derived strains). When appropriate, kanamycin (25 μg/mL), ampicillin (100 μg/mL), chloramphenicol (30 μg/mL) or streptomycin (100 µg/mL) was used. Medium X (M9 + 5 g/L casamino acids, 40 mM succinate, 4 mM glycerol, modified after (51)) was used for maintenance of glycolysis deletions strains Δ*eno* and Δ*pgk*. Minimal MA medium (31 mM Na_2_HPO_4_, 25 mM KH_2_PO_4_, 18 mM NH_4_Cl, 1 mM MgSO_4,_ 40 µM trisodic nitrilotriacetic acid, 3 μM CaCl_2_, 3 μM FeCl_3_·6H_2_O, 0.3 μM ZnCl_2_, 0.3 μM CuCl_2_·2H_2_O, 0.3 μM CoCl_2_·2H_2_O, 0.3 μM H_3_BO_3_, 1 μM MnCl_2_, 0.3 µM CrCl_3_, 6 H_2_O, 0.3 µM Ni_2_Cl, 6 H_2_O, 0.3 µM Na_2_MoO_4_, 2 H_2_O, 0.3 µM Na_2_SeO_3_, 5 H_2_O) was used for long-term continuous cultures. For growth analysis M9 minimal medium was used (50 mM Na_2_HPO_4_, 20 mM KH_2_PO_4_, 1 mM NaCl, 20 mM NH_4_Cl, 2 mM MgSO_4_ and 100 μM CaCl_2_, 134 μM EDTA, 13 μM FeCl_3_·6H_2_O, 6.2 μM ZnCl_2_, 0.76 μM CuCl_2_·2H_2_O, 0.42 μM CoCl_2_·2H_2_O, 1.62 μM H_3_BO_3_, 0.081 μM MnCl_2_·4H_2_O). The minimal media were supplemented with various carbon sources as indicated in the main text and hereafter.

### Strains and plasmids

*E. coli* strains used in this study were generated from MG1655 derivative strain SIJ488 (52), which was used as wildtype reference (Table 2). The deletions were carried out by λ-Red recombineering using kanamycin resistance cassettes generated via PCR using the FRT-PGK-gb2-neo-FRT (Km) cassette (Gene Bridges, Germany) for deletion of *sdaA, sdaB, tdcB, tdcG* and *mgsA*. For the deletion of *dld, gldA, gloA, aldA. hchA*, and *ppsA*, pKD3 (chloramphenicol) and pKD4 (kanamycin) where used as a template for amplification of deletion cassettes (53). Primer pairs used are indicated in Supplementary Table S3. Cell preparation and transformation for gene deletion was carried out as described (52, 54). The coding sequences of the WT sequences of *serA* and the mutated genes were amplified by PCR using the primer pairs serA-pet-F and serA-pet-R (Supplementary Table S3). The amplified fragments were cloned into a modified pET16b expression vector (Table 2) by using In-Fusion cloning kit (Takara, Shiga, Japan). The sequence of the inserts of the resulting plasmids was verified by Sanger sequencing. For exchanging the native promoter of *sdaA* with a constitutive strong promoter, a chloramphenicol resistance cassette was amplified from pKD3 by using the primer pair SdaA-ProEx-F and CAP-sdaA-R. In a second PCR the promotor sequence (5’ -ACCTATTGACAATTAAAGGCTAAAATGCTATAATTCCAC-3’, (54)) was amplified from pZ-ASS using the primer pair pS-bridge and SdaA-ProEx-R. The purified PCR products were used in a fusion PCR together with the primer pair SdaA-ProEx-F and SdaA-ProEx-R. This resulted in a promoter exchange cassette containing 50 bp flanks to integrate into the intergenic region between *nudL* and *sdaA*. To integrate the feedback-resistant version of 3-phosphoglycerate dehydrogenase (SerA*, H344A N346A N364A) and simultaneously replace the native promotor of the gene with a strong constitutive one, a chloramphenicol resistance cassette was amplified from pKD3 using primer pair serA*-ProEx-F and CAP-SerA*-R. A PCR product containing a constitutive strong promotor (5’ - ACCTATTGACAATTAAAGGCTAAAATGCTATAATTCCAC-3’, (54)) and the *serA*\* gene was amplified using primer pair pS-bridge and serA*-ProEx-R. The purified PCR products were used in a fusion PCR together with the primer pair serA*-ProEx-F and serA*-ProEx-R, resulting in a chloramphenicol cassette containing *serA*\* behind a strong promotor and a 50 bp flank upstream of the CAT cassette to integrate into the intergenic region behind *rpiA*.

**Table 2:**
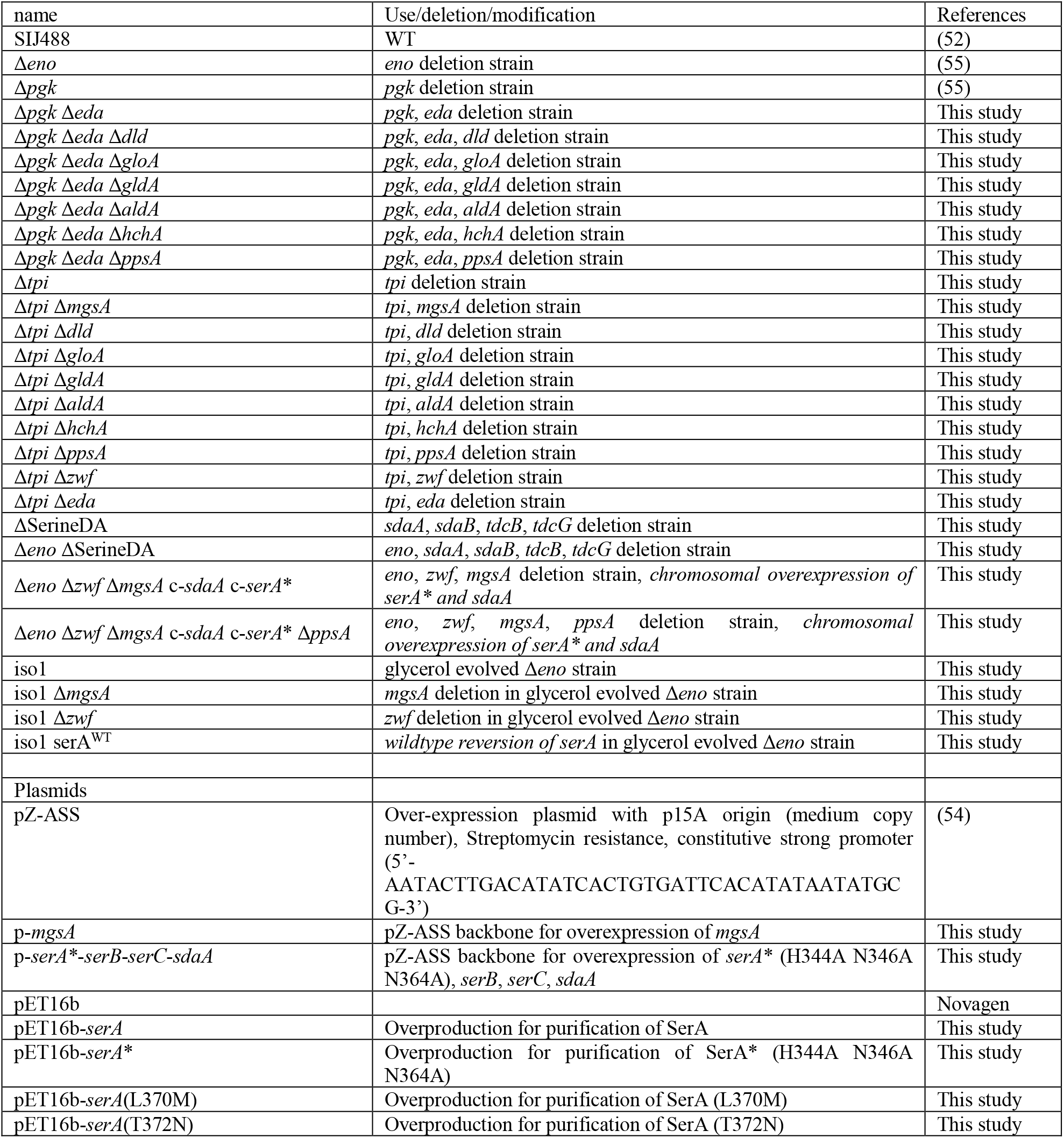
Strains and plasmids used in this study

### Evolution in GM3-driven long-term continuous culture

For evolution experiments pre-cultures of Δ*eno* strain were obtained in permissive minimal MA medium supplemented with 20 mM glycerol and 10 mM succinate. The pre-culture was used to inoculate the growth chambers (16 ml per chamber) of two parallel independent GM3 devices (37). A continuous gas flow of sterile air through the culture vessel ensured constant aeration and growth in suspension by counteracting cell sedimentation. The cultures were grown in the corresponding medium under turbidostat mode (dilution threshold set to 80 % transmittance (OD ≈ 0.4, 37°C) until stable growth of the bacterial population. The cultures were then submitted to a conditional medium swap regime. This regime enabled gradual adaptation of the bacterial populations to grow in a non-permissive or stressing medium which contained 20 mM glycerol only. Dilutions of the growing cultures were triggered every 10 minutes with a fixed volume of medium calculated to impose a generation time of 3.5 hours on the cell population, if not otherwise stated. The growing cultures were fed by permissive or stressing medium depending on the turbidity of the culture with respect to a set OD threshold (OD_600_ value of 0.4). When the OD exceeded the threshold, a pulse of stressing medium was injected; otherwise a pulse of permissive medium was injected. When the cultures grew in 100 % stressing medium the feeding mode was set to turbidostat to increase growth rates. Three isolates were obtained on agar-solidified stressing medium from both evolution experiments and further analyzed.

### Genomic analysis of evolved strains

Pair-end libraries (2×150 bp) were prepared with 1 µg genomic DNA from the evolved strains as well as from the ancestor Δ*eno* strain and sequenced using a MiSeq sequencer (Illumina). The PALOMA pipeline, integrated in the platform Microscope (http://www.genoscope.cns.fr/agc/microscope) was used to map the reads against *E. coli* K12 wildtype strain MG1655 reference sequence (NC_000913.3) for detecting single nucleotide variations, short insertions or deletions (in/dels) as well as read coverage variations (56). For genomic analysis of serine-resistant Δ*eno* mutants, strains were cultured overnight at 37°C in 4 mL medium X (see Methods, Media). Genomic DNA from overnight cultures was extracted using the NucleoSpin Microbial DNA kit (Macherey-Nagel, Düren, Germany). Construction of libraries for single-nucleotide variant detection and generation of 150 bp paired-end reads on an Illumina Novaseq 6000 platform, were performed by Novogene (Cambridge, United Kingdom). Reads were mapped to the reference genome of *E. coli* MG1655 (GenBank accession no. U000913.3). *breseq* pipeline (57) was applied to map the reads against the reference for identification of genomic variants, including SNPs and insertion-deletion polymorphisms (INDELs).

### Growth experiments

4 mL M9 medium containing 10 mM glycerol and 40 mM succinate (permissive growth condition) was used as pre-cultures for growth experiments. Strains were harvested (6,000\**g*, 3 min, RT) and washed three times in M9 medium without carbon source. Cultures were inoculated into the M9 media to an OD_600_ of 0.01 in a 96-well microtiter plate (Nunclon Delta Surface, Thermo Scientific). Each well contained 150 μL of culture and 50 μL mineral oil (Sigma-Aldrich) to avoid evaporation. Growth monitoring and incubation at 37 °C was carried out in a microplate reader (EPOCH 2, BioTek). In the program 4 shaking phases of 60 seconds were repeated three times (linear shaking 567 cpm (3 mm), orbital shaking 282 cpm (3 mm), linear shaking 731 cpm (2 mm), orbital shaking 365 cpm (2 mm)). After the shaking cycles absorbance at 600 nm was measured. Raw data were converted to 1 cm-wide standard cuvette OD values according to OD_cuvette_ = OD_plate_ / 0.23. Matlab was used to calculate growth parameters. All experiments were carried out in at least three replicates. Average values were used to generate the growth curves. Variability between triplicate measurements was less than 5% in all cases displayed.

### Expression analysis by reverse transcriptase quantitative PCR (RT-qPCR)

mRNA levels of *mgsA, serA, serB, serC and sdaA* were determined by RT-qPCR. Cells were harvested in exponential phase (OD_600_ 0.5-0.6) on M9 minimal medium cultures with either 20 mM glycerol (WT and Δ*tpi*, iso1) or 4 mM glycerol and 40 mM succinate (WT, Δ*tpi*, Δ*pgk*) as carbon source. RNA was extracted by using the RNeasy Mini Kit (Qiagen, Hilden, Germany) as described in the manufacturer’ s manual. In brief, ∼2.5×10^8^ cells (0.5 ml of OD_600_ 0.5) were mixed with 2 volumes of RNAprotect Bacteria Reagent (Qiagen, Hilden, Germany) and pelleted, followed by enzymatic lysis, on-column removal of genomic DNA with RNase-free DNase (Qiagen, Hilden, Germany) and spin-column-based purification of RNA. Integrity and concentration of the isolated RNA were determined by NanoDrop and gel electrophoresis. Reverse transcription to synthesize cDNA was performed on 500 ng RNA with the qScript cDNA Synthesis Kit (QuantaBio, Beverly, MA USA). Quantitative real-time PCR was performed in technical triplicates per three biological replicates using the Maxima SYBR Green/ROX qPCR Master Mix (Thermo Scientific, Dreieich, Germany). An input corresponding to 20.833 pg total RNA was used per reaction. Non-specific amplification products were excluded by melting curve analysis. The gene encoding 16S rRNA (*rrsA*) was chosen as a well-established reference transcript for expression normalization (58). Primer pairs for amplification of *mgsA, serA, serB, serC and sdaA* used are shown in Supplementary Table S3. Equal amplification efficiencies between the primers for the genes of interest and the reference gene were assumed. Negative control assays with direct input of RNA (without previous reverse transcription) confirmed that residual genomic DNA contributed to less than 9% of the signal (ΔCt between +RT/-RT samples >12 for all). Differences in expression levels were calculated according to the 2^-ΔΔCT^ method (59, 60). Reported data represents the 2^-ΔΔCT^ value that was calculated for each sample individually relative to the average of all biological WT replicate ΔCt^(Ct(GOI)-Ct(rrsA))^ values.

### Protein Expression and purification

The His-tagged WT and mutated SerA proteins were expressed in *E. coli* BL21 (DE3) Codon+ (Invitrogen). Cells in 400 ml Terrific broth containing 100 µg/mL carbenicillin were grown at 37°C until they reached an OD_600nm_ = 2 upon which expression for 16 h at 20 °C was induced by addition 500 μM IPTG. Cells were harvested by centrifugation for 30 min at 10000 g at 4°C. Cell pellets were frozen at -80°C for one night. Thawed cells were then suspended in 32 ml of Buffer A (50 mM phosphate (Na/K), 500 mM NaCl, 30 mM imidazole, 15% glycerol, pH 8.0) and lysed for 30 min at room temperature after addition of 3.6 ml of Bug Buster (Novagen), 32 µl DTT (dithiothreitol) 1M, 320 µl Pefabloc 0.1 M (Millipore) and 23 µl Lysonase (Novagen). Lysate was clarified at 9000g for 45 min at 4°C then loaded onto a 5 ml HisTrap FF column pre-equilibrated in Buffer A. The protein was eluted in Buffer B (50 mM phosphate (Na/K), 500 mM NaCl, 250 mM imidazole, 1 mM DTT, 15% glycerol, pH 8.0) and desalted on a gel-filtration column Hi Load 16/60 Superdex 200 pg in Buffer C (50mM Tris, 50 mM NaCl, glycerol 15%, 1 mM DTT, pH8.0). The protein was frozen and stored at -80°C if not immediately used for assays.

### Characterization of SerA kinetic parameters

Assays were performed using a Safas UV mc2 double beam spectrophotometer at room temperature using quartz cuvettes (0.6 cm path length). Assays of SerA-catalyzed reduction of 2-ketoglutarate were conducted in 40 mM potassium phosphate, 1 mM DTT, pH 7.5. Kinetic parameters for 2-ketoglutarate were determined by varying its concentration (from 0.005 to 1 mM) in the presence of a saturating concentration of NADH (250 µM). The reactions were monitored by recording the disappearance of NADH at 340 nm (molar extinction coefficient = 6220 M^-1^.cm^-1^). Kinetic constants were determined by non-linear analysis of initial rates from duplicate experiments using SigmaPlot 9.0 (Systat Software, Inc.).

### Characterization of serine inhibition of SerA 2-ketoglutarate reductase activity

Standard enzyme assays were conducted with both substrates at saturating concentrations (250 µM NADH, 600 µM 2-ketoglutarate) in 120 µL final volume. Reaction mixes contained 5 µg of purified enzyme. Serine was added at concentrations ranging from 0.5 to 10 µM.

## Acknowledgments

We thank Enrico Orsi for critical reading of the manuscript. This study was funded by the Max Planck Society and the CEA Genoscope.

## Data availability

The data supporting the presented findings are available within the paper or in its Supplementary files. Strains and plasmids used here are available on request from the corresponding author. Data from the following public repositories were used in this study: BiGG [http://bigg.ucsd.edu/], KEGG [https://www.kegg.jp/], and eQuilibrator [http://equilibrator.weizmann.ac.il/].

## Code availability

The code used in this study can be found at GitLab [https://gitlab.com/elad.noor/glycolysis-bypass/-/tree/master/results].

## Author contributions

S.N.L. and A.B.-E. conceived and supervised the study. M.B., V.D., A.B.-E. and S.N.L. designed the experiments. E.N. performed the computational work. C.I., K.M., M.H., V.B., H.S.M. and A.S. constructed plasmids and strains and performed growth experiments. M.B. and V.D. supervised the evolution experiments and analyzed the genome sequencing data. V.A.D. and I.D. ran the continuous cultures, isolated and characterized evolved strains. A.B. performed biochemical experiments. C.I. and B.D. performed qPCR experiments. S.N.L., V.D., and M.B. analyzed the results and wrote the manuscript with contributions from all authors.

## Competing Interest Statement

The authors declare no competing interests.

## Notes

### Competing Interest Statement

The authors have declared no competing interest.

