## Supplementary Material for "Activating silent glycolysis bypasses in *Escherichia coli*"

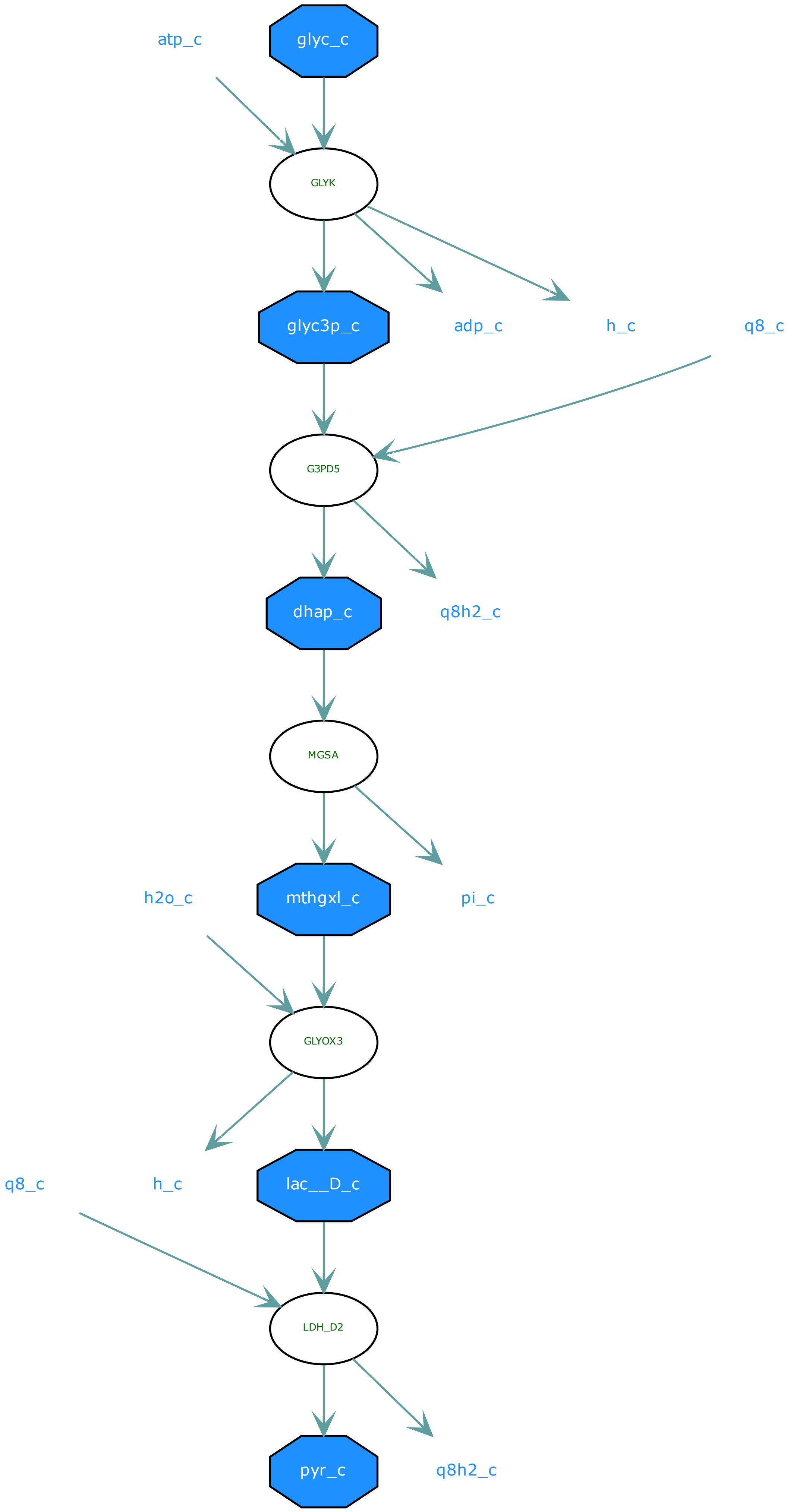

glyc_c
atp_c
GLYK
glyc3p_c
adp_c
h_c
q8_c
G3PD5
dhap_c
q8h2_c
MGSA
mthgxl_c
h2o_c
pi_c
GLYOX3
lac__D_c
q8_c
h_c
LDH_D2
pyr_c
q8h2_c

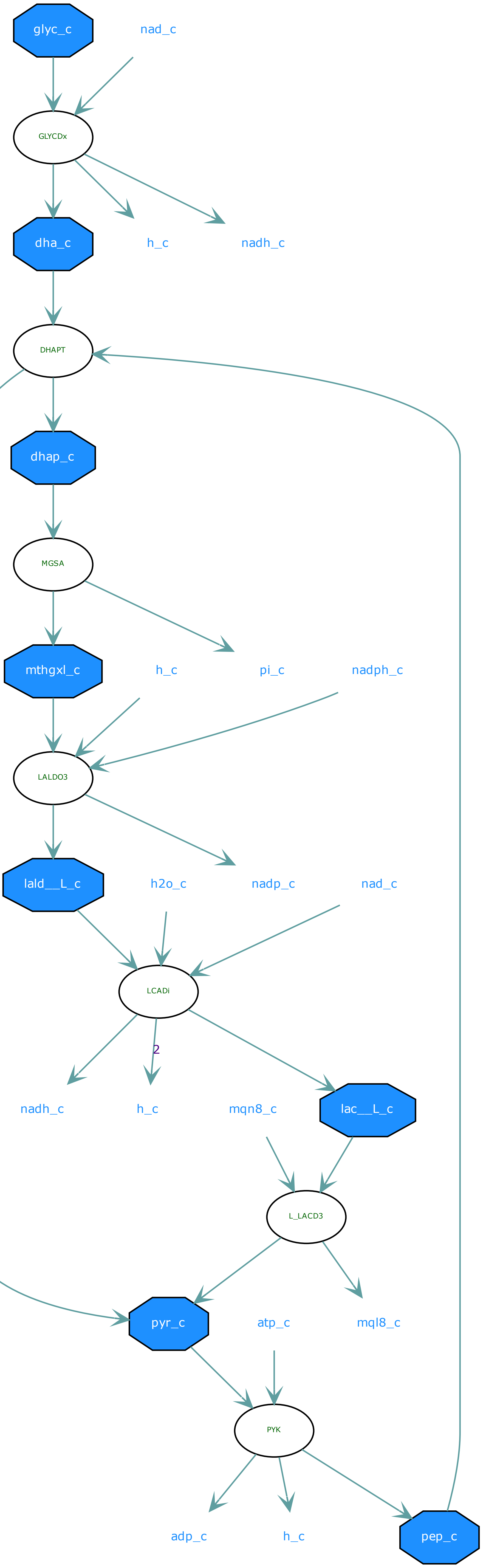

glyc_c
nad_c
GLYCDx
dha_c
h_c
nadh_c
DHAPT
dhap_c
MGSA
mthgxl_c
h_c
pi_c
nadph_c
LALDO3
lald__L_c
h2o_c
nadp_c
nad_c
LCADi
2
nadh_c
h_c
mqn8_c
lac__L_c
L_LACD3
pyr_c
atp_c
mql8_c
PYK
adp_c
h_c
pep_c

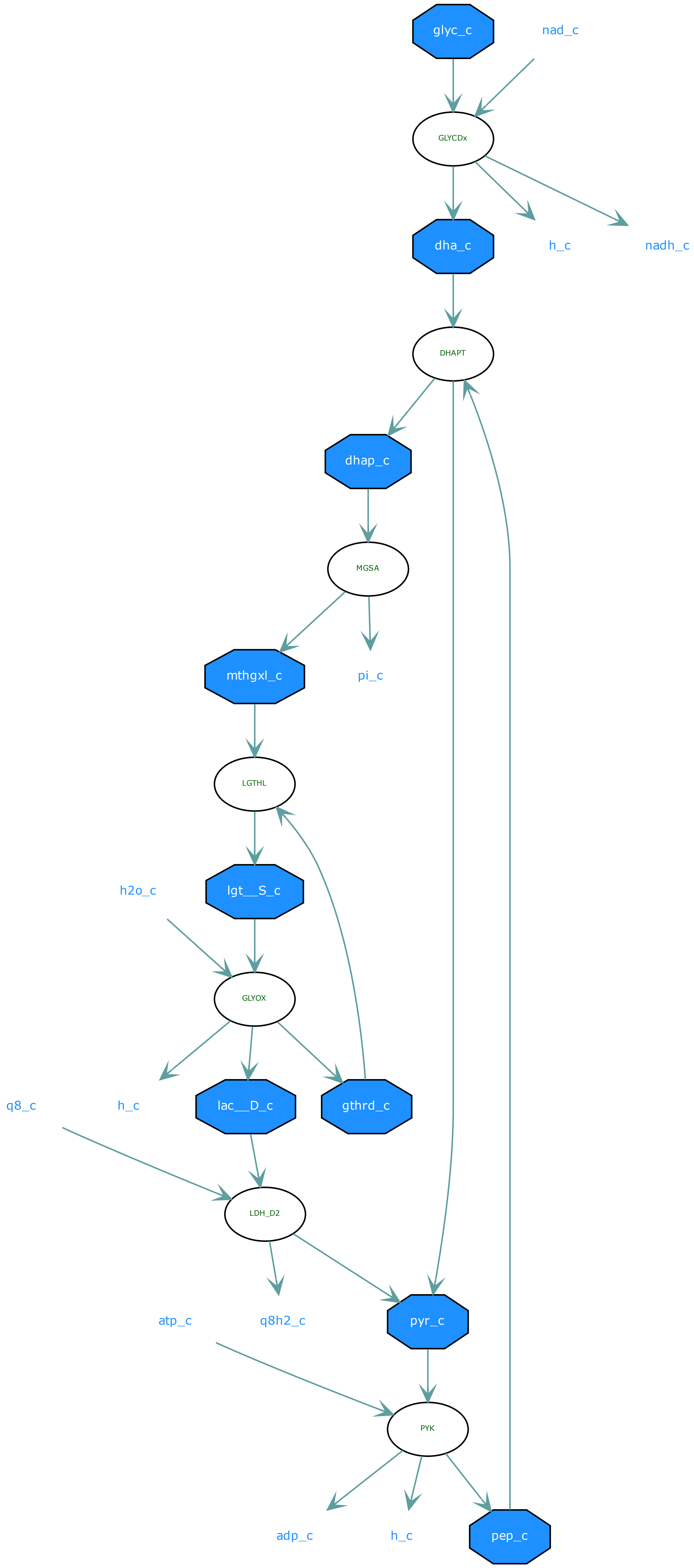

glyc_c
nad_c
GLYCDx
dha_c
h_c
nadh_c
DHAPT
dhap_c
MGSA
mthgxl_c
pi_c
LGTHL
lgt__S_c
h2o_c
GLYOX
q8_c
h_c
lac__D_c
gthrd_c
LDH_D2
atp_c
q8h2_c
pyr_c
PYK
adp_c
h_c
pep_c

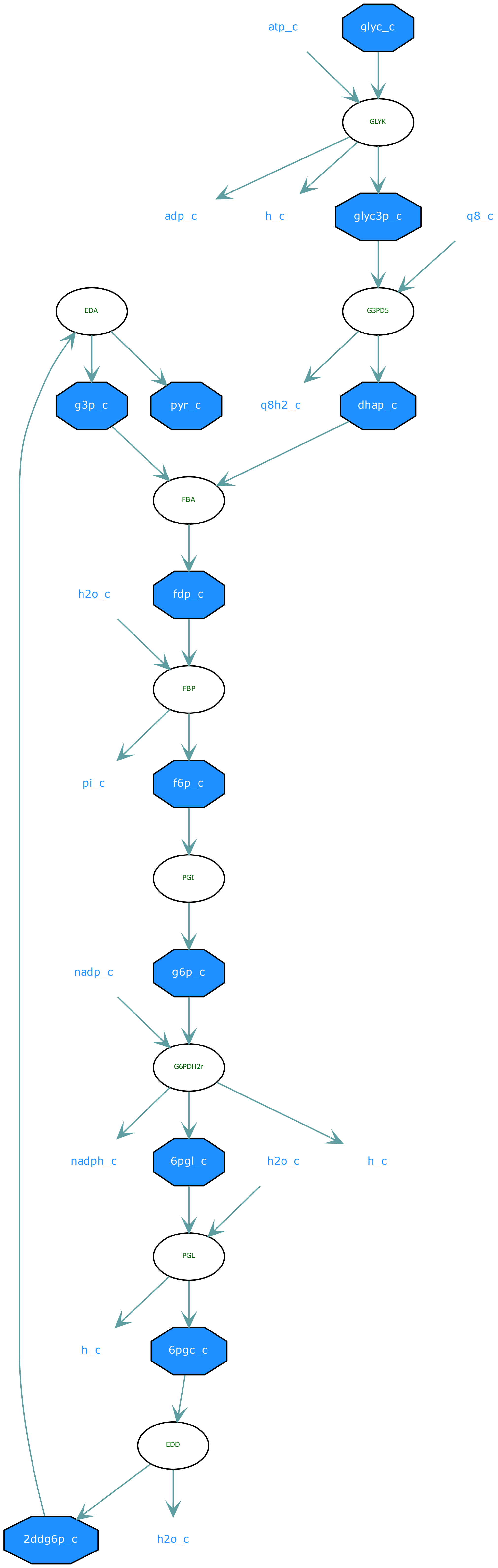
atp_c
glyc_c
GLYK
adp_c
h_c
glyc3p_c
q8_c
EDA
G3PD5
g3p_c
pyr_c
q8h2_c
dhap_c
FBA
h2o_c
fdp_c
FBP
pi_c
f6p_c
PGI
nadp_c
g6p_c
G6PDH2r
nadph_c
6pgl_c
h2o_c
h_c
PGL
h_c
6pgc_c
EDD
2ddg6p_c
h2o_c

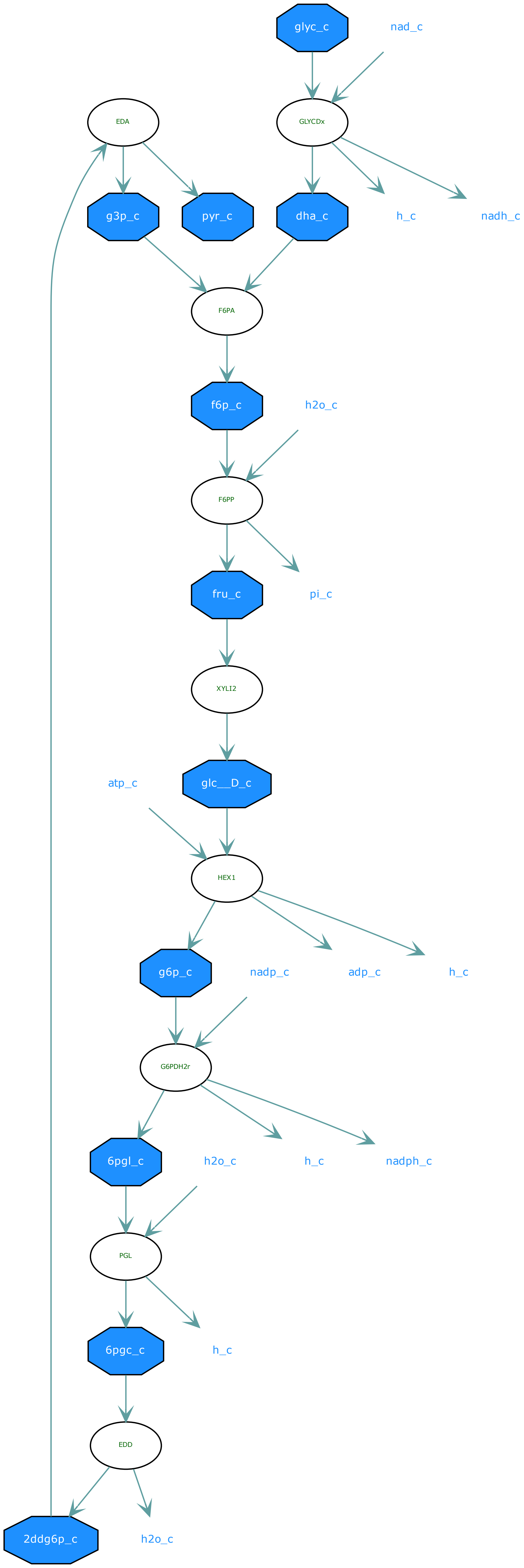
glyc_c
nad_c
EDA
GLYCDx
g3p_c
pyr_c
dha_c
h_c
nadh_c
F6PA
f6p_c
h2o_c
F6PP
fru_c
pi_c
XYLI2
atp_c
glc__D_c
HEX1
g6p_c
nadp_c
adp_c
h_c
G6PDH2r
6pgl_c
h2o_c
h_c
nadph_c
PGL
6pgc_c
h_c
EDD
2ddg6p_c
h2o_c

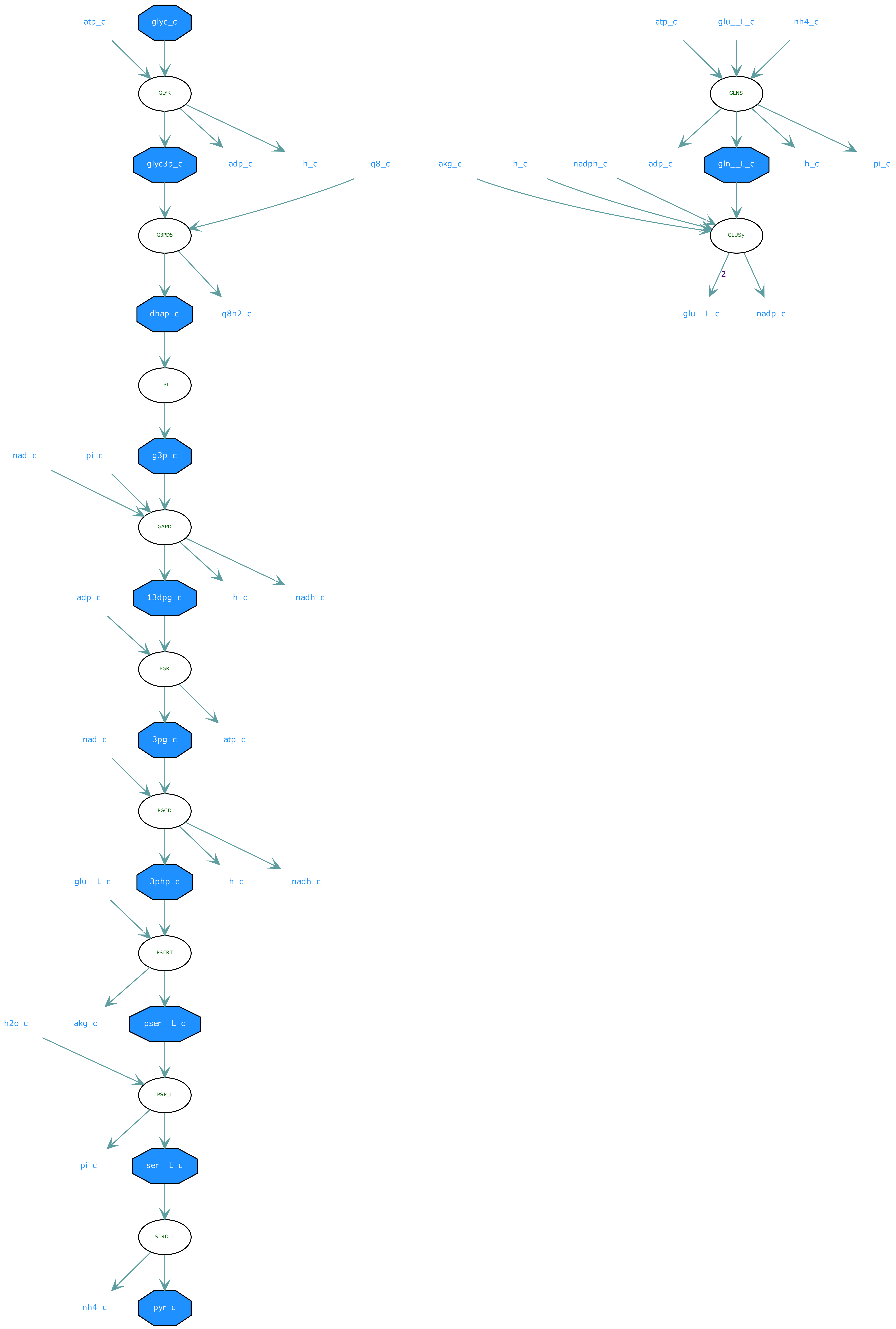
atp_c
glyc_c
GLYK
glyc3p_c
adp_c
h_c
q8_c
G3PD5
dhap_c
q8h2_c
TPI
g3p_c
nad_c
pi_c
GAPD
13dpg_c
h_c
nadh_c
adp_c
PGK
3pg_c
atp_c
nad_c
PGCD
3php_c
h_c
nadh_c
glu__L_c
PSERT
pser__L_c
akg_c
h2o_c
PSP_L
ser__L_c
pi_c
SERD_L
pyr_c
nh4_c
atp_c
glu__L_c
nh4_c
GLNS
adp_c
gln__L_c
h_c
pi_c
akg_c
h_c
nadph_c
GLUSy
2
glu__L_c
nadp_c

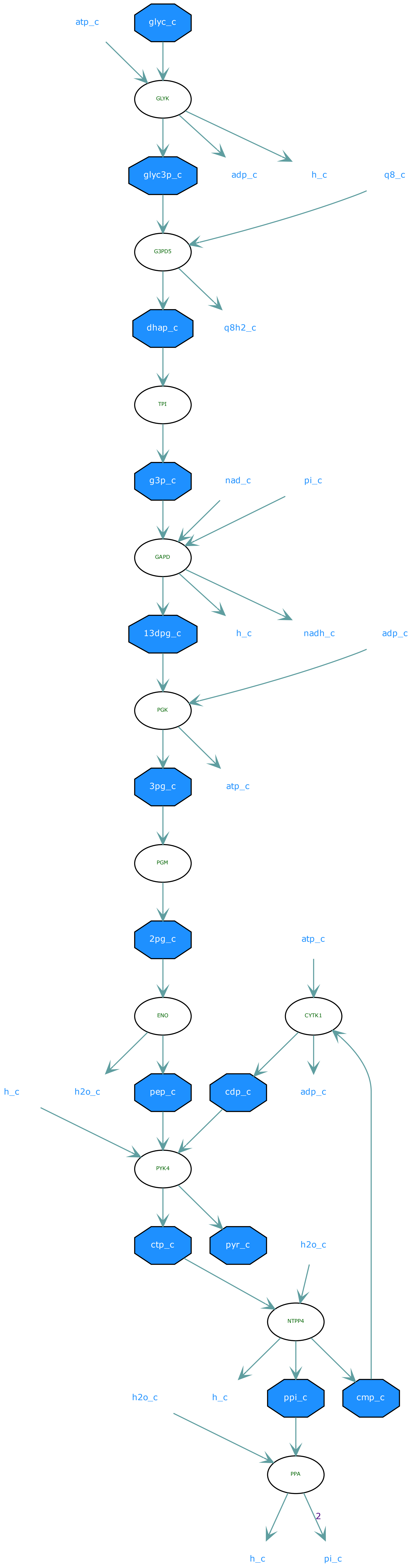

atp_c
glyc_c
GLYK
glyc3p_c
adp_c
h_c
q8_c
G3PD5
dhap_c
q8h2_c
TPI
g3p_c
nad_c
pi_c
GAPD
13dpg_c
h_c
nadh_c
adp_c
PGK
3pg_c
atp_c
PGM
2pg_c
atp_c
ENO
CYTK1
h_c
h2o_c
pep_c
cdp_c
adp_c
PYK4
ctp_c
pyr_c
h2o_c
NTPP4
h2o_c
h_c
ppi_c
cmp_c
PPA
2
h_c
pi_c

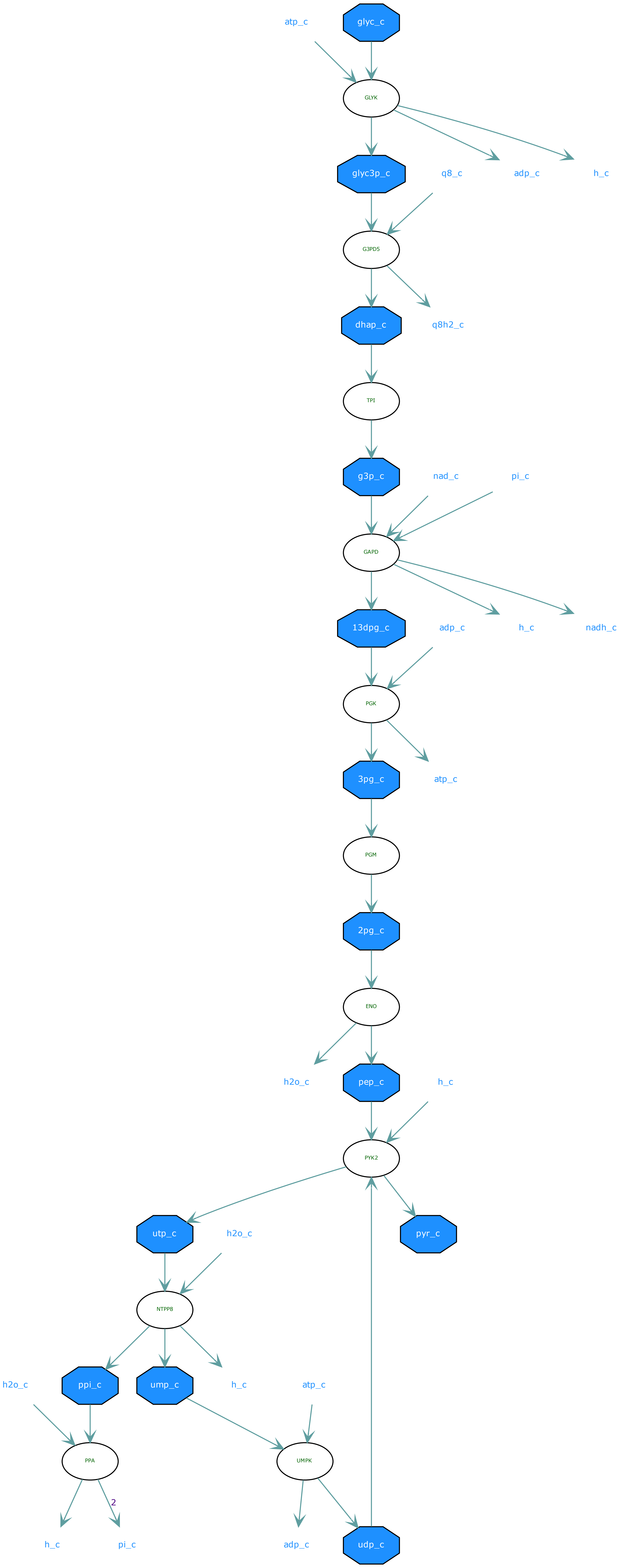
atp_c
glyc_c
GLYK
glyc3p_c
q8_c
adp_c
h_c
G3PD5
dhap_c
q8h2_c
TPI
g3p_c
nad_c
pi_c
GAPD
13dpg_c
adp_c
h_c
nadh_c
PGK
3pg_c
atp_c
PGM
2pg_c
ENO
h2o_c
pep_c
h_c
PYK2
utp_c
h2o_c
pyr_c
NTPP8
h2o_c
ppi_c
ump_c
h_c
atp_c
PPA
UMPK
2
h_c
pi_c
adp_c
udp_c

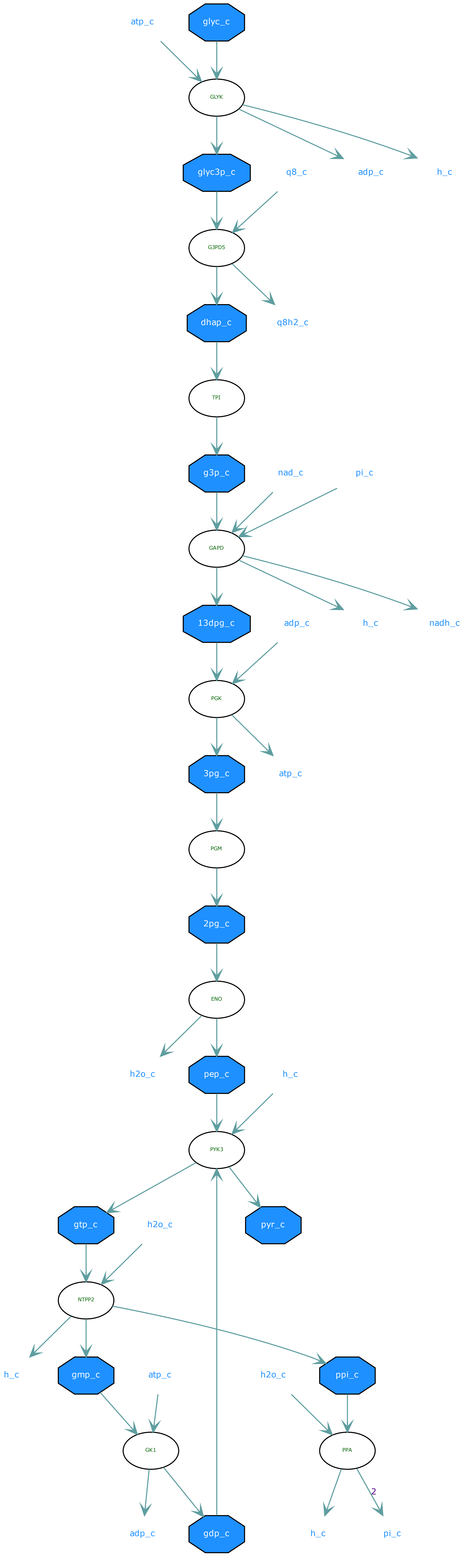

atp_c
glyc_c
GLYK
glyc3p_c
q8_c
adp_c
h_c
G3PD5
dhap_c
q8h2_c
TPI
g3p_c
nad_c
pi_c
GAPD
13dpg_c
adp_c
h_c
nadh_c
PGK
3pg_c
atp_c
PGM
2pg_c
ENO
h2o_c
pep_c
h_c
PYK3
gtp_c
h2o_c
pyr_c
NTPP2
h_c
gmp_c
atp_c
h2o_c
ppi_c
GK1
PPA
2
adp_c
gdp_c
h_c
pi_c

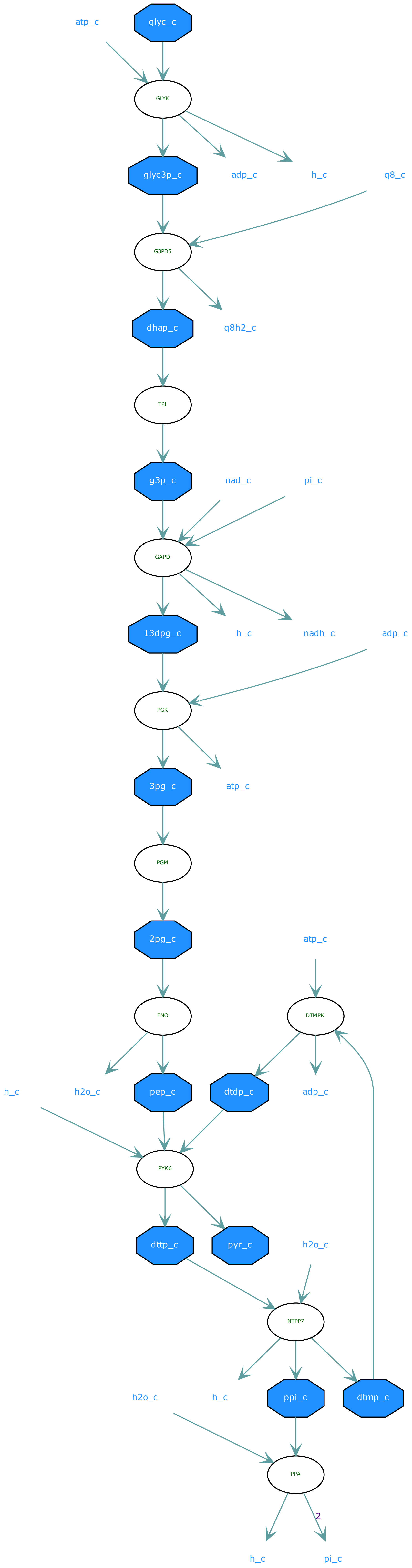

atp_c
glyc_c
GLYK
glyc3p_c
adp_c
h_c
q8_c
G3PD5
dhap_c
q8h2_c
TPI
g3p_c
nad_c
pi_c
GAPD
13dpg_c
h_c
nadh_c
adp_c
PGK
3pg_c
atp_c
PGM
2pg_c
atp_c
ENO
DTMPK
h_c
h2o_c
pep_c
dtdp_c
adp_c
PYK6
dttp_c
pyr_c
h2o_c
NTPP7
h2o_c
h_c
ppi_c
dtmp_c
PPA
2
h_c
pi_c

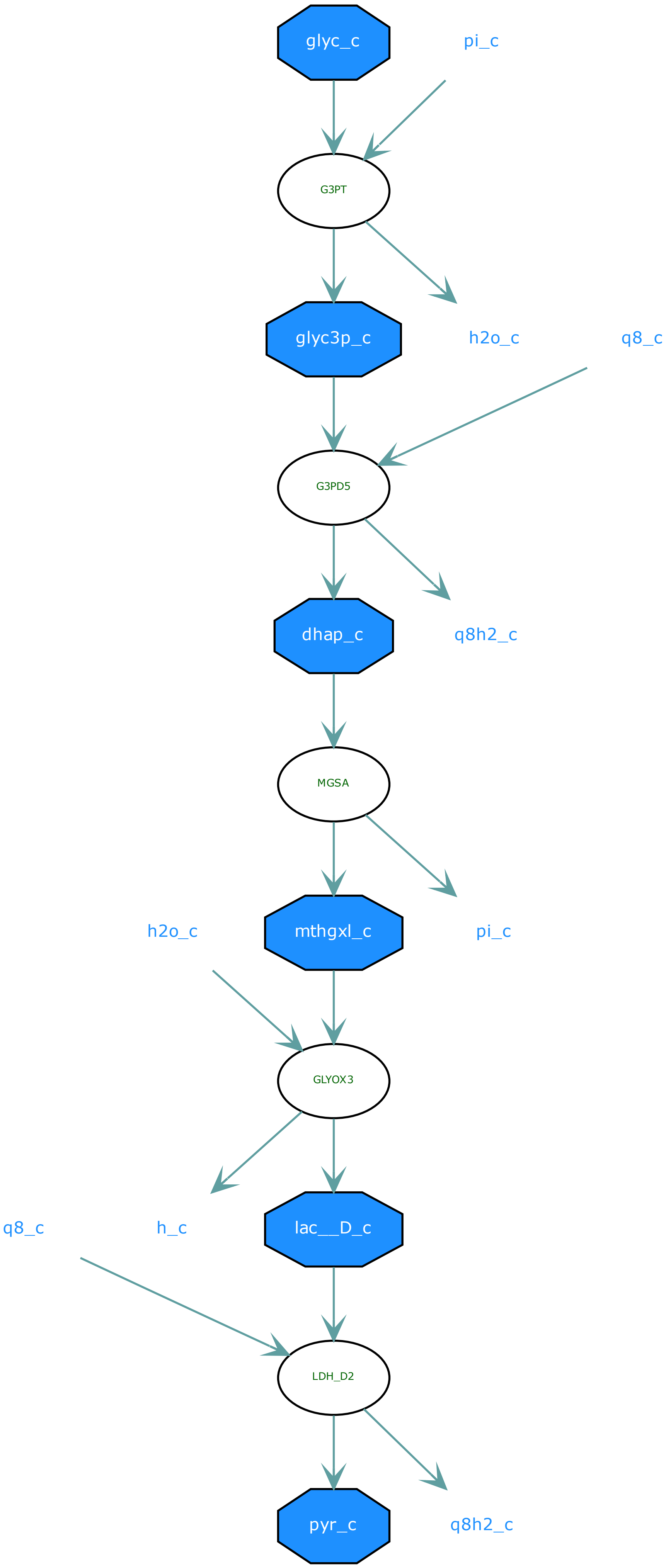
glyc_c
pi_c
G3PT
glyc3p_c
h2o_c
q8_c
G3PD5
dhap_c
q8h2_c
MGSA
h2o_c
mthgxl_c
pi_c
GLYOX3
q8_c
h_c
lac__D_c
LDH_D2
pyr_c
q8h2_c

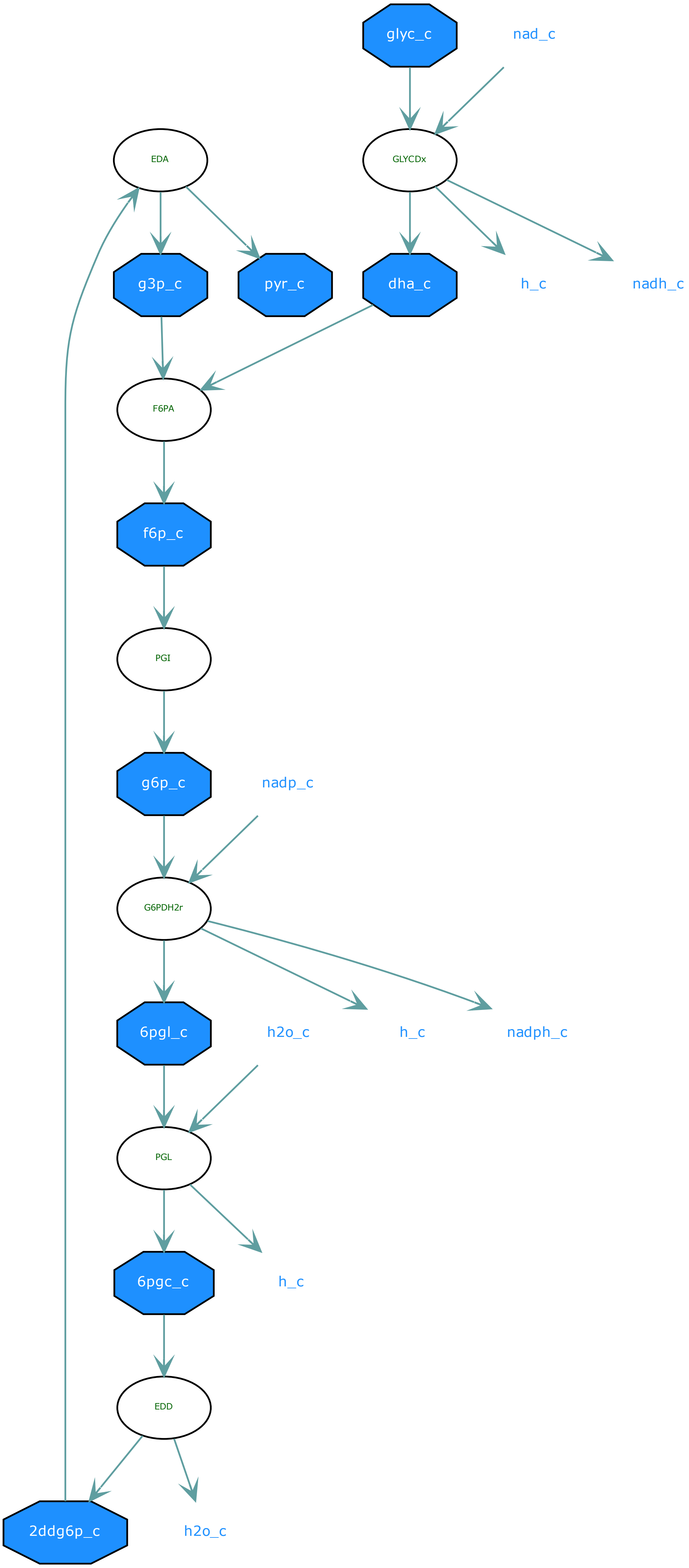
glyc_c
nad_c
EDA
GLYCDx
g3p_c
pyr_c
dha_c
h_c
nadh_c
F6PA
f6p_c
PGI
g6p_c
nadp_c
G6PDH2r
6pgl_c
h2o_c
h_c
nadph_c
PGL
6pgc_c
h_c
EDD
2ddg6p_c
h2o_c

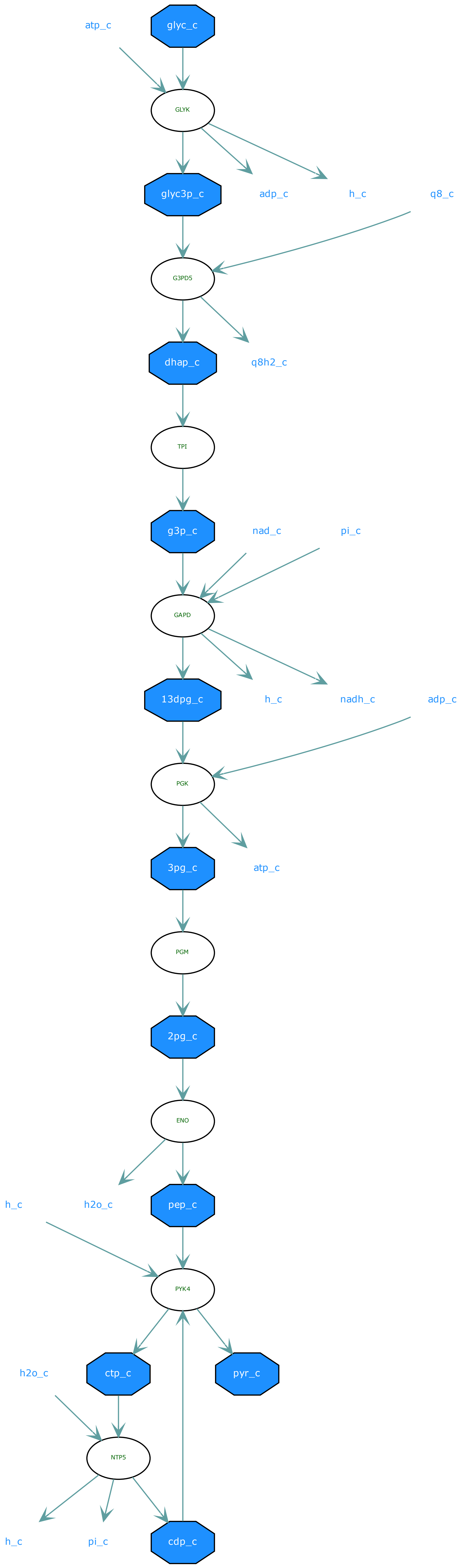

atp_c
glyc_c
GLYK
glyc3p_c
adp_c
h_c
q8_c
G3PD5
dhap_c
q8h2_c
TPI
g3p_c
nad_c
pi_c
GAPD
13dpg_c
h_c
nadh_c
adp_c
PGK
3pg_c
atp_c
PGM
2pg_c
ENO
h_c
h2o_c
pep_c
PYK4
h2o_c
ctp_c
pyr_c
NTP5
h_c
pi_c
cdp_c

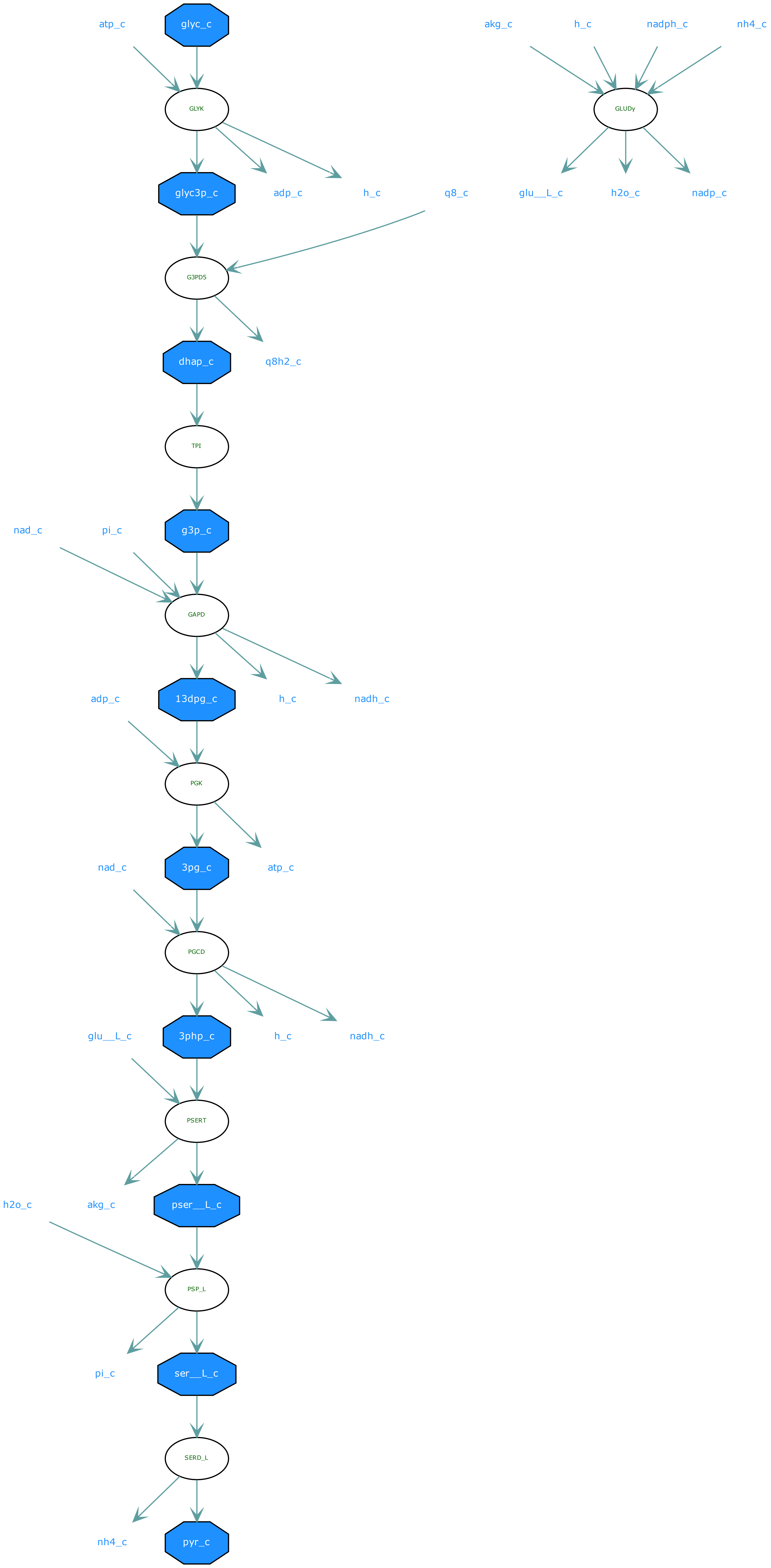
atp_c
glyc_c
akg_c
h_c
nadph_c
nh4_c
GLYK
GLUDy
glyc3p_c
adp_c
h_c
q8_c
glu__L_c
h2o_c
nadp_c
G3PD5
dhap_c
q8h2_c
TPI
nad_c
pi_c
g3p_c
GAPD
adp_c
13dpg_c
h_c
nadh_c
PGK
nad_c
3pg_c
atp_c
PGCD
glu__L_c
3php_c
h_c
nadh_c
PSERT
h2o_c
akg_c
pser__L_c
PSP_L
pi_c
ser__L_c
SERD_L
nh4_c
pyr_c

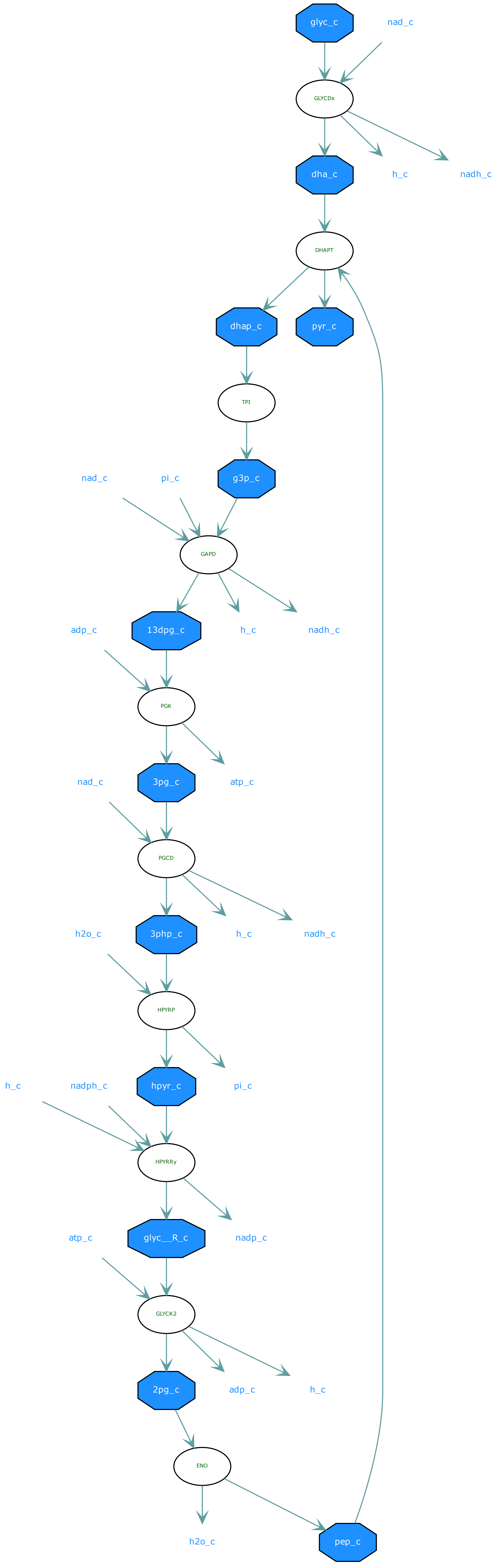
glyc_c
nad_c
GLYCDx
dha_c
h_c
nadh_c
DHAPT
dhap_c
pyr_c
TPI
nad_c
pi_c
g3p_c
GAPD
adp_c
13dpg_c
h_c
nadh_c
PGK
nad_c
3pg_c
atp_c
PGCD
h2o_c
3php_c
h_c
nadh_c
HPYRP
h_c
nadph_c
hpyr_c
pi_c
HPYRRy
atp_c
glyc__R_c
nadp_c
GLYCK2
2pg_c
adp_c
h_c
ENO
h2o_c
pep_c

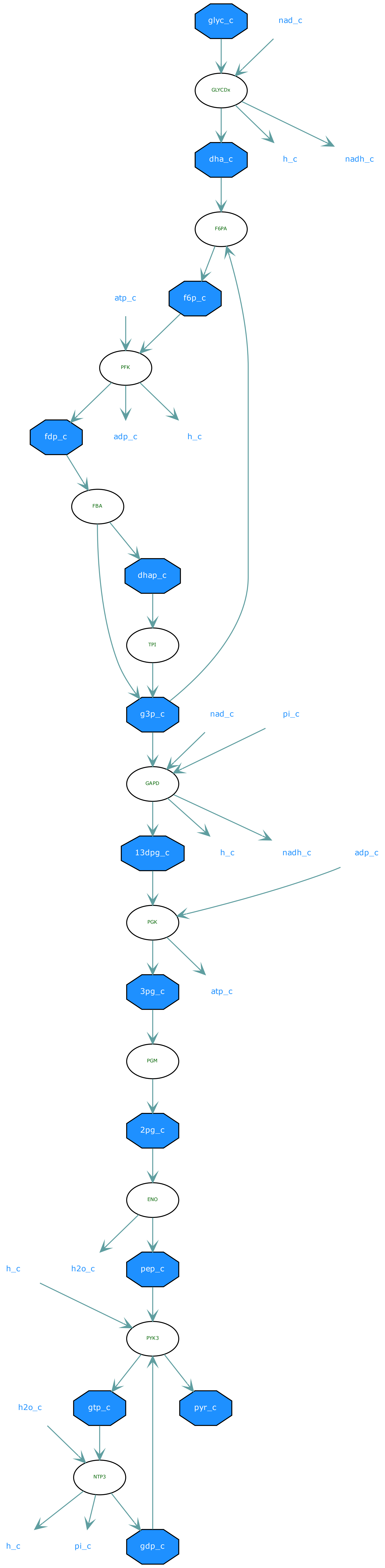
glyc_c
nad_c
GLYCDx
dha_c
h_c
nadh_c
F6PA
atp_c
f6p_c
PFK
fdp_c
adp_c
h_c
FBA
dhap_c
TPI
g3p_c
nad_c
pi_c
GAPD
13dpg_c
h_c
nadh_c
adp_c
PGK
3pg_c
atp_c
PGM
2pg_c
ENO
h_c
h2o_c
pep_c
PYK3
h2o_c
gtp_c
pyr_c
NTP3
h_c
pi_c
gdp_c

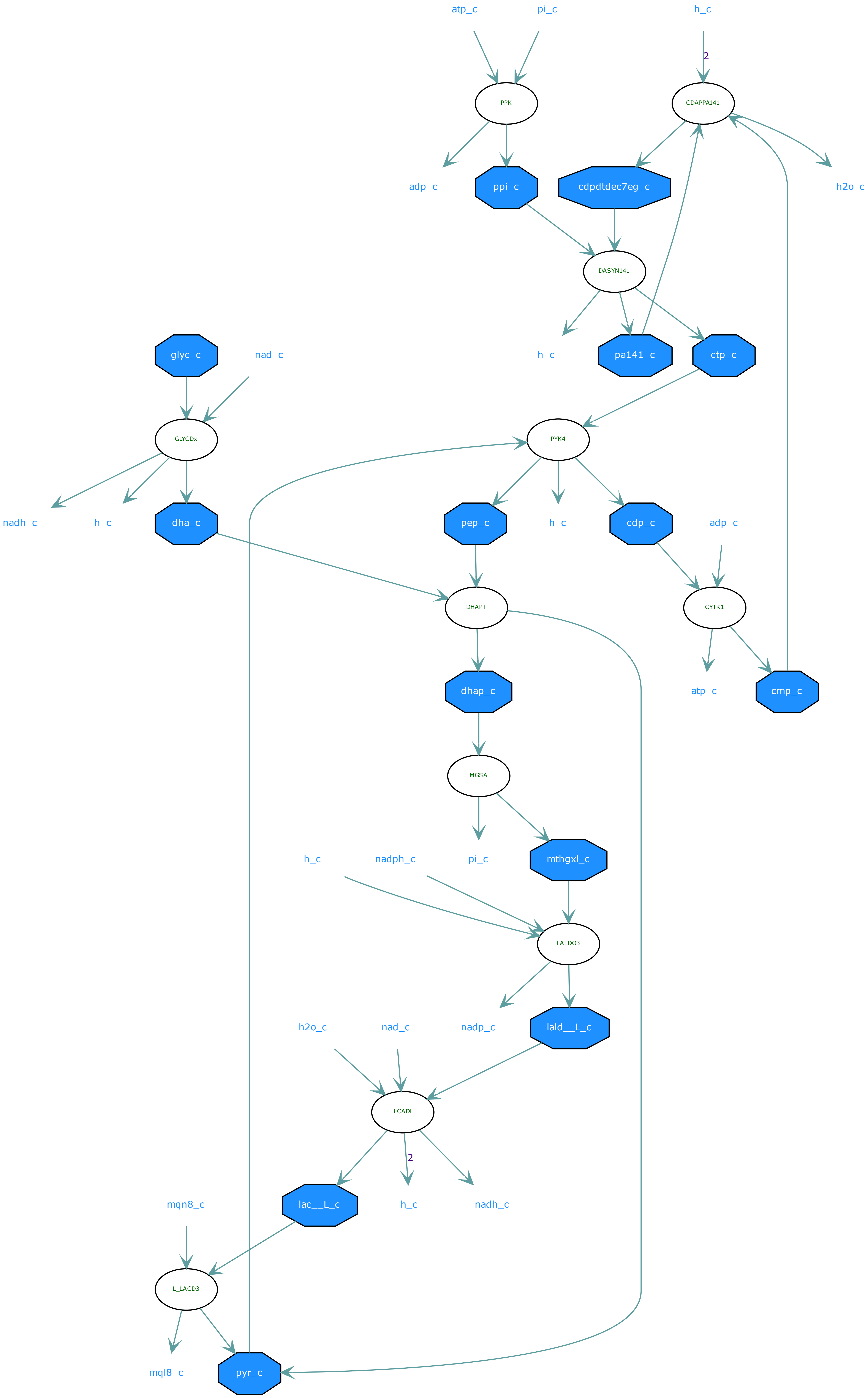

atp_c
pi_c
h_c
2
PPK
CDAPPA141
adp_c
ppi_c
cdpdtdec7eg_c
h2o_c
DASYN141
glyc_c
nad_c
h_c
pa141_c
ctp_c
GLYCDx
PYK4
nadh_c
h_c
dha_c
pep_c
h_c
cdp_c
adp_c
DHAPT
CYTK1
dhap_c
atp_c
cmp_c
MGSA
h_c
nadph_c
pi_c
mthgxl_c
LALDO3
h2o_c
nad_c
nadp_c
lald__L_c
LCADi
2
mqn8_c
lac__L_c
h_c
nadh_c
L_LACD3
mql8_c
pyr_c

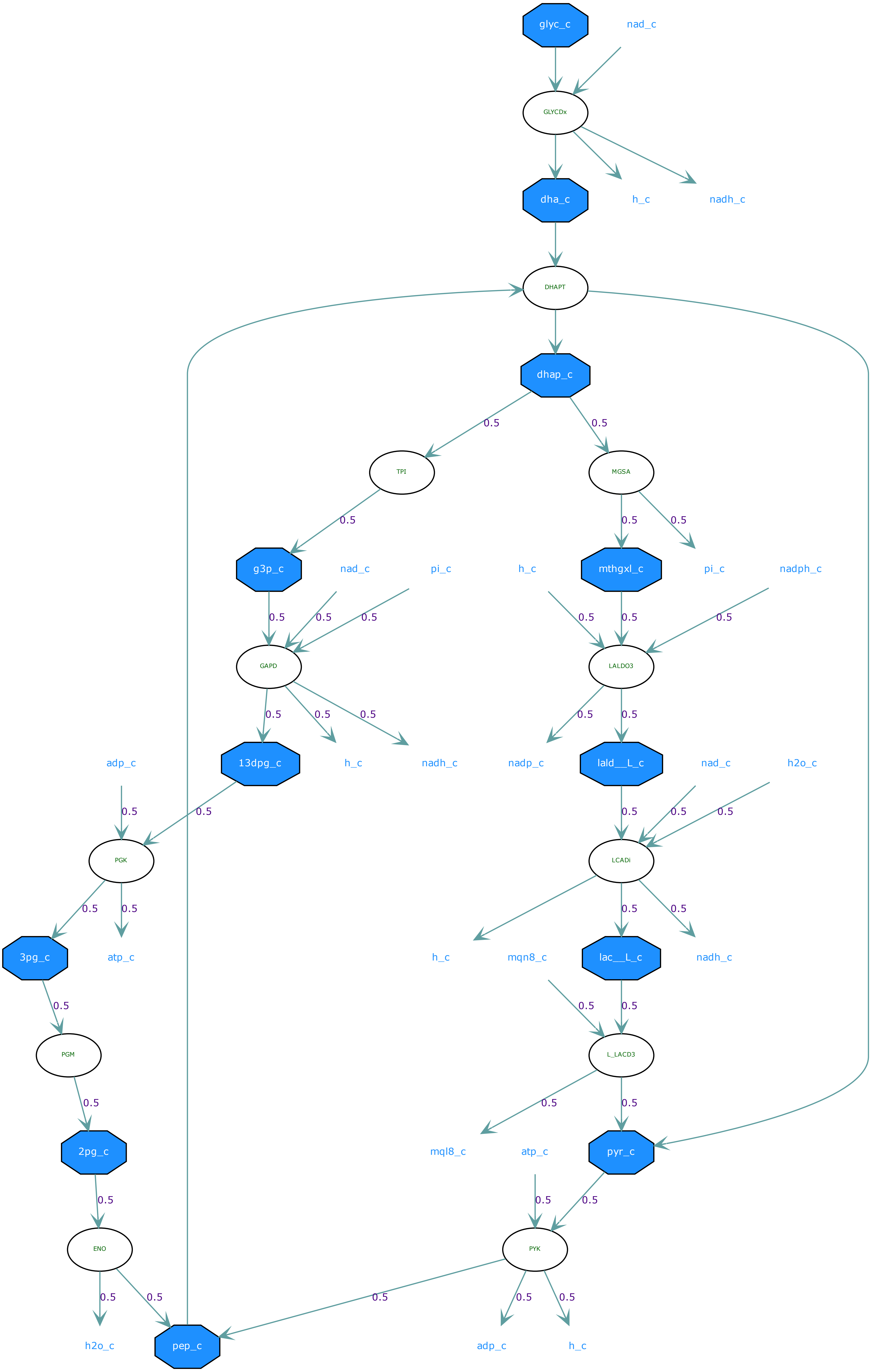
glyc_c
nad_c
GLYCDx
dha_c
h_c
nadh_c
DHAPT
dhap_c
TPI
MGSA
g3p_c
nad_c
pi_c
h_c
mthgxl_c
pi_c
nadph_c
GAPD
LALDO3
13dpg_c
h_c
nadh_c
nadp_c
lald__L_c
nad_c
h2o_c
adp_c
PGK
LCADi
3pg_c
atp_c
h_c
mqn8_c
lac__L_c
nadh_c
PGM
L_LACD3
2pg_c
mql8_c
atp_c
pyr_c
ENO
PYK
h2o_c
pep_c
adp_c
h_c
0.5

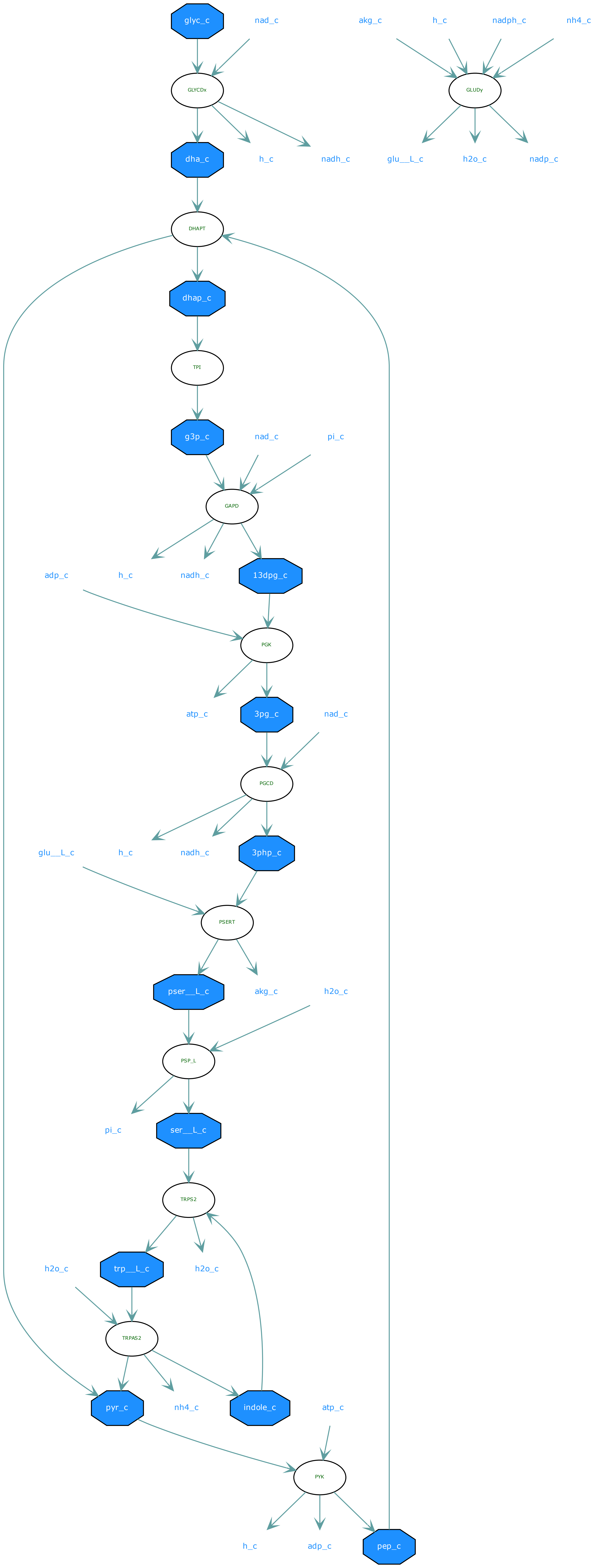
glyc_c
nad_c
akg_c
h_c
nadph_c
nh4_c
GLYCDx
GLUDy
dha_c
h_c
nadh_c
glu__L_c
h2o_c
nadp_c
DHAPT
dhap_c
TPI
g3p_c
nad_c
pi_c
GAPD
adp_c
h_c
nadh_c
13dpg_c
PGK
atp_c
3pg_c
nad_c
PGCD
glu__L_c
h_c
nadh_c
3php_c
PSERT
pser__L_c
akg_c
h2o_c
PSP_L
pi_c
ser__L_c
TRPS2
h2o_c
trp__L_c
h2o_c
TRPAS2
pyr_c
nh4_c
indole_c
atp_c
PYK
h_c
adp_c
pep_c

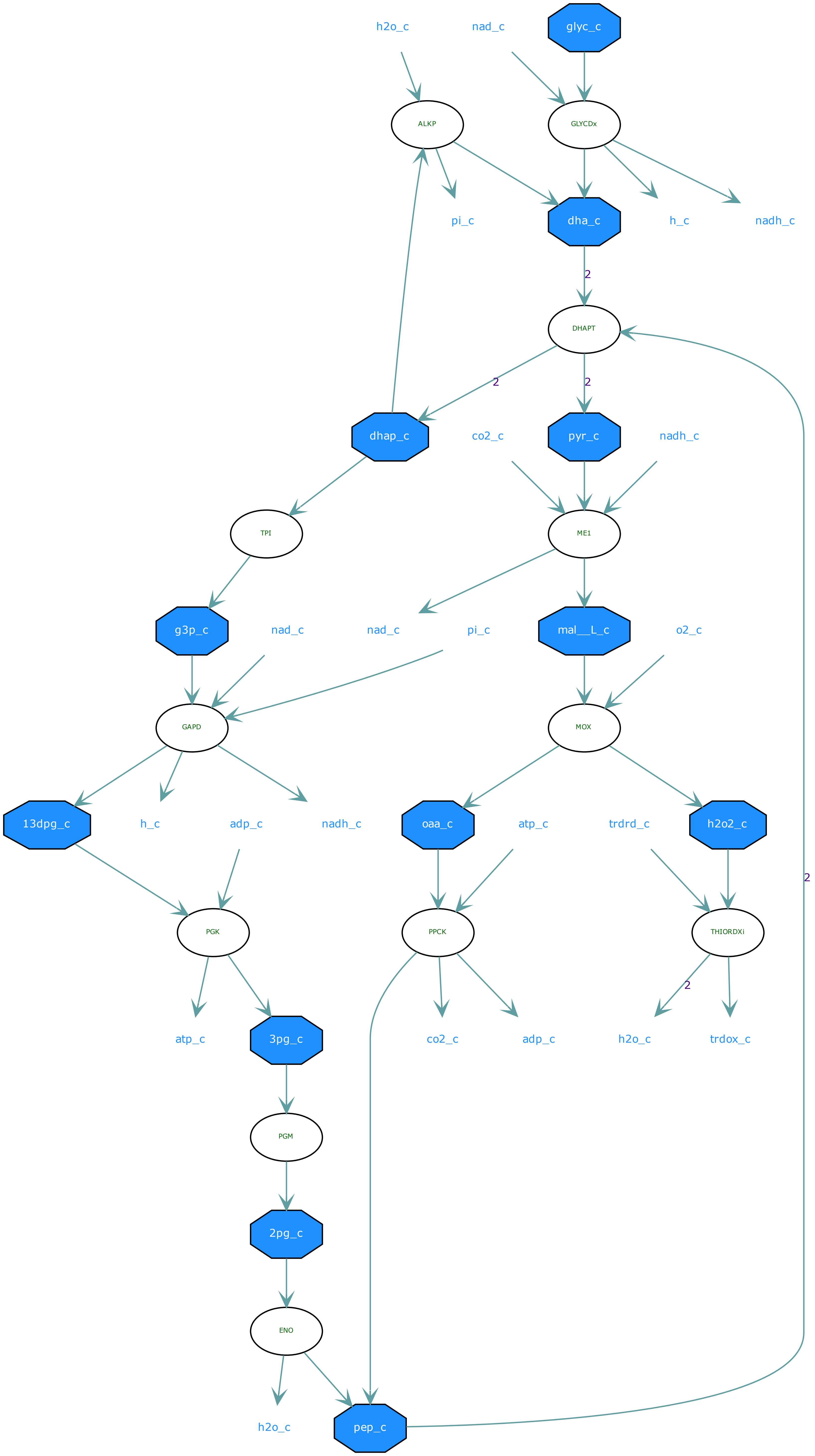
h2o_c
nad_c
glyc_c
ALKP
GLYCDx
pi_c
dha_c
h_c
nadh_c
2
DHAPT
2
2
dhap_c
co2_c
pyr_c
nadh_c
TPI
ME1
g3p_c
nad_c
nad_c
pi_c
mal__L_c
o2_c
GAPD
MOX
13dpg_c
h_c
adp_c
nadh_c
oaa_c
atp_c
trdrd_c
h2o2_c
2
PGK
PPCK
THIORDXi
2
atp_c
3pg_c
co2_c
adp_c
h2o_c
trdox_c
PGM
2pg_c
ENO
h2o_c
pep_c

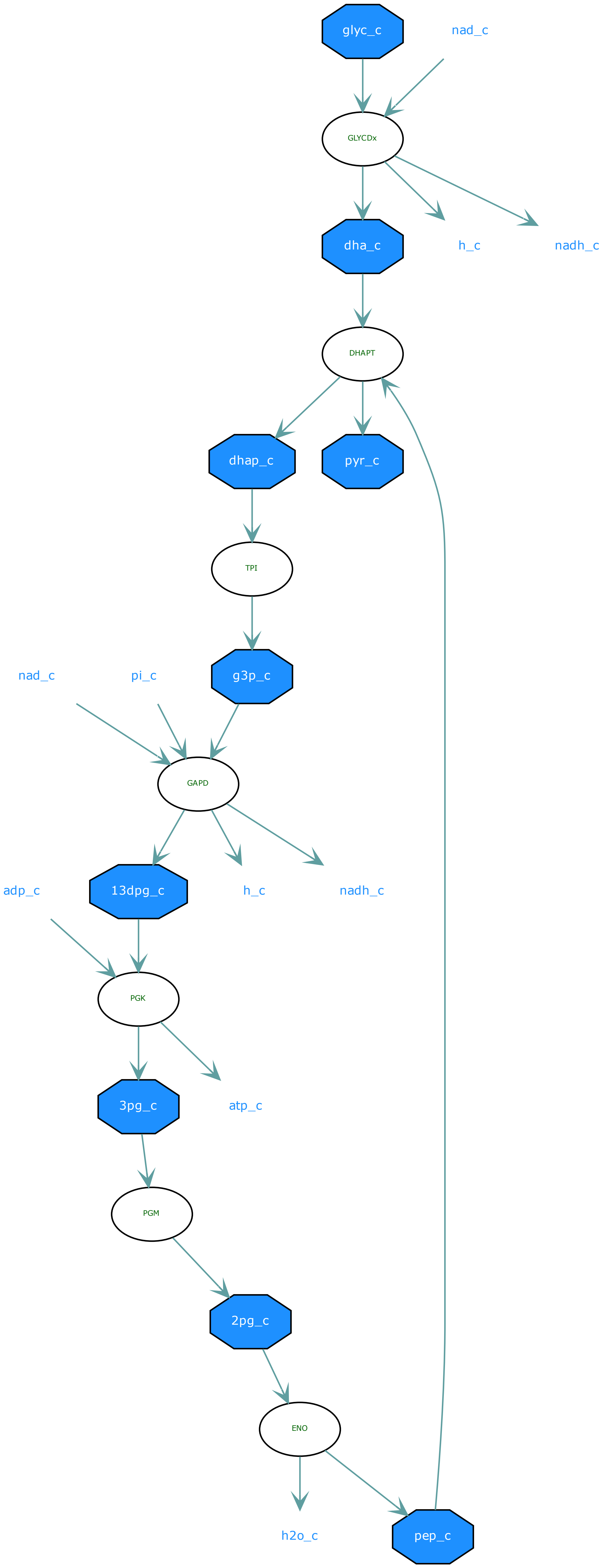
glyc_c
nad_c
GLYCDx
dha_c
h_c
nadh_c
DHAPT
dhap_c
pyr_c
TPI
nad_c
pi_c
g3p_c
GAPD
adp_c
13dpg_c
h_c
nadh_c
PGK
3pg_c
atp_c
PGM
2pg_c
ENO
h2o_c
pep_c

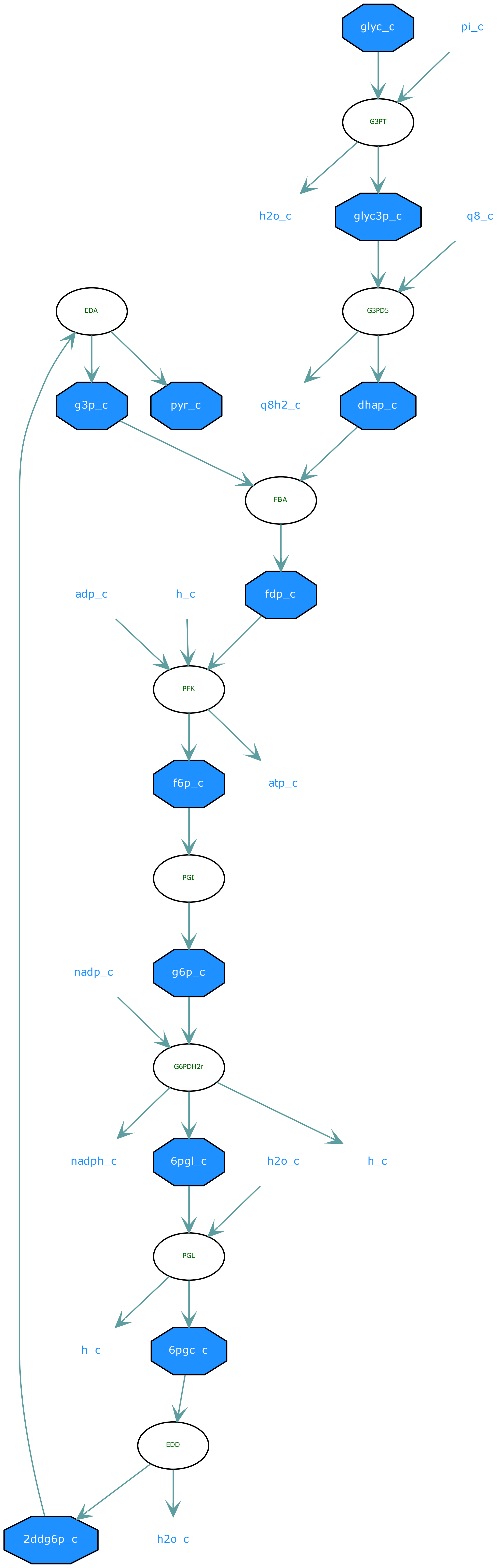

glyc_c
pi_c
G3PT
h2o_c
glyc3p_c
q8_c
EDA
G3PD5
g3p_c
pyr_c
q8h2_c
dhap_c
FBA
adp_c
h_c
fdp_c
PFK
f6p_c
atp_c
PGI
nadp_c
g6p_c
G6PDH2r
nadph_c
6pgl_c
h2o_c
h_c
PGL
h_c
6pgc_c
EDD
2ddg6p_c
h2o_c

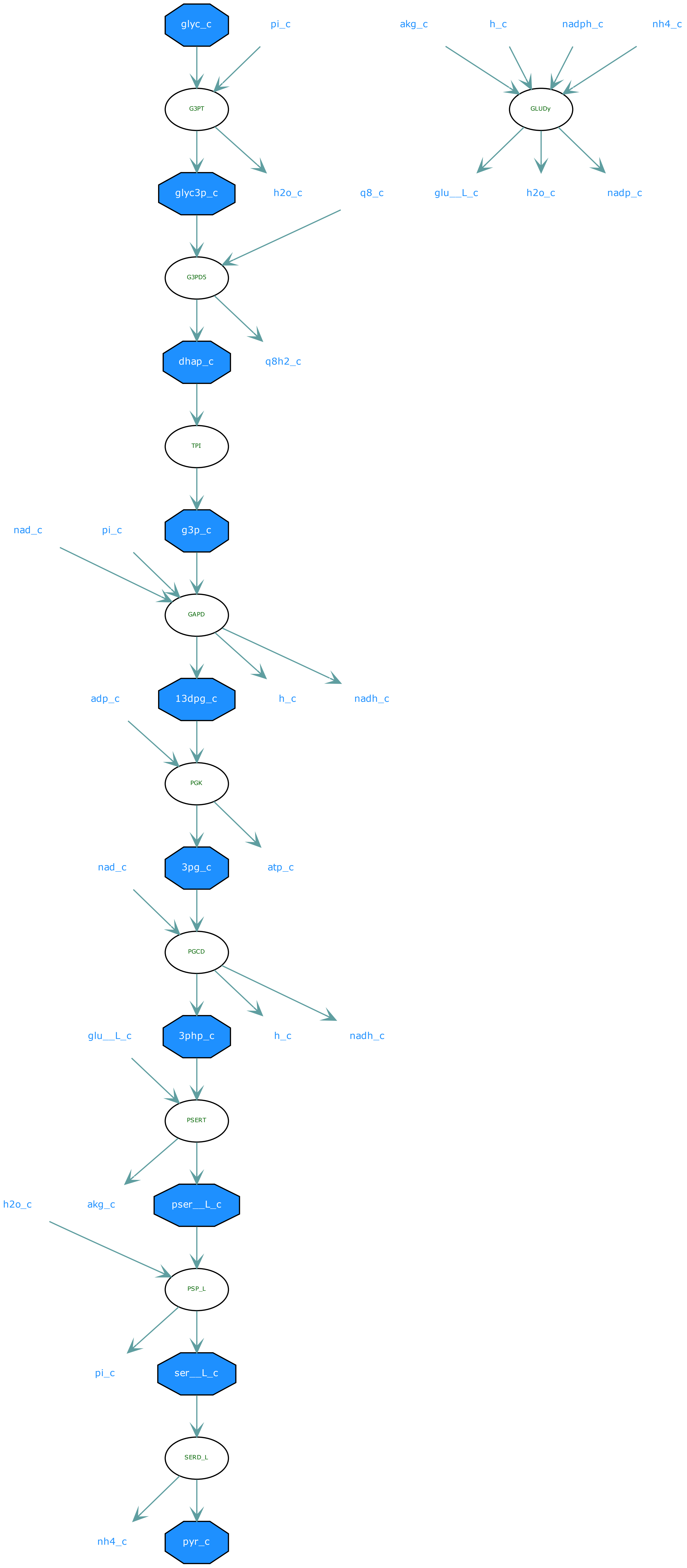

glyc_c
pi_c
akg_c
h_c
nadph_c
nh4_c
G3PT
GLUDy
glyc3p_c
h2o_c
q8_c
glu__L_c
h2o_c
nadp_c
G3PD5
dhap_c
q8h2_c
TPI
nad_c
pi_c
g3p_c
GAPD
adp_c
13dpg_c
h_c
nadh_c
PGK
nad_c
3pg_c
atp_c
PGCD
glu__L_c
3php_c
h_c
nadh_c
PSERT
h2o_c
akg_c
pser__L_c
PSP_L
pi_c
ser__L_c
SERD_L
nh4_c
pyr_c

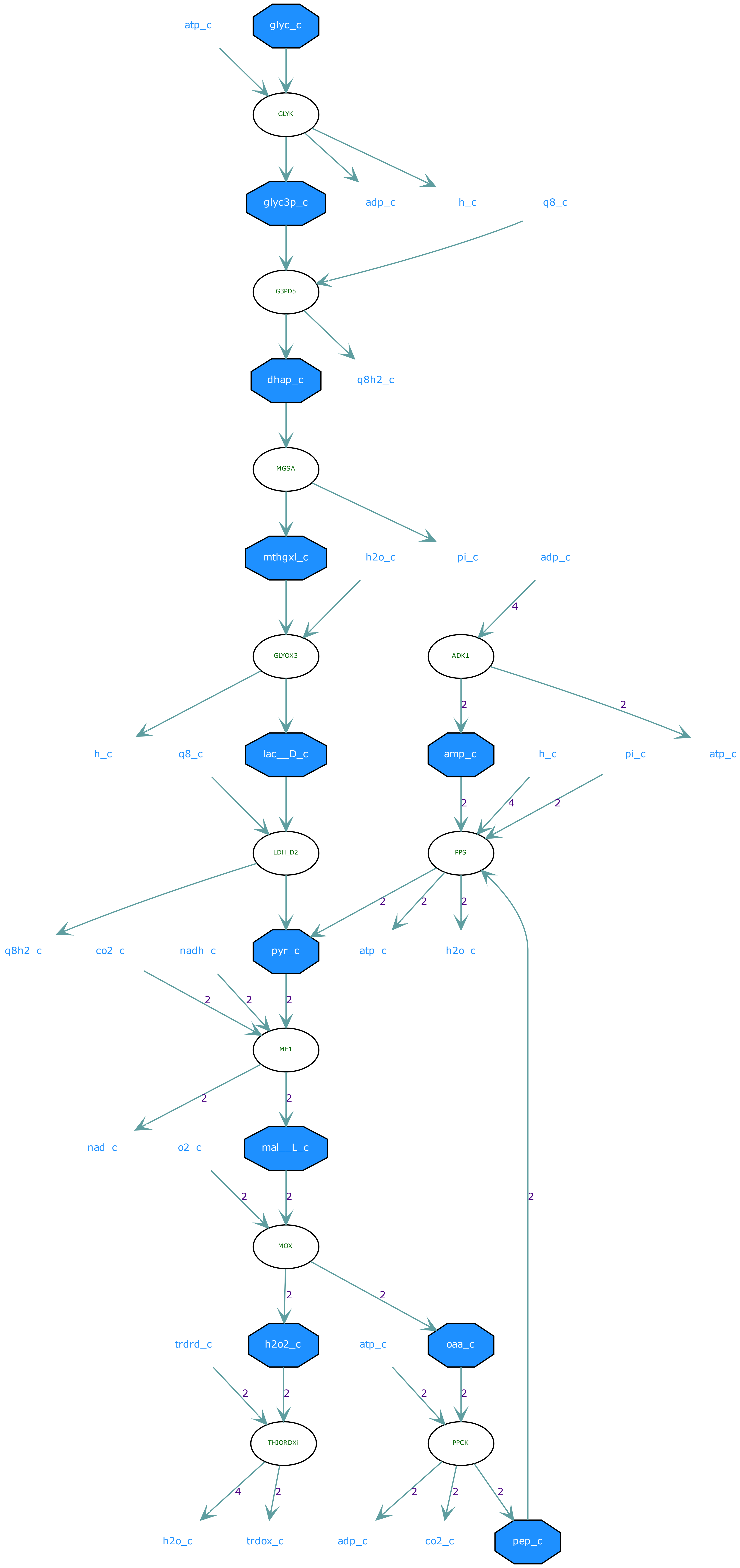

atp_c
glyc_c
GLYK
glyc3p_c
adp_c
h_c
q8_c
G3PD5
dhap_c
q8h2_c
MGSA
mthgxl_c
h2o_c
pi_c
adp_c
GLYOX3
ADK1
h_c
q8_c
lac__D_c
amp_c
h_c
pi_c
atp_c
LDH_D2
PPS
q8h2_c
co2_c
nadh_c
pyr_c
atp_c
h2o_c
ME1
nad_c
o2_c
mal__L_c
MOX
trdrd_c
h2o2_c
atp_c
oaa_c
THIORDXi
PPCK
h2o_c
trdox_c
adp_c
co2_c
pep_c

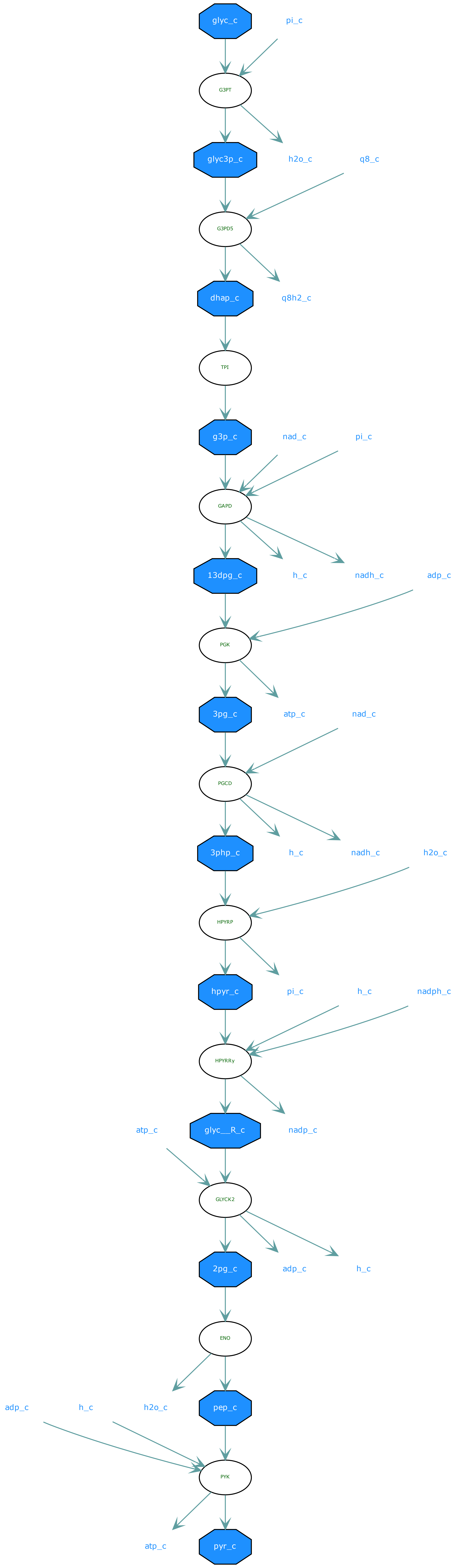

glyc_c
pi_c
G3PT
glyc3p_c
h2o_c
q8_c
G3PD5
dhap_c
q8h2_c
TPI
g3p_c
nad_c
pi_c
GAPD
13dpg_c
h_c
nadh_c
adp_c
PGK
3pg_c
atp_c
nad_c
PGCD
3php_c
h_c
nadh_c
h2o_c
HPYRP
hpyr_c
pi_c
h_c
nadph_c
HPYRRy
atp_c
glyc__R_c
nadp_c
GLYCK2
2pg_c
adp_c
h_c
ENO
adp_c
h_c
h2o_c
pep_c
PYK
atp_c
pyr_c

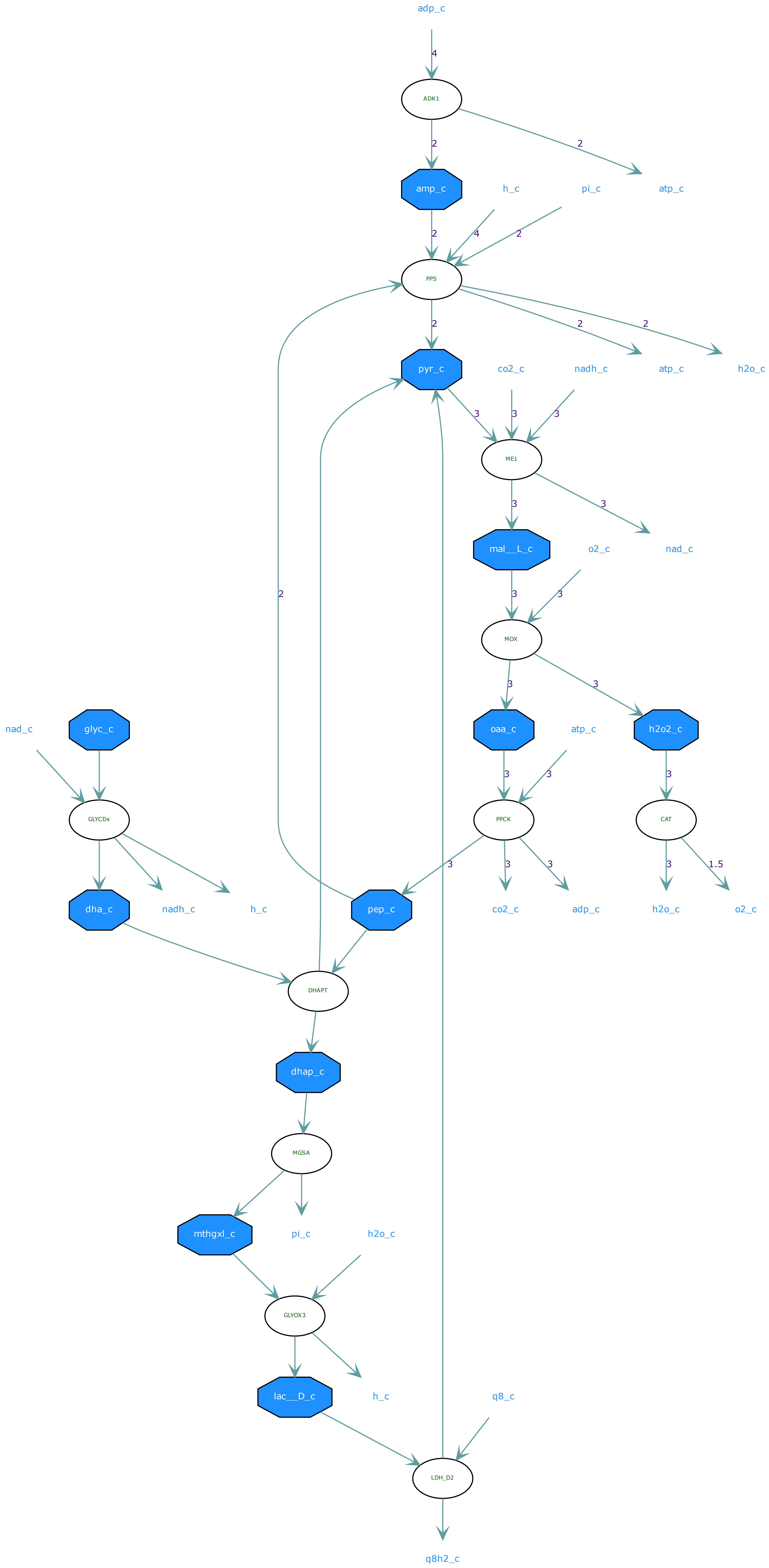

adp_c
4
ADK1
2
2
amp_c
h_c
pi_c
atp_c
2
4
2
PPS
2
2
2
pyr_c
co2_c
nadh_c
atp_c
h2o_c
3
3
3
ME1
3
3
mal__L_c
o2_c
nad_c
2
3
3
MOX
3
3
nad_c
glyc_c
oaa_c
atp_c
h2o2_c
3
3
3
GLYCDx
PPCK
CAT
3
3
3
3
1.5
dha_c
nadh_c
h_c
pep_c
co2_c
adp_c
h2o_c
o2_c
DHAPT
dhap_c
MGSA
mthgxl_c
pi_c
h2o_c
GLYOX3
lac__D_c
h_c
q8_c
LDH_D2
q8h2_c

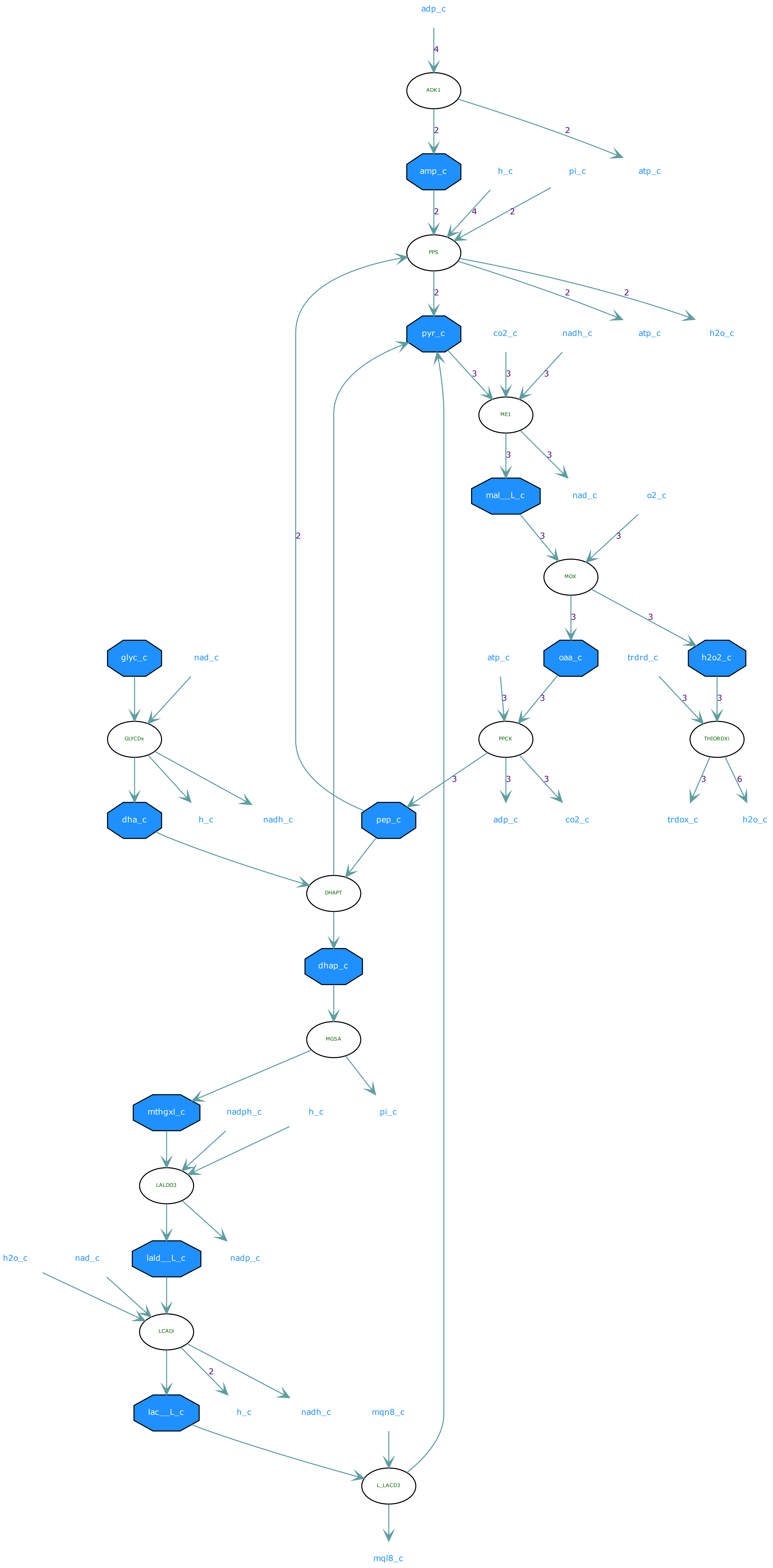
adp_c
ADK1
amp_c
h_c
pi_c
atp_c
PPS
pyr_c
co2_c
nadh_c
atp_c
h2o_c
ME1
mal__L_c
nad_c
o2_c
MOX
glyc_c
nad_c
atp_c
oaa_c
trdrd_c
h2o2_c
GLYCDx
PPCK
THIORDXi
dha_c
h_c
nadh_c
pep_c
adp_c
co2_c
trdox_c
h2o_c
DHAPT
dhap_c
MGSA
mthgxl_c
nadph_c
h_c
pi_c
LALDO3
h2o_c
nad_c
lald__L_c
nadp_c
LCADi
lac__L_c
h_c
nadh_c
mqn8_c
L_LACD3
mql8_c

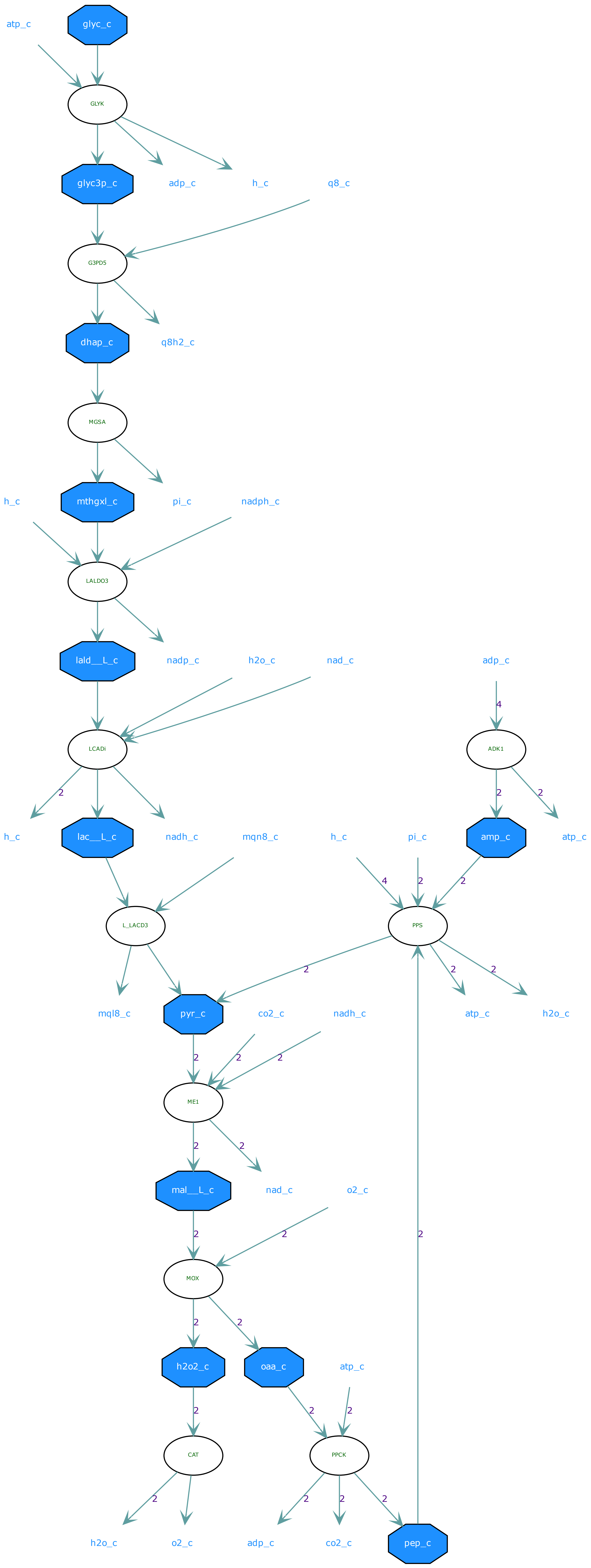
atp_c
glyc_c
GLYK
glyc3p_c
adp_c
h_c
q8_c
G3PD5
dhap_c
q8h2_c
MGSA
h_c
mthgxl_c
pi_c
nadph_c
LALDO3
lald__L_c
nadp_c
h2o_c
nad_c
adp_c
LCADi
ADK1
h_c
lac__L_c
nadh_c
mqn8_c
h_c
pi_c
amp_c
atp_c
L_LACD3
PPS
mql8_c
pyr_c
co2_c
nadh_c
atp_c
h2o_c
ME1
mal__L_c
nad_c
o2_c
MOX
h2o2_c
oaa_c
atp_c
CAT
PPCK
h2o_c
o2_c
adp_c
co2_c
pep_c

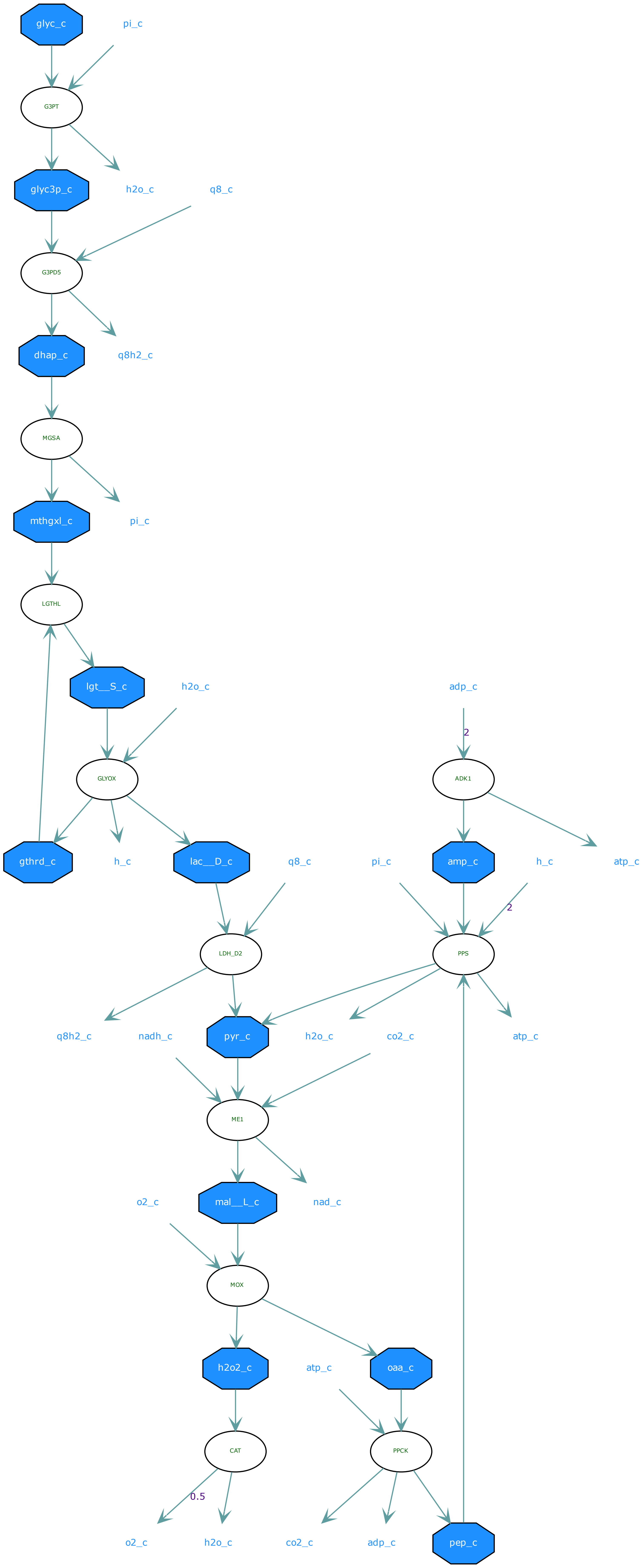
glyc_c
pi_c
G3PT
glyc3p_c
h2o_c
q8_c
G3PD5
dhap_c
q8h2_c
MGSA
mthgxl_c
pi_c
LGTHL
lgt__S_c
h2o_c
adp_c
2
GLYOX
ADK1
gthrd_c
h_c
lac__D_c
q8_c
pi_c
amp_c
h_c
atp_c
2
LDH_D2
PPS
q8h2_c
nadh_c
pyr_c
h2o_c
co2_c
atp_c
ME1
o2_c
mal__L_c
nad_c
MOX
h2o2_c
atp_c
oaa_c
CAT
PPCK
0.5
o2_c
h2o_c
co2_c
adp_c
pep_c

**Supplementary Figure S1**: *In silico* identified thermodynamically feasible glycolytic routes identified which could potentially convert glycerol to pyruvate. For the identification the latest metabolic model of *E. coli* from the BiGG database (1) was used.

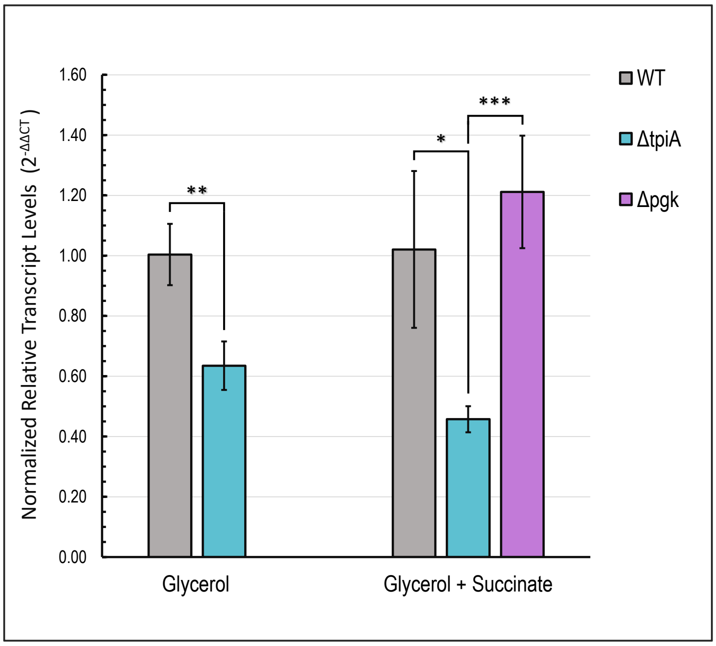


**Supplementary Figure S2**: Relative transcript levels of *mgsA*. Cells were harvested in exponential phase (OD600 0.5-0.6) from M9 minimal medium cultures. For comparison of WT to Δ*tpi* 20 mM glycerol was used as a carbon source, for comparison of WT, Δ*tpi* and Δ*pgk* 4 mM glycerol and 40 mM succinate were used as a carbon source. mRNA levels of *mgsA* was determined by RT-qPCR as described in the methods section and normalized to transcript level of 16S rRNA (*rrsA*). Data significance: *, $p < 0.05$, **, $p < 0.01$, ***, $p < 0.005$.

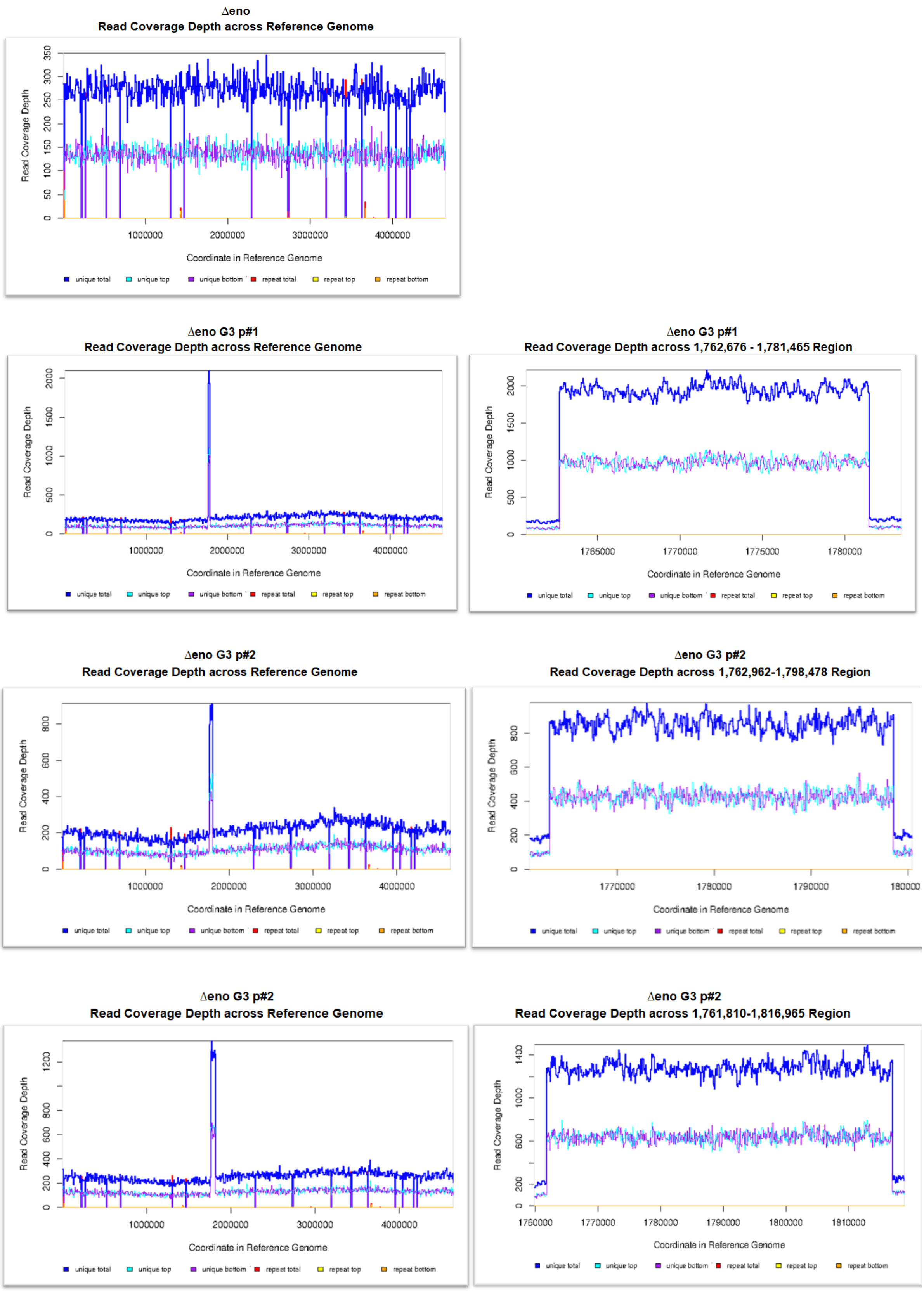


**Supplementary Figure S3**: **Genome sequencing coverage in the serine tolerant strain as analyzed by Breseq.** After sequencing the genomes of the three isolated strains (Δ*eno* G3 #1,2,3) and their ancestor strain (Δeno), the results were mapped against the *E. coli* MG1655 reference genome (GenBank accession no. U000913.3). The graphs indicate the coverage of the reads against the genome position for the whole length (left) and the read increased area (right).

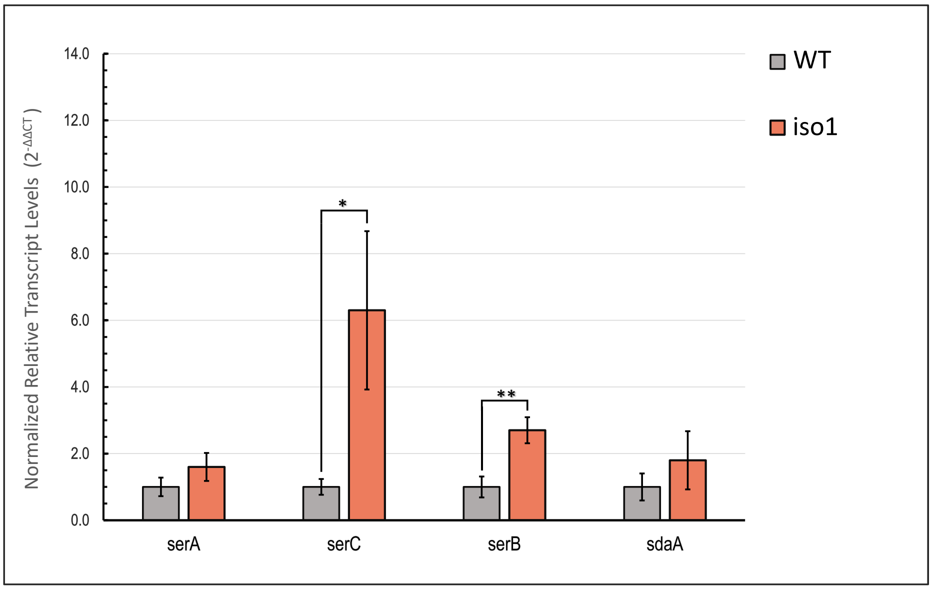


**Supplementary Figure S4**: Relative transcript levels of *serA, serB, serC* and *sdaA* in iso1 and WT strains. Cells were harvested in exponential phase ($OD_{600}$ 0.5-0.6) after growth on M9 minimal medium containing 20 mM glycerol. mRNA levels of *serA, serB, serC* and *sdaA* were determined by RT-qPCR as described in the methods section and normalized to transcript level of 16S rRNA (*rrsA*). Data significance: *, $p < 0.05$, **, $p < 0.005$.

**Supplementary Table S1: Mutations identified in the serine tolerant Δ*eno* strain.** The genomes of three isolates growing with increased serine concentration (Δ*eno* G3 #1,2,3) and their ancestor strain (Δ*eno*) were sequenced. X indicates the presence of the mutation (compared to the *E. coli* MG1655 reference genome (GenBank accession no. U000913.3) in the respective sequencing result.

### Predicted mutations

| position | mutation | annotation | gene | Δeno G3 #1 | Δeno G3 #2 | Δeno G3 #3 | Δeno |
|---|---|---|---|---|---|---|---|
| 81,261 | A→T | L233Q (CTG→CAG) | *leuB* ← | | | | X |
| 158,823 | Δ4 bp | coding (301-304/1398 nt) | *pcnB* ← | X | | | |
| 158,845 | (AGGCGGCAGTTACGGAACAGT)$_{1\rightarrow 2}$ | coding (282/1398 nt) | *pcnB* ← | | | | X |
| 158,883 | C→A | D82Y (GAC→TAC) | *pcnB* ← | | X | | |
| 257,908 | Δ776 bp | | *[crl]* | X | X | X | X |
| 1,405,877 | T→C | intergenic (+228/-102) | *abgR* → / → *smrA* | X | X | X | X |
| 1,978,503 | Δ776 bp | | *insB1–insA* | X | X | X | X |
| 1,979,486 | IS*5* (+) +4 bp | intergenic (-271/-264) | *insA* ← / → *uspC* | X | X | X | X |
| 2,173,365 | Δ2 bp | pseudogene (913-914/914 nt) | *gatC* ← | X | X | X | X |
| 3,560,455 | +G | intergenic (-2/+1) | *glpR* ← / ← *glpR* | X | X | X | X |
| 4,296,381 | +GC | intergenic (+587/+55) | *gltP* → / ← *yjcO* | X | X | X | X |

### Unassigned new junction evidence

| position | | | gene | | | | |
|---|---|---|---|---|---|---|---|
| = 70387 | | | *araC* | X | X | X | X |
| 4171751 = | | | *rrfB* | | | | |
| 1207790 = | | coding (290/630 nt) | *stfP* | | X | | |
| 1209619 = | | pseudogene (1/501 nt) | *stfE* | | | | |
| = 1299498 | | coding (305/630 nt) | *stfP* | | X | | |
| 1300698 = | | pseudogene (18/501 nt) | *stfE* | | | | |
| = 1299498 | | intergenic (+253/-1684) | *ychE/oppA* | X | X | | |
| 1300698 = | | intergenic (+1453/-484) | *ychE/oppA* | | | | |
| 1761810 = | | coding (703/747 nt) | *sufC* | | | 1061 (4.000) | |
| = 1816965 | | coding (171/750 nt) | *chbG* | | | | |
| 1762676 = | | coding (1334/1488 nt) | *sufB* | 680 (3.340) | | | |
| = 1781465 | | coding (71/1290 nt) | *ydiS* | | | | |
| 1762962 = | | coding (1048/1488 nt) | *sufB* | | 1718 (8.160) | | |
| = 1798478 | | coding (465/984 nt) | *pheS* | | | | |
| 3911839 = | | coding (1830/1830 nt) | *glmS* | X | X | X | |
| = 4171849 | | intergenic (+93/-208) | *rrfB/murB* | | | | |

### Supplementary Table S2: Identified mutations in the evolved Δ*eno* strains.

**NGR1**

| **chromosomal position*** | **mutation type** | **mutation event** | ***mutated gene*** | **intragenic position** | **amino acid change** | **intergenic mutation (distance to the flanking genes)** | **amplified / deleted region** | **mutations identified in the indicated strain (•)** | |
|---|---|---|---|---|---|---|---|---|---|
| | | | | | | | | **G5181** | **G5182** |
| 4114812 | A/T | SNP | *glpK* | 434 | I145N | | | • | • |
| 2098629 | A/T | SNP | *gnd* | 664 | W222R | | | • | • |
| 1913509 | A/T | SNP | *proQ* | 50 | L17Q | | | • | • |
| 3813831 | A/C | SNP | | | | *pyrE/rph(40/55)* | | • | • |
| 4183596-604 | | DEL 9 bp | *rpoC* | 224-232 | | | | • | • |
| 3055318 | G/T | SNP | *serA* | 1115 | T372N | | | • | • |
| | | | | | | | DEL(*gntU-yhhY*) | • | • |
| | | | | | | | DEL(*cpxP*) | • | • |
| | | | | | | | x2 *(zur-insA*) | • | • |

**NGR2**

| **chromosomal position*** | **mutation type** | **mutation event** | ***mutated gene*** | **intragenic position** | **amino acid change** | **intergenic mutation (distance to the flanking genes)** | **amplified / deleted region** | **mutations identified in the indicated strain (•)** | |
|---|---|---|---|---|---|---|---|---|---|
| | | | | | | | | **G5196** | **G5197** |
| 2973447 | C/T | SNP | *aas* | 590 | R197Q | | | • | • |
| 2058550 | G/A | SNP | *cbl* | 389 | T130M | | | • | |
| 2291468 | T/C | SNP | *ccmF* | 1459 | M487V | | | • | • |
| 2534052 | G/A | SNP | *crr* | 197 | G66D | | | • | • |
| 2872235 | G/A | SNP | *cysN* | 1207 | P403S | | | • | |
| 1250341 | C/T | SNP | *dhaR* | 53 | T18I | | | • | |
| 3533277 | A/T | SNP | *envZ* | 614 | L205Q | | | • | • |
| 2460903 | -/C | INS | | | | *fadL/yfdF* (235/131) | | • | |
| 1140280 | T/- | DEL | | | | *flgL/rne* (71/125) | | • | • |
| 1975804 | A/T | SNP | *flhC* | 65 | L22Q | | | • | • |
| 379237 | G/- | DEL | | | | *frmR/frmRAB (132/56)* | | • | • |
| 2940307 | C/- | DEL | *gcvA* | 283 | | | | • | • |
| 3048203 | C/T | SNP | *gcvT* | 487 | A163T | | | • | |
| 4115926 | G/T | SNP | *glpF* | 188 | S63Y | | | • | • |
| 3090583 | -/G | INS | *gshB* | 684 | | | | • | • |
| 2600510 | T/C | SNP | *hyfB* | 671 | M224T | | | • | |
| 2601551 | G/A | SNP | *hyfB* | 1712 | G571D | | | • | • |
| 722302 | A/G | SNP | *kdpD* | 1336 | W446R | | | • | • |
| 3476916 | C/T | SNP | *kefB* | 1714 | A572T | | | • | |
| 673763 | A/G | SNP | *leuS* | 244 | F82L | | | • | • |
| 1937431 | G/A | SNP | *lpxM* | 787 | R263C | | | | • |
| 2155124 | G/A | SNP | *mdtB* | 1838 | G613D | | | • | • |
| 2157817 | G/A | SNP | *mdtC* | 1408 | A470T | | | | • |
| 819024 | A/G | SNP | | | | *moaE/ybhL* (54/83) | | • | • |
| 979114 | C/T | SNP | *mukB* | 3566 | A1189V | | | • | • |
| 2857276 | C/A | SNP | *mutS* | 2162 | A721E | | | • | • |
| 1283205 | C/T | SNP | *narH* | 379 | R127C | | | • | • |
| 1534614 | -/A | INS | *narV* | 28 | | | | | • |
| 1913362 | G/T | SNP | *proQ* | 197 | S66* | | | • | • |
| 1368060 | T/- | DEL | | | | *pspE/ycjM* ((33/180) | | • | • |
| 3928170 | A/T | SNP | *ravA* | 947 | L316H | | | • | • |
| 1413508 | C/T | SNP | *recE* | 1903 | A635T | | | | • |
| 4180066 | C/T | SNP | *rpoB* | 799 | R267C | | | • | • |
| 1708175 | T/C | SNP | *rsxD* | 1010 | L337P | | | • | • |
| 3055325 | G/T | SNP | *serA* | 1108 | L370M | | | • | • |
| 560505 | C/T | SNP | *sfmD* | 1586 | S529L | | | | • |
| 3105232 | G/A | SNP | *speC* | 1946 | A649V | | | • | • |
| 3259260 | T/C | SNP | *tdcE* | 1181 | Q394R | | | • | • |
| 3078126 | T/C | SNP | *tktA* | 1532 | D511G | | | • | |
| 3177089 | G/A | SNP | *tolC* | 953 | G318E | | | • | • |
| 1329648 | G/A | SNP | *topA* | 577 | A193T | | | | • |
| 1331254 | A/G | SNP | *topA* | 2183 | Y728C | | | • | |
| 1058654 | -/G | INS | *torA* | 176 | | | | • | |

| | | | | | | | | | |
|---|---|---|---|---|---|---|---|---|---|
| 1491778 | A/T | SNP | *trg* | 1285 | I429F | | | | • |
| 3606830 | C/T | SNP | *tusA* | 190 | V64I | | | • | |
| 131055 | A/G | SNP | *yacH* | 206 | V69A | | | • | |
| 248170 | T/C | SNP | | | | *yafM/fhiA* (36/188) | | • | • |
| 1259939 | T/C | SNP | *ychM* | 88 | T30A | | | • | |
| 1492848 | T/C | SNP | *ydcI* | 248 | D83G | | | • | • |
| 1630183 | T/C | SNP | *ydfJ* | 127 | I43V | | | • | • |
| 1749334 | T/C | SNP | *ydhW* | 415 | T139A | | | • | |
| 1786127 | A/G | SNP | *ydiA* | 659 | E220G | | | • | • |
| 1766716 | G/A | SNP | | | | *ydiJ/ydiK (7/382)* | | • | • |
| 1869851 | -/G | INS | *yeaI* | 1443 | | | | • | • |
| 1882983 | G/A | SNP | *yeaW* | 295 | G99S | | | • | • |
| 2027827 | G/A | SNP | *yedA* | 265 | A89T | | | | • |
| 2203311 | G/A | SNP | *yehL* | 694 | A232T | | | | • |
| 2515043 | G/A | SNP | *yfeA* | 812 | T271M | | | • | • |
| 2628977 | A/G | SNP | | | | *yfgI/guaA* (90/3) | | | • |
| 2646876 | G/A | SNP | *yfhM* | 3434 | A1145V | | | • | |
| 2767665 | -/C | INS | | | | *yfjQ/yfjR* (157/60) | | • | • |
| 2993349 | G/A | SNP | *ygeM* | 419 | P140L | | | | • |
| 3064285 | G/A | SNP | *ygfH* | 1462 | G488S | | | • | • |
| 3351624 | T/C | SNP | *yhcC* | 449 | H150R | | | | • |
| 3637895 | T/- | DEL | | | | *yhiO/uspA* (152/239) | | • | • |
| 4198220 | T/C | SNP | | | | *yjaG/hupA* (103/84) | | • | • |
| 868537 | G/A | SNP | *yliA* | 1762 | A588T | | | • | • |
| 871599 | G/A | SNP | *yliD* | 487 | G163S | | | • | • |
| 1899328 | -/C | INS | *yoaE* | 282 | | | | • | • |
| 2778387 | C/T | SNP | *ypjA* | 2362 | D788N | | | • | • |
| 2988177 | -/T | INS | *yqeK* | 206 | | | | • | • |
| | | | | | | | x2(*rrfE-insL*) | • | • |
| | | | | | | | DEL(*insH-2 - insL-2*) | • | • |
| | | | | | | | DEL(*glpF-rraA*) | • | • |

**Supplementary Table S3:** Oligonucleotide primers used in this study.

| name | sequence | use |
|---|---|---|
| serA_FBR_A_F | CTTTAAAGTTAAGAGGCAAGAATGCATCATCACCATCACCACGCAAAGGTATCGCTGGAGAAAGACAAGATTAAG | Cloning of feedback resistant serA |
| serA_FBR_B | GAAGGACGTTGTCAAATTCACACAGCGG | Cloning of feedback resistant serA |
| serA_FBR_C | CCGCTGTGTGAATTTGACAACGTCCTTC | Cloning of feedback resistant serA |
| serA_FBR_D | GCGCGGCGATCGCGACGCCCTGCTCGGCGAAGATTTTGTTCAGCGCAGTTAGCACGCCCGGACGCGCTTCCGCGATGTGCATC | Cloning of feedback resistant serA |
| serA_FBR_E | GATGCACATCGCGGAAGCGCGTCCGGGCGTGCTAACTGCGCTGAACAAAATCTTCGCCGAGCAGGGCGTCGCGATCGCCGCGC | Cloning of feedback resistant serA |
| serA_FBR_F_R | CTCTTACGTGCCCGATCAACGCTAGCTTAGTACAGCAGACGGGCGCGAATGGTACCCGGAATAGCTTTCATTGCTTGCAGCGCTTTTTC | Cloning of feedback resistant serA |
| serB_A_F | CTTTAAAGTTAAGAGGCAAGAATGCATCATCACCATCACCACCCTAACATTACCTGGTGCGACCTGCC | Cloning of serB |
| serB_B_R | CTCTTACGTGCCCGATCAACGCTAGCTTACTTCTGATTCAGGCTGCCTGAGAGGATG | Cloning of serB |
| serC_A_F | CTTTAAAGTTAAGAGGCAAGAATGCATCATCACCATCACCACGCTCAAATCTTCAATTTTAGTTCTGGTCCGGCAATG | Cloning of serC |
| serC_B | GGTTTCATTCGGGCAATAGTGCATATAAGCAGCATTATC | Cloning of serC |
| serC_C | GATAATGCTGCTTATATGCACTATTGCCCGAATGAAACC | Cloning of serC |
| serC_D_R | CTCTTACGTGCCCGATCAACGCTAGCTTAACCGTGACGGCGTTCGAACTCAACC | Cloning of serC |
| sdaA_F | ATGCATCATCACCATCACCACATTAGTCTATTCGACATGTTTAAGGTGG | Cloning of sdaA |
| sdaA_R | CTCTTACGTGCCCGATCAACGCTAGCTTAGTCACACTGGACTTTGATTGCC | Cloning of sdaA |
| mgsA_A_F | CTTTAAAGTTAAGAGGCAAGAATGCATCATCACCATCACCACGAACTGACGACTCGCACTTTACCTGCG | Cloning of mgsA |
| mgsA_B_R | CTCTTACGTGCCCGATCAACGCTAGCTTACTTCAGACGGTCCGCGAGATAACGC | Cloning of mgsA |
| serA-pet-F | GGCCATATCGAAGGTCGTCATATGGCAAAGGTATCGCTGGAGAAAGACAAGATTAAG | Cloning of serA variants into pet16B |
| serA-pet-R | CTTTGTTAGCAGCCGGATCCTCGAGTTAGTACAGCAGACGGGCGCGAATGG | Cloning of serA variants into pet16B |
| serA_KO_F | CCTGCCCGTTGATTTTCAGAGAAGGGGAATTAGTACAGCAGACGGGCGCGAATTAACCCTCACTAAAGGGCG | serA KO-cassette amp. Gene Bridges, Km |
| serA_KO_R | GCGGATGCAAATCCGCACACAACATTTCAAAAGACAGGATTGGGTAAATGTAATACGACTCACTATAGGGCTC | serA KO-cassette amp. Gene Bridges, Km |
| sdaA_KO_F | TATTAGTTCGTTACTGGAAGTCCAGTCACCTTGTCAGGAGTATTATCGTGAATTAACCCTCACTAAAGGGCG | *sdaA* KO-cassette amp. Gene Bridges, Km |
| sdaA_KO_R | ATCCGTTGCAGATGGGCGAGTAAGAAGTATTAGTCACACTGGACTTTGATTAATACGACTCACTATAGGGCTC | *sdaA* KO-cassette amp. Gene Bridges, Km |
| sdaB_KO_F | GCCGCTTTCGGGCGGCGCTTCCTCCGTTTTAACGCGATGTATTTCCTATGAATTAACCCTCACTAAAGGGCG | *sdaB* KO-cassette amp. Gene Bridges, Km |
| sdaB_KO_R | CCTCGCAAAACGAGGCCTTTGGAGAGCGATTAATCGCAGGCAACGATCTTTAATACGACTCACTATAGGGCTC | *sdaB* KO-cassette amp. Gene Bridges, Km |
| tdcB_KO_F | GTACTCATATCCTATCCTCAACGAATTAATTAAGCGTCAACGAAACCGGTAATTAACCCTCACTAAAGGGCG | *tdcB* KO-cassette amp. Gene Bridges, Km |
| tdcB_KO_R | TCGGTTACGGTTACCTACATATTTAATTCAGGCGAAGAGGTTTTATAATGTAATACGACTCACTATAGGGCTC | *tdcB* KO-cassette amp. Gene Bridges, Km |
| tdcG_KO_F | GCACCCAAGGATGAAAGCTGACAGCAATGTCAGCCGCAGACCACTTTAATAATTAACCCTCACTAAAGGGCG | *tdcG* KO-cassette amp. Gene Bridges, Km |
| tdcG_KO_R | AGGTCGTTCCGCTCCACTTCACTGAACGGCAATCCGAGGGTGTGGATATGTAATACGACTCACTATAGGGCTC | *tdcG* KO-cassette amp. Gene Bridges, Km |
| mgsA_KO_K_F | AACAGGTGGCGTTTGCCACCTGTGCAATATTACTTCAGACGGTCCGCGAGAATTAACCCTCACTAAAGGGCG | *mgsA* KO-cassette amp. Gene Bridges, Km |
| mgsA_KO_K_R | TAAGTGCTTACAGTAATCTGTAGGAAAGTTAACTACGGATGTACATTATGTAATACGACTCACTATAGGGCTC | *mgsA* KO-cassette amp. Gene Bridges, Km |
| dld_KO_F | CGCTATTCTAGTTTGTGATATTTTTTCGCCACCACAAGGAGTGGAAAATGGTGTAGGCTGGAGCTGCTTC | *dld* KO-cassette amp. pKD3/4, CAP/Km |

| | | |
|---|---|---|
| dld_KO_R | GGATGGCGATACTCTGCCATCCGTAATTTTTACTCC<br>ACTTCCTGCCAGTTCATATGAATATCCTCCTTAG | *dld* KO-cassette amp. pKD3/4, CAP/Km |
| gldA_KO_F | CGACTGGAATGCCGCATTTGGCACTACTCATCTCTA<br>AAGGAGCAATTATGGTGTAGGCTGGAGCTGCTTC | *gldA* KO-cassette amp. pKD3/4, CAP/Km |
| gldA_KO_R | TCCCGGACAAGCCGGGAGTTTGGAGTAGGTTATTC<br>CCACTCTTGCAGGAACATATGAATATCCTCCTTAG | *gldA* KO-cassette amp. pKD3/4, CAP/Km |
| gloA_KO_F | TACTAAAACAACATTTTGAATCTGTTAGCCATTTTGA<br>GGATAAAAAGATGGTGTAGGCTGGAGCTGCTTC | *gloA* KO-cassette amp. pKD3/4, CAP/Km |
| gloA_KO_R | GGCGCGATGAGTTCACGCCCGGCAGGAGATTAGTT<br>GCCCAGACCGCGACCCATATGAATATCCTCCTTAG | *gloA* KO-cassette amp. pKD3/4, CAP/Km |
| aldA_KO_F | AACAATGTATTCACCGAAAACAAACATATAAATCACA<br>GGAGTCGCCCATGGTGTAGGCTGGAGCTGCTTC | *aldA* KO-cassette amp. pKD3/4, CAP/Km |
| aldA_KO_R | GAGGAAAAAACCTCCGCCTCTTTCACTCATTAAGAC<br>TGTAAATAAACCACCATATGAATATCCTCCTTAG | *aldA* KO-cassette amp. pKD3/4, CAP/Km |
| hchA_KO_C_F | CGCAAATATAGTGACTACCCTAACTAAGCAACAATAA<br>GGAATACACTATGGTGTAGGCTGGAGCTGCTTC | *hchA* KO-cassette amp. pKD3/4, CAP/Km |
| hchA_KO_C_R | TATGCGCTTACATTCAAACGTAACAGGGATTAACCC<br>GCGTAAGCTGCCAGCATATGAATATCCTCCTTAG | *hchA* KO-cassette amp. pKD3/4, CAP/Km |
| ppsA_KO_C_F | CGGCGACTAAACGCCGCCGGGGATTTATTTTATTTC<br>TTCAGTTCAGCCAGGTGTAGGCTGGAGCTGCTTC | *ppsA* KO-cassette amp. pKD3/4, CAP/Km |
| ppsA_KO_C_R | AGAAATGTGTTTCTCAAACCGTTCATTTATCACAAAA<br>GGATTGTTCGATGCATATGAATATCCTCCTTAG | *ppsA* KO-cassette amp. pKD3/4, CAP/Km |
| serA_KO_Ver_F | CTCAACATCGCGACGCAAAC | PCR verification of serA deletion |
| serA_KO_Ver_R | TCTGGAGCAGACTCGCAAAG | PCR verification of serA deletion |
| tdcG_KO_Ver_F | CTTATTATTTTTTTCCGAGCCGCATCAAGGCGATATG | PCR verification of tdcG deletion |
| tdcG_KO_Ver_R | CGTGTTTATCACCGATCTGAATGATTTTGCCAC | PCR verification of tdcG deletion |
| tdcB_KO_Ver_F | CGATTGCCGTACCAAACAAGCC | PCR verification of tdcB deletion |
| tdcB_KO_Ver_R | AGCAGCATCGGTTTTGGTGGAA | PCR verification of tdcB deletion |
| sdaB_KO_Ver_F | GGGTCTGATTGCAATCTCCGCA | PCR verification of sdaB deletion |
| sdaB_KO_Ver_R | ATGAACAGCCACGATAACCCCC | PCR verification of sdaB deletion |
| sdaA_KO_Ver_F | GGCGCTGCAAATTGGTGTGAAA | PCR verification of sdaA deletion |
| sdaA_KO_Ver_R | CCTGACGCAACAGTGGAAGTGT | PCR verification of sdaA deletion |
| mgsA_KO_Ver_F | CACCGCAGTCTCAGGTGCTCAC | PCR verification of mgsA deletion |
| mgsA_KO_Ver_R | CTGACCCGGGCACGCCATCG | PCR verification of mgsA deletion |
| dld_KO_Ver_F | TTCTTCCTTTGTTGCCCGACGT | PCR verification of dld deletion |
| dld_KO_Ver_R | TAGTGATGGACGCGTTTGGCAA | PCR verification of dld deletion |
| gldA_KO_Ver_F | CGGCCTACAAAAGCACGCAAAT | PCR verification of gldA deletion |
| gldA_KO__Ver_R | CACCCTGCCCTTAGATGTAGCG | PCR verification of gldA deletion |
| gloA_KO_Ver_F | GTAATCCAACATTGCGAGCGGC | PCR verification of gloA deletion |
| gloA_KO_Ver_R | TCCATTTTCAGGGTGATGGCGG | PCR verification of gloA deletion |
| aldA_KO_Ver_F | CCACTTGTTTGCAAACGGGCAT | PCR verification of aldA deletion |
| aldA_KO_Ver_R | GTTTGATGCCACGCAAACGGAA | PCR verification of aldA deletion |
| hchA_KO_Ver_F | CAGCACTAAATCTCTCCCCGCC | PCR verification of hchA deletion |
| hchA_KO_Ver_R | CGTAGGTCAGGGACTAGGCCTT | PCR verification of hchA deletion |
| ppsA_KO_Ver_F | TCTCTGCCGTATGGATGAGGCT | PCR verification of ppsA deletion |
| ppsA_KO_Ver_R | GCGTGTCCTAATACCTCCGCAG | PCR verification of ppsA deletion |
| SdaA-ProEx-F | CCGCAGGCATAATTCGTGAGCTGGCGCTGCAAATT<br>GGTGTGAAACCCTGAAATTAACCCTCACTAAAGGGC<br>GGAGCTGCTTCGAAGTTC | sdaA promoter exchange |
| CAP sdaA-R | TAATACGACTCACTATAGGGCTCCATATGAATATCCT<br>CCTTAG | sdaA promoter exchange |
| pS-bridge | GAGCCCTATAGTGAGTCGTATTAAATACTTGACATAT<br>CACTGTGATTCACATATAATATGCG | sdaA / serA promotor exchange |
| SdaA-ProEx-R | GAAGATGAGGGACCAATCCCCACCTTAAACATGTCG<br>AATAGACTAATCATTCTTGCCTCTTAACTTTAAAGTTA<br>AACAAAATTATTTCTATTAACTAGTGAATTC | sdaA promoter exchange |
| serA*-ProEx-F | AATTAACCCTCACTAAAGGGCGGAGCTGC | serA promoter exchange |
| CAP-SerA*-R | CTAAGGAGGATATTCATATGGAGCCCTATAGTGAGT<br>CGTATTA | serA promoter exchange |
| serA*-ProEx-R | TTAGTACAGCAGACGGGCGCGAATGG | serA promoter exchange |
| 2660_sdaA_Pro_V_F | GCCAGTGAAGATGAAGTCTC | PCR verification of sdaA promotor exchange |
| 2661_sdaA_Pro_V_R | ACAGTGAACCATAAACGTCC | PCR verification of sdaA promotor exchange |
| mgsA_qPCR_F | GCACACGATCACTGCAAAC | Amplifying 3' portion of mgsA for qPCR |
| mgsA_qPCR_R | CTTCTGAGATCAATGCGCC | Amplifying 3' portion of mgsA for qPCR |
| serA_qPCR_F | AAGAATCCATCCGCGATGCC | Amplifying 3' portion of serA for qPCR |
| serA_qPCR_R | ACAGAAACAGCCAATAGCGAC | Amplifying 3' portion of serA for qPCR |
| serB_qPCR_F | GACCCAATACCAGAGCAAAC | Amplifying 3' portion of serB for qPCR |
| serB_qPCR_R | ATCCATCACCAGCAAACCC | Amplifying 3' portion of serB for qPCR |
| serC_qPCR_F | TTAAACAGGCTCAACAGGAAC | Amplifying 3' portion of serC for qPCR |
| serC_qPCR_R | CGTGCAGTATTTTTTCGCTTC | Amplifying 3' portion of serC for qPCR |
| sdaA_qPCR_F | GTCCCTCATCTTCCCATACC | Amplifying 3' portion of sdaA for qPCR |
| sdaA_qPCR_R | AAATCCACTTCATGCCGTC | Amplifying 3' portion of sdaA for qPCR |
| rrsA_qPCR_F | CTCTTGCCATCGGATGTGCCCA | Amplifying 3' portion of rrsA for qPCR |
| rrsA_qPCR_R | CCAGTGTGGCTGGTCATCCTCTCA | Amplifying 3' portion of rrsA for qPCR |

**Supplementary Method: A constraint-based method for finding glycolysis bypasses**

We use an approach that is very similar to the one used recently for finding latent carbon fixation cycles in *E. coli*. We set up a Mixed Integer Linear Problem (MILP)-based optimization problem which simultaneously looks for solutions that balance an objective reaction (here, glycerol to pyruvate) while minimizing the number of reactions. In the section titled Alterations to the iML1515 model, we lay out the changes we made to the set of reactions in the iML1515 model. In the section titled Formulation of the Mixed Integer Linear Problem, we describe how the MILP problem is formulated. In the section titled Iterating the space of solutions, we describe how we use the MILP framework to cover a large part of the feasible solution space of glycolysis bypass pathways.

**Alterations to the iML1515 model**

We made a few changes to the genome-scale model of *E. coli* (iML1515) (1):

- Removing all non-cytoplasmic reactions (i.e. exchange or transport reactions), except for exchange reactions of inorganic metabolites: protons, water, orthophosphate, ammonium, and oxygen
- Removing all boundary reactions (i.e. sink reactions needed to allow certain co-factors to leave the system)
- Replacing all flavoredoxins and thioredoxins with NADP(H): We replaced all flavoredoxins and thioredoxins with NADP(H), since we do not have a good estimate of their reduction potential, and therefore the MDF for pathways using them was artificially high. We can assume that the electrons used for reducing CO2 in the carbon fixation cycle ultimately have to pass through NADPH, and therefore a simple solution was to replace the electron donor with NADPH. This way, we could keep the flavoredoxins/thioredoxins-dependent reactions in the model while having a more realistic estimate of their thermodynamics.
- Removing the formate-tetrahydrofolate ligase reaction (FTHFLi): We found that the reaction formate-tetrahydrofolate ligase (FTHFLi) appears in some of the solutions although the gene associated with this reaction is unknown. FTHFLi was thus excluded from our model altogether. Notably, removing this reaction does not significantly affect the space of solutions, because it can be easily replaced by GAR transformylase-T (GART) and the reverse reaction of Phosphoribosylglycinamide formyltransferase (GARFT).
- Add the objective reaction (OBJ): glycerol + n ADP + n Pi → pyruvate + n ATP + n H2O where "n" refers to the ATP yield. In this study the allowed values were -1, 0, and 1.
- Setting the bounds (the range of possible fluxes) of all remaining reactions to be between -10 and 10

**Formulation of the Mixed Integer Linear Problem**

The mixed integer linear problem was formulated as follows:

Minimize

$$\Sigma_i \; z_i \qquad \textit{Eq. 1}$$

Such that

$$Sv = 0 \qquad \textit{Eq. 2}$$

$$v_{obj} = -1 \qquad \textit{Eq. 3}$$

$$v - \beta z \leq 0 \qquad \textit{Eq. 4}$$

$$0 \leq -g^o - S^T x + M(1-z) \qquad \textit{Eq. 5}$$

$$z \in \{0, 1\}^n \qquad \textit{Eq. 6}$$

$$0 \leq v \leq \beta \qquad \textit{Eq. 7}$$

$$ln(C_{min}) \leq x \leq 1n(C_{max}) \qquad \textit{Eq. 8}$$

where the vector $v$ contains the relative reaction rates, $z$ are the Boolean reaction indicators, $x$ are the log-scaled metabolite concentrations, and $g^0$ is a vector of all the reactions' standard Gibbs free energies in units of $RT$, i.e. $g_i^0 = \Delta_r G'^0(i) / RT$. $\beta$ is a parameter that limits the maximal rate for each single reaction in the pathway relative to the objective reaction, i.e. glycerol → pyruvate (with varying amounts of ATP production), and was set arbitrarily to 10. The rate of the objective reaction ($v_{OBJ}$) is set to be exactly -1 (Equation 3). This ensures that any pathway solution would exactly balance it, i.e. the overall reaction in the pathway would be glycerol → pyruvate. $M$ is a parameter which has a large value, much higher than any of the values in $g^0$. The lower and upper bounds on the concentrations of most metabolites were set to 1 μM and 10 mM. Only 14 central metabolites and co-factors were confined to more specific ranges based on physiological data (see Supplementary Table 4).

Note that as a pre-processing step, we split all reactions to a forward and backward reaction and therefore all (uni-directional) rates must be positive. Equation 4 ensures that each reaction indicator ($z_i$) can be equal to 0, only if the rate is 0. We don't need to care about it being equal to 1 even if a reaction is not active, since the optimization goal (which maximizes the sum of all indicators) will prevent that. Equation 5 ensures that every active reaction (i.e. with $z_i = 1$) has a positive driving force (which is given by $-g_i{}^0 - \Sigma_i S_{ij} x_j$).

#### Iterating the space of solutions

Our MILP objective function is to minimize the sum of all indicators (which is equal to the number of reactions in the pathway). In order to explore the space of possible glycolysis bypasses, we then iterate the solution space using integer-cuts to eliminate all the potential solutions within a radius of 2 around each one of the previously found solutions (using the same method as in (2)). We stopped the search after 10 solutions were found (for each possible ATP yield).

**Supplementary Table S4. The allowed concentration ranges for metabolites in the model.**

| Compound | BiGG identifier | Concentration range |
|---|---|---|
| ATP | atp_c | 5 mM |
| ADP | adp_c | 0.5 mM - 2.5 mM |
| AMP | amp_c | 0.5 mM - 2.5 mM |
| NAD+ | nad_c | 1 mM |
| NADH | nadh_c | 10 µM - 100 µM |
| NADP+ | nadp_c | 10 µM |
| NADPH | nadph_c | 10 µM - 100 µM |
| O2 | o2_c | 273 µM |
| CO2 | co2_c | 6.3 mM |
| CoA | coa_c | 1 mM - 5 mM |
| orthophosphate | pi_c | 1 mM - 10 mM |
| pyrophosphate | ppi_c | 0.5 mM - 1.5 mM |
| ammonia | nh4_c | 1 mM - 10 mM |
| alpha-ketoglutarate | akg_c | 0.5 mM - 5 mM |
| glutamate | glu__L_c | 30 mM - 150 mM |

Note: All metabolites that do not appear in this table were constrained by the default ranges of 1 µM to 10 mM.
